# The broad-spectrum RumC1 bacteriocin targets a transient peptidoglycan intermediate of the nascent cell wall

**DOI:** 10.64898/2026.03.19.712952

**Authors:** Anne Boyeldieu, Mathieu Bergé, Clarisse Roblin, Lama Shamseddine, Anna M. Diaz-Rovira, Anne-Lise Soulet, Christian Basset, Lauriane Plouhinec, Agnès Amouric, Salomé Milhavet, Léa Mélissa Perrault, Joana Marx Pereira da Cunha, Calum Johnston, Sylvie Kieffer-Jaquinod, Marc Maresca, Josette Perrier, Anne Chouquet, Cécile Morlot, Victor Guallar, Mickael Lafond, Victor Duarte, Nathalie Campo, Patrice Polard

**Affiliations:** University of Toulouse, Centre National de la Recherche Scientifique, Laboratoire de Microbiologie et Génétique Moléculaires (LMGM; UMR5100), Centre de Biologie Intégrative (CBI), 31062 Toulouse, France; Aix-Marseille Univ., CNRS, Centrale Med, iSm2, 13013 Marseille, France; Univ. Grenoble Alpes, CNRS, CEA, IRIG, Laboratoire Chimie et Biologie des Métaux (LCBM), 38054 Grenoble, France; Barcelona Supercomputing Center, Barcelona, Spain; INRAE, Aix-Marseille Univ., BBF, Biodiversité et Biotechnologie Fongiques, Marseille 13009, France; Univ. Grenoble Alpes, INSERM, CEA, IRIG, Biologie à Grande Echelle (BGE), 38054 Grenoble, France; Univ. Grenoble Alpes, CNRS, CEA, IBS, F-38000 Grenoble, France; Institució Catalana de Recerca i Estudis Avançats (ICREA), Barcelona, Spain

**Author notes:** Thornhill Therapeutics, 329 Oyster Point Boulevard, South San Francisco, CA 94080.

**Keywords:** Bacteriocin, Sactipeptide, Peptidoglycan, Cell wall synthesis, antibacterial mechanism of action, Bacteriocin immunity protein, Vancomycin, *Streptococcus pneumoniae*

## Abstract

RumC1 is a structurally unique bacteriocin with broad-spectrum efficacy, including against multidrug-resistant pathogens, yet acting by an undefined mechanism. By integrating genetics, biochemistry, computational modeling and single-cell fluorescence microscopy, we demonstrate that RumC1 is a distinct cell-wall-targeting toxin. First, all RumC1-resistant mutants isolated through a high-rate, genome-wide mutagenic screening exhibited specific impairments in peptidoglycan homeostasis regulation, pinpointing this pathway as critical for RumC1 activity. Second, RumC1 selectively accumulates within neosynthesized peptidoglycan, leading to cell growth arrest and death in a dose-dependent manner. Third, we characterize the RumI_c_1 immunity protein of the RumC1 biosynthetic cluster as a peptidase acting at the cell surface to protect the cells by trimming the stem peptide crucial for cell-wall assembly. As such, RumI_c_1 provides cross-protection against vancomycin, while RumC1 is demonstrated to act differently from this glycopeptide antibiotic. Collectively, these findings establish RumC1 as a toxin targeting a key peptidoglycan intermediate of cell wall maturation.

## Introduction

A central concern for global health is the growing difficulty of treating bacterial infections caused by species resistant to multiple drugs (MDR), including all conventional antibiotics^1,2^. The development of improved clinical versions of antibiotics has failed to provide a sustainable solution against these pathogens^3,4^. This limitation has intensified the search for new antibacterial agents, with a particular focus on antimicrobial peptides (AMPs) produced by microbiota from various ecological niches^5,6^. Innovative approaches have enabled the discovery and analysis of novel AMPs, either *via* their direct isolation from uncultivable microbial species or by identifying their biosynthetic gene clusters (BGCs) within large metagenomic databases, followed by heterologous expression and purification in laboratory strains^7,8^. Such advanced screens have uncovered several AMPs with unprecedented structural scaffolds and robust activity against prominent MDR pathogens, suggesting that these newly identified compounds may employ distinct mechanisms of action^9^. However, to exploit their potential as next-generation antibacterial agents, it is essential to identify their cellular target(s) and to evaluate how sensitive species might evolve resistance to their effects. Here, we followed these two interlinked aims for a bacteriocin of the human gut microbiota that belongs to this shortlist of promising AMPs acting *via* an undefined mode of action, the Ruminoccocin C1 (RumC1).

RumC1 originates from a complex chromosomally-encoded BGC in the commensal bacterium *Ruminococcus gnavus* E1^10,11^ (Fig. S1A). It belongs to the large family of ribosomally synthetized and post-translationally modified peptides (RiPPs)^12^. RumC1 is an atypical 44-amino-acids long sactipeptide, characterized by a compact double hairpin-like structure stabilized by four sulfur to α-carbon thioether bonds (Fig. S1B). These bonds are introduced by one of the two BGC-encoded sactisynthases (i.e., RumM_c_1 or RumM_c_2; Fig. S1A), and the peptide is subsequently exported as a pre-peptide by an ABC transporter encoded in the same BGC (RumT_c_)^11,13–15^. Notably, the final maturation of RumC1 is performed by the pancreatic trypsin of the human host^13^. This elaborate post-translational processing of RumC1 has been successfully recapitulated in *Escherichia coli*, allowing its purification as a homogeneous peptide species that has facilitated its comprehensive structural and functional characterization^13,14^. RumC1 exhibits remarkable antibacterial efficiency, demonstrating activity at micromolar and lower concentrations than conventional antibiotics, against monoderm species including many MDR pathogens, and LPS-poor or LPS-deficient diderms^13,14,16^. RumC1 is highly resistant to heat-shock, extreme pHs, and digestive proteases^14^. Importantly, RumC1 remains non-toxic to several human cell types, and is effective in clearing *Bacillus cereus* and *Clostridium perfringens* infections in simulated intestinal epithelium models and mice, respectively^14^.

Singularly, the *rumC* BGC encodes 4 additional paralogues of identical size (RumC2 to RumC5; Fig. S1A), which are expressed in *vivo*^11^. RumC5 exhibits an equivalent bactericidal activity as RumC1, while RumC3 and RumC4 are equally less efficient and RumC2 is surprisingly inactive^16^. A single amino acid substitution in RumC1 (the A12E mutation carried by RumC2) markedly reduces its antibacterial activity, while the converse mutation restores RumC2 activity^16^. Studies conducted on the RumC1-sensitive pathogen *Clostridium perfringens* showed that when applied several folds above the minimal inhibitory concentration (MIC), RumC1 impairs the synthesis of essential macromolecules, i.e. DNA, RNA, proteins and precursors of the peptidoglycan (PG), leading to a global metabolic collapse accompanied by gradual ATP depletion^14^. Unlike many bacteriocins, RumC1 achieves this without direct alteration of the cell membrane, as it does not form pores or induce depolarization^13^. Finally, repeated exposure of *C. perfringens* cultures to RumC1 over 30 days yielded only low-level resistance, with MIC values increased at most four-fold, in contrast to the several-log increases in MIC typically observed with conventional antibiotics^13^.

Here, we extended the characterization of the puzzling mode of action of RumC1 in the sensitive species *Streptococcus pneumoniae* (the pneumococcus). To this end, we combined genetic, biochemical and single-cell imaging approaches in a laboratory strain of this prominent human pathogen. In particular, following the screening and functional study of clones resistant to RumC1 toxicity, we visualized RumC1 interaction with individual cells during the cell cycle, and characterized the self-protection mechanism mediated by six putative immunity proteins encoded in its BGC. Altogether, we provide evidence that RumC1 operates via an unprecedented antibacterial mechanism, involving direct interaction with newly assembled PG in the cell wall of growing cells.

## Results

### Selective mutagenic screening by natural transformation and characterization of RumC1-resistant mutants in *S. pneumoniae*

To obtain and identify RumC1-resistant mutants in *S. pneumoniae*, we developed an ordered genome wide mutagenesis strategy taking advantage of the ability of this species to undergo natural genetic transformation at high frequency^17^. Natural transformation is a horizontal gene transfer mechanism enabling cells to uptake and integrate exogenous DNA by homologous recombination at a complementary chromosomal locus (referred hereafter to as *donor* and *recipient* DNA, respectively). Generating mutagenic PCR fragments under low-fidelity conditions resulted in a pool of donor DNA molecules capable of introducing random point mutations into the recipient genomic locus by transformation, providing an efficient means to screen for transformants with a selectable phenotype. This selection by natural transformation was performed at the whole genome scale by using ordered arrays of such mutagenic PCR fragments that cover the entire circular pneumococcal chromosome^18,19^. We implemented this mutagenic strategy in a laboratory strain of *S. pneumoniae* mutated for the DNA mismatch Hex repair system to improve recombinational integration of point mutations, routinely resulting in 100 % transformation efficiency for a single selective mismatch^20^. This method enables efficient screening for transformants exhibiting a selectable phenotype - such as resistance to RumC1 - while mutations can be precisely mapped and validated through re-amplification, high-fidelity sequencing, and inheritance testing in wild-type strains. In total, 505 mutagenic PCR fragments, each approximately 4.5 kb in length and overlapping by about 0.5 kb at their extremities, were necessary to cover the entire 2 Mb-long pneumococcal chromosome (Fig. 1A; Fig. S2A). These randomly mutated PCR fragments were grouped into 21 distinct pools (numbered P1 to P21), clockwise from the chromosomal replication origin and composed of 10 to 26 consecutive PCR fragments before transformation and antibiotic selection (Fig S2A-B).

**Figure 1:**
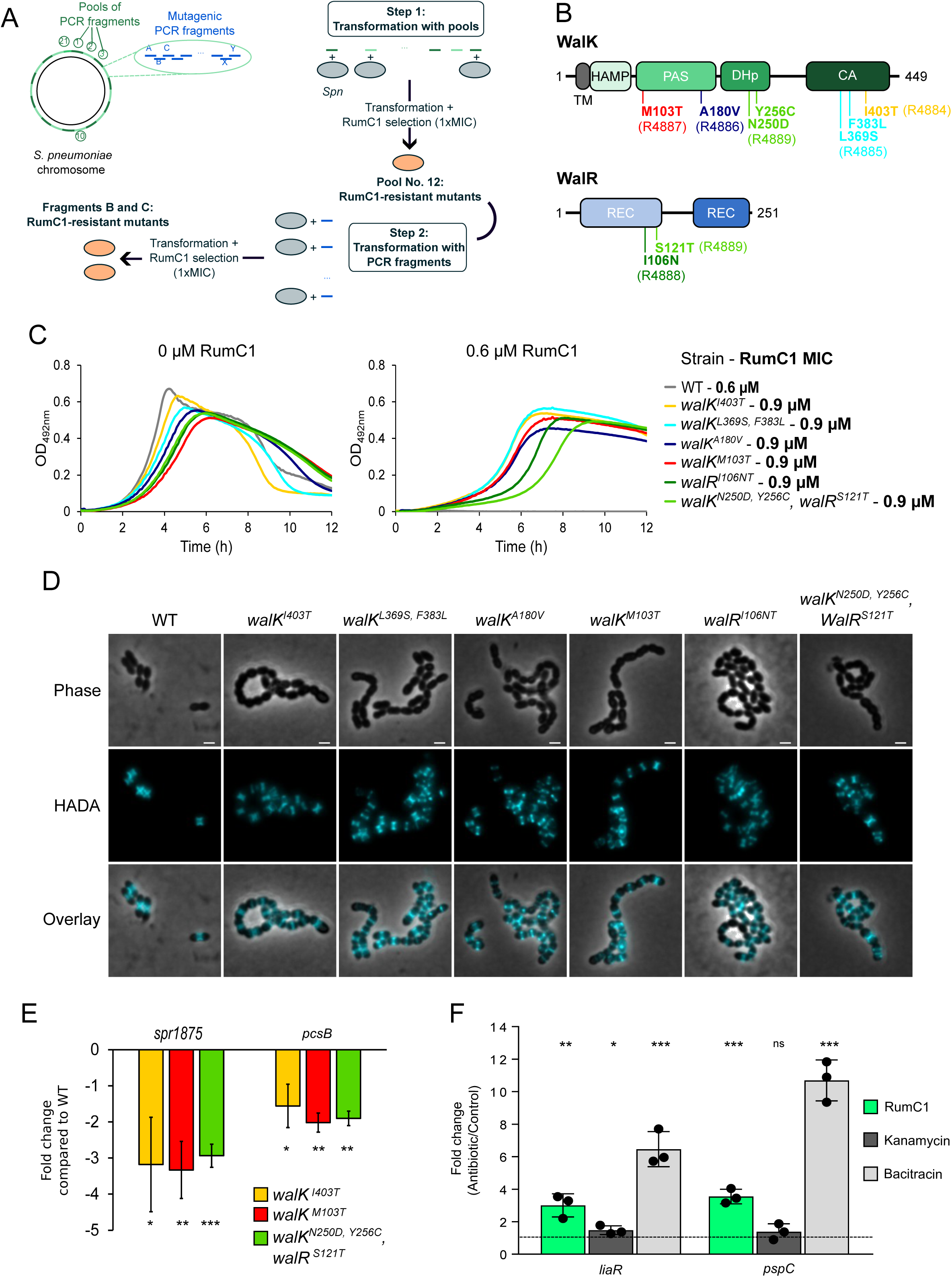
Pneumococcal RumC1-resistant mutants map in the essential WalRK two-component system that coordinates cell-wall homeostasis. (A) Schematic representation of the two-step NT-based mutagenesis method used to identify RumC1-resistant mutants. In the first step, *S. pneumoniae* R1818 (*comC0, hexAΔ3::ermAM*) competent cells were transformed with each of the 21 independent pools of mutagenic amplicons constituting the library and then selected on plates containing 3.5 µM RumC1 (MIC defined for these conditions, see materials ant methods). RumC1-resistant mutants were obtained following transformation with pool No. 12. A second transformation and RumC1 selection step was performed with individual amplicons of the pool No. 12. RumC1-resistant mutants were obtained following transformation with amplicons B and C. A detailed description of the mutagenic PCR fragment library and the description and calibration of the NT-based method used to screen for antibiotic-resistant strains is presented in Fig. S1. *Spn*: *Streptococcus pneumoniae*. (B) Mutations identified in RumC1-resistant strains. Domains organization of WalK and WalR is indicated. HAMP: Histidine kinases, Adenyl cyclases, Methyl-accepting proteins and Phosphatases domain; TM: Transmembrane domain; PAS: *Per-ARNT-Sim domain;* DHp: dimerization and histidine phosphorylation domain; CA: catalytic and HATPase (ATP-binding) domain; REC: Receiver domain; REG: Regulatory (DNA-binding) domain. The six RumC1-resistant strain names are indicated in brackets. (C) Growth of wild-type (strain R3584) and RumC1-resistant strains (strains R4884, R4885, R4886, R4887, R4888 and R4889) in the absence or in the presence of 0.6 µM RumC1. The RumC1 MIC of each strain is indicated in brackets. Data shown are representative of three independent experiments. Additional data used for MIC determination are presented in Fig. S3A. (D) Morphology and HADA incorporation in wild-type (strain R3584) and RumC1-resistant cells (strains R4884, R4885, R4886, R4887, R4888 and R4889) of *S. pneumoniae*. Phase contrast, fluorescent (false-colored in light blue), and false-colored overlay images are shown. Scale bar, 1 µm. Images are representative of three independent replicates. (E) Expression level of two WalR-regulated genes (*pcsB* and *spr1875*) in RumC1-resistant mutants. WT (R3584) and RumC1-resistant strains (R4884, R4887 and R4889) were grown until early log phase (OD_550_=0.1) before RNA extraction and RT-qPCR. The relative expression of each gene was standardized with the expression of *rpoD*. Results are expressed as fold changes in gene expression level relative to that in the wild-type strain. Standard deviations from three independent replicates are indicated. Statistical analysis was performed using an unpaired t-test (*, p-value < 0,05; **, p-value < 0,01; ***, p-value < 0,001). (F) RumC1 activates the cell wall stress-sensing two-component system LiaFSR. Relative expression of two genes of the LiaFSR regulon (*liaR* and *pspC*) determined by RT-qPCR is indicated. WT strain (R3584) was incubated with no antibiotic or with RumC1 (1.2 µM), Kanamycin (250 µg mL^-1,^ negative control) or Bacitracin (80 µg mL^-1,^ positive control) for 10 min before RNA extraction. For each gene, the relative expression in the presence of each antibiotic is standardized with the *rpoD* gene amplification and given versus the control condition with no antibiotic. Data are represented as individual data points with bars and error bars indicating averages ± SD, (*n* = 3, measured over three independent biological replicates). Standard deviations are indicated. Statistical analysis was performed using an unpaired t-test (n.s., non-significant, p-value > 0,05; *, p-value < 0,05; **, p-value < 0,01; ***, p-value < 0,001).

We first calibrated this methodology by screening for resistance to streptomycin (Sm^R^) and rifampicin (Rif^R^), which are well-known to arise from single point mutations in the ribosomal gene *rpsL* and the RNA polymerase subunit gene *rpoB*, respectively. Sm^R^ and Rif^R^ transformants were readily obtained with a single pool of PCR fragments and, next, with a single PCR fragment containing either the *rpsL* or *rpoB* gene, respectively (Fig. S2C-F). We found that this procedure markedly improved the rate of Sm^R^ and Rif^R^ mutations up to 5 logs compared to the spontaneous mutation rates in the *hex*^-^ strain. Indeed, Sm^R^ and Rif^R^ transformants were obtained at a frequency of about 10^-4^ and 10^-3^ when using pools or individual PCR fragments, respectively (Fig S2C-F).

We next applied this procedure twice to screen for RumC1-resistant transformants able to grow on solid medium supplemented with RumC1 at concentrations just above the MIC value established in those conditions (3.5 µM; see M&M). RumC1-resistant mutants were exclusively obtained with the P12 pool of mutagenic donor PCR fragments, with an initial frequency of 10^-6^ (Fig. 1A). This frequency improved to 10^-5^ in transformation assays with two consecutive fragments from the same pool (Fragments B and C, Fig. 1A). We confirmed the heritability of the newly acquired RumC1 resistance by transferring the corresponding PCR fragments B and C generated under high fidelity conditions from 6 randomly picked resistant clones into a *hex*+ isogenic strain. Sequencing of these PCR fragments revealed that resistance was consistently associated with single or double missense mutations in two adjacent genes, *walR* and *walK*, which encode components of a conserved two-component system (TCS) in Firmicutes^21^ (Fig. 1B). In particular, this TCS plays a critical role in coordinating PG synthesis and maturation during cell growth in *S. pneumoniae*, *Staphylococcus aureus* and *Bacillus subtilis*, in which it controls the transcription of a species-specific subset of genes^22–24^. The mutations identified in *walR* and *walK* varied among resistant clones and mapped to distinct functional domains of these proteins, reinforcing their involvement in RumC1 resistance. These findings suggest that RumC1 impacts cell wall integrity, as resistance mutations are exclusively localized to a regulatory system essential for PG biosynthesis.

### Functional and phenotypic analysis of RumC1-resistant mutants in *S. pneumoniae*

Further analysis of the six isolated RumC1-resistant mutants showed that their gained resistance against RumC1 was consistently weak, as none could grow at concentrations exceeding twice the MIC of RumC1 established for the wild-type strain (Fig. 1C and S3A). In addition, they exhibited a reduced growth rate in liquid culture, with variations observed among individual mutants. This growth impairment became even more pronounced in the presence of RumC1 applied at the MIC value of the parental strain (Fig. 1C). Thus, the limited protection conferred by these mutations against RumC1 toxicity appears to come at the cost of a compromised cellular fitness.

Importantly, distinct point mutants in *walR* or *walK* providing resistance to the PG-targeting antibiotic vancomycin have been repeatedly identified in clinical *S. aureus* isolates^25–27^. These isolates exhibited only a minor rise in vancomycin MIC, defining them as VISA (<u>V</u>ancomycin intermediate *<u>S</u>. <u>a</u>ureus*)^28^. Like the RumC1-resistant pneumococcal mutants, VISA strains also display a slower growth rate. In addition, VISA mutants exhibit a diminished transcription of the *walRK* regulon, resulting in various structural modifications of the cell envelope^29–31^.

The pneumococcal *walRK* regulon encodes three well-defined cell wall hydrolases: PcsB, a DL-endopeptidase that localizes to the division septum and is the only essential effector of the regulon; VldE (Spr1875), an LD-endopeptidase that cleaves cross-linked stem peptides in the PG; and LytB, a glucosaminidase acting in daughter cell separation^22,32,33^.To assess the transcriptional impact of RumC1 resistance, we measured the basal transcription levels of *pcsB* and *spr1875*, the latter being the most highly expressed gene in the pneumococcal *walRK* regulon^22,34^, using RT-qPCR in three resistant mutants. In all cases, basal transcription of these genes was diminished by 3- to 4-fold for *spr1875* and by 1.5- to 2-fold for *pcsB* (Fig. 1E). These findings indicate that a transcriptional downregulation of the WalRK regulon correlates with increased resistance to RumC1 in pneumococci, a pattern also reported in VISA mutants^30^. However, unlike VISA, all six RumC1-resistant derivatives remained sensitive to vancomycin as the parental strain (Fig. S3B).

Next, we observed these mutants at the single-cell level and we analyzed the substitution of the terminal D-Alanine (D-Ala) residue of the pentapeptide of the PG with the fluorescent D-Ala analog 7-hydroxycoumarincarbonylamino-D-alanine (HADA). This futile reaction allows the visualization of the transpeptidase activity at mid-cell of growing pneumococcal cells using fluorescent microscopy^35,36^. First, bright-field microscopy revealed a common defect in cell separation for all these mutants, resulting in a chaining phenotype, with a little effect on cell shape that appeared to be wider and squarer than the ovoid form of wild-type cells (Fig. 1D). The chaining phenotype is not sufficient to improve resistance to RumC1, as evidenced by the unchanged sensitivity of a *lytB* mutant, which forms large chains (Fig. S3C). Instead, the gained resistance in *walRK* mutants likely arises from a reduced action of another gene, or a combination of genes, within the regulon. Similar to the WT strain, all six RumC1-resistant mutants exhibited mid-cell HADA labeling patterns, but at reduced levels (Fig. 1D), a trend that correlated with their reduced growth rates (Fig. 1C). These results suggest that cells may gain partial protection against RumC1 by reducing the rate of PG assembly. Reciprocally, we wondered whether RumC1 exposure triggers a transcriptional response in the WalRK-system. To test this, we measured by RT-qPCR *pcsB* and *spr1875* expression in cells grown without RumC1 or with RumC1 for 10 min. As shown in Figure S3D, RumC1 did not change the basal transcriptional level of these genes. In parallel, we examined the LiaFSR system, which is a well-known pneumococcal TCS system that senses and signals PG damages by inducing the expression of a distinct subset of genes, including its own activation^37^. RT-qPCR analysis of *liaR* and *pspC*, key genes of the LiaFSR regulon, revealed that a 10-min treatment with 1.2 µM RumC1 induced the expression of both genes in comparison with untreated cells (Fig. 1F). Thus, defects mediated by RumC1 are sensed by LiaFSR, similar to other bacteriocins or antibiotics known to target PG synthesis^37^.

Altogether, these analyses demonstrate that RumC1 toxicity in *S. pneumoniae* can be partially mitigated by downmodulating the WalRK system responsible for PG homeostasis. In addition, the activation of the cell envelope stress sensor LiaFSR further support the conclusion that RumC1 interferes with cell wall integrity, providing mechanistic insight into its antibacterial mode of action.

### RumC1 alters the rate of PG assembly

To further investigate how RumC1 interferes with PG assembly, we analyzed its impact on the growth rate and viability of pneumococcal cells in liquid cultures, in parallel to its effect on cell morphology and PG biogenesis at the single-cell level over short periods of time, i.e. within the time frame of 30 min, corresponding to the duration of one cell-cycle. We determined three RumC1 concentrations (0.6, 1.2 and 3 µM) that gradually affected growth and viability of pneumococcal cells when added during early log phase (OD_550_ = 0.1; see M&M). In these conditions, 0.6 µM of RumC1 provoked a slight reduction in growth rate without affecting cell viability, as measured by colony forming units (CFUs; Fig. 2A and 2B). At 1.2 µM of RumC1, cellular growth stopped rapidly, though viability remained largely unaffected. By contrast, exposure to 3 µM RumC1 not only blocked growth but also resulted in rapid cell death, as indicated by the loss of CFUs within 10 min of treatment (Fig. 2A and B).

**Figure 2:**
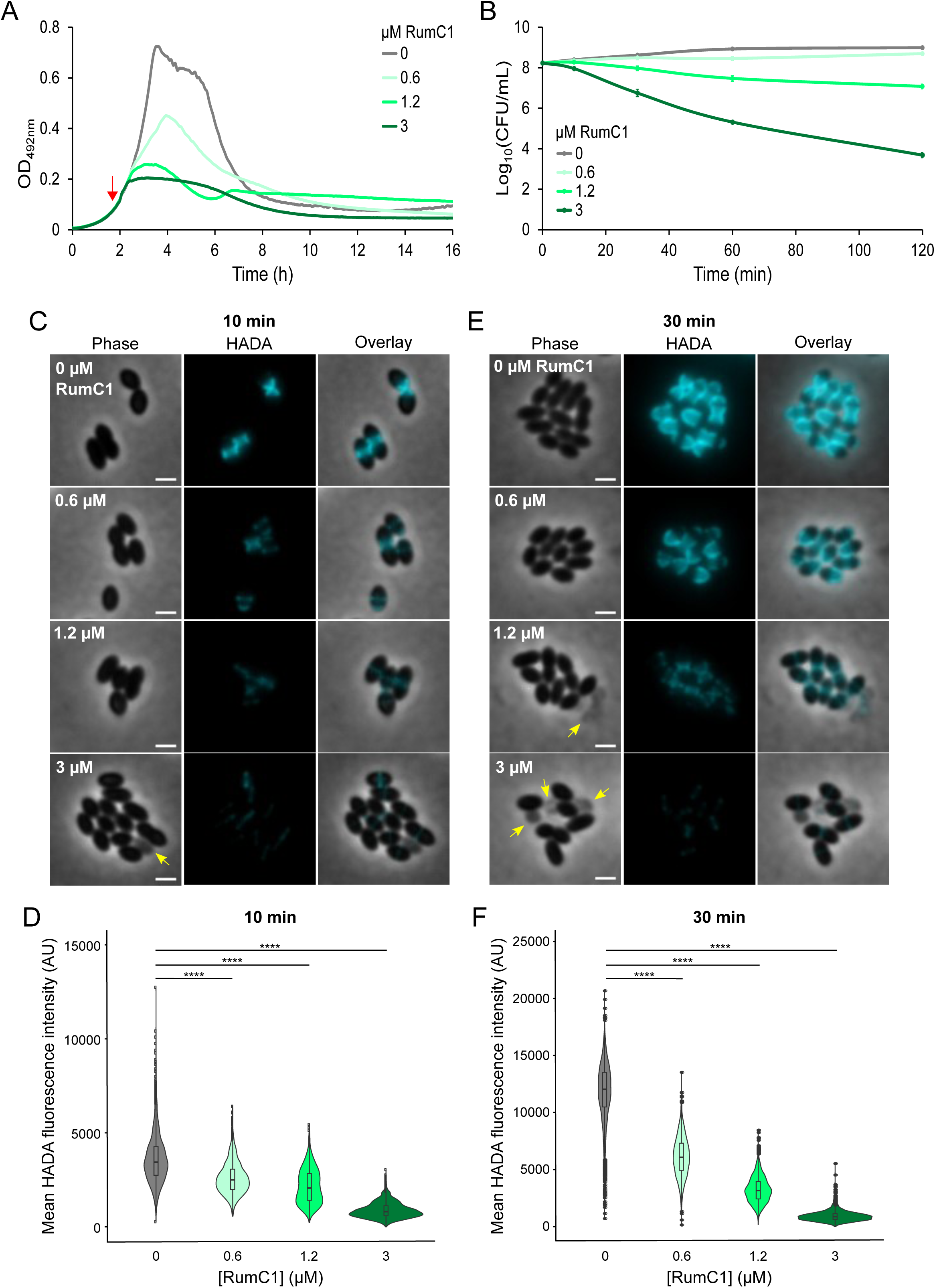
RumC1 affects peptidoglycan synthesis in *S. pneumoniae*. (A) Effect of RumC1 on growth of high-cell-density cultures of a wild-type strain of *S. pneumoniae* (R1501). Increasing concentrations of RumC1 were added at OD_550_=0.1 (red arrow). Data shown here are representative of three independent replicates. (B) Time-kill kinetics of RumC1 against early log phase cultures of *S. pneumoniae* R1501. CFU were determined after 10, 30, 60 and 120 min of treatment with 0, 0.6, 1.2 and 3 µM RumC1. Mean and standard deviations obtained from three independent experiments are indicated. (C) Morphology and HADA incorporation in wild-type cells of *S. pneumoniae* (strain R1501) untreated or treated with 0.6, 1.2 or 3 µM RumC1 for 10 min at 37°C. Phase contrast, fluorescent (false-colored in light blue), and false-colored overlay images are shown. The yellow arrows indicate “ghost cells”. Scale bars, 1 µm. Images are representative of three independent replicates. (D) Violin plots representing the mean HADA fluorescence intensity in WT (R1501) cells untreated (n=1446) or treated with 0.6 (n=2082), 1.2 (n=1765) or 3 µM (n=1692) RumC1 for 10 min at 37°C. Boxes extend from the 25^th^ percentile to the 75^th^ percentile, with the horizontal line at the median. Dots represent outliers. Statistical analysis was performed using the U test of Mann-Whitney (****, p-value < 0,0001). (E) Morphology and HADA incorporation in wild-type cells of *S. pneumoniae* (strain R1501) untreated or treated with 0.6, 1.2 or 3 µM RumC1 for 30 min at 37°C. Phase contrast, fluorescent (false-colored in light blue), and false-colored overlay images are shown. The yellow arrows indicate “ghost cells”. Scale bars, 1 µm. Images are representative of three independent replicates. (F) Violin plots representing the mean HADA fluorescence intensity in WT (R1501) cells untreated (n=3400) or treated with 0.6 (n=1791), 1.2 (n=2474) or 3 µM (n=1335) RumC1 for 30 min at 37°C. Boxes extend from the 25^th^ percentile to the 75^th^ percentile, with the horizontal line at the median. Dots represent outliers. Statistical analysis was performed using the U test of Mann-Whitney (****, p-value < 0,0001).

To assess the impact of RumC1 on transpeptidase activity, the HADA probe was added alongside each RumC1 dose to exponentially growing pneumococci and incorporation was monitored after 10 and 30 min. As seen in Fig. 2C-F, RumC1 led to a significant and dose-dependent decrease in HADA incorporation, detectable as early as 10 min of exposure to the lowest RumC1 concentration (0.6 µM), showing that RumC1 rapidly alters global transpeptidase activity. Importantly, cells treated with RumC1 still incorporated HADA at mid-cell, indicating that RumC1 does not alter the positioning of the cell wall synthesis machinery. Besides this effect on HADA incorporation, RumC1 caused the emergence of cells with a pale, “ghost-like” aspect (Fig. 2C and E, yellow arrows), likely reflecting reduced cytoplasmic density and a loss of cellular content. This bacteriolytic effect appeared at 3 µM of RumC1 after 10 min, and at 1.2 µM after 30 min, which is consistent with the loss of CFUs under these conditions (Fig. 2B, C and E). While RumC1 did not significantly alter overall cell shape, it did reduce the average cell length at concentrations above 3 µM, potentially indicating an inhibition of cell elongation (Fig. 2C, E and S4). These effects of RumC1 on cell morphology and PG synthesis differ from those caused by vancomycin, a well-characterized antibiotic interfering with late steps of PG assembly. Indeed, after 30 min of exposure to a low dose of vancomycin (0.17 µM) that caused a growth rate reduction equivalent to 0.6 µM RumC1, cells became elongated and incorporated at least as much HADA as untreated cells. At a higher dose of vancomycin (0.34 µM; equivalent to the lethal 3 µM RumC1 concentration), ghost cells emerged, as well as cells with polar HADA foci, suggesting delocalization of PG synthesis sites or altered PG dynamics (Fig. S5).

Taken together, these results show that RumC1 rapidly impairs PG synthesis, even at sub-lethal concentrations, before inducing cell death at higher doses or after prolonged exposure. In addition, RumC1-mediated disruption of PG assembly appears to occur through a mechanism distinct from that of vancomycin.

### RumC1 binds to a neo-assembled PG intermediate

Next, we studied the physical interaction of RumC1 with individual growing cells by fluorescence microscopy. To this end, we covalently linked the fluorescent dye Alexa Fluor 488 to mature RumC1 (see M&M). Following purification, AF488-RumC1 remained in monomeric form as RumC1 (Fig. S6A), and exhibited only a minor reduction in antibacterial activity against *S. pneumoniae* compared to the native peptide (Fig. S6B). When exponentially growing cells were exposed to a sub-inhibitory concentration of AF488-RumC1 (2.5 µM, Fig. S6C) for 10 min, we observed a distinctive binding pattern: two fluorescent bands emerged at mid-cell, flanking the unlabeled central division zone where nascent PG is synthesized to generate the typical ovoid shape of pneumococci (Fig. 3A). Co-labelling experiments with HADA, used as a marker of PG synthesis, confirmed that AF488-RumC1 accumulated on both sides of the central HADA incorporation area, indicating that AF488-RumC1 preferentially interacts with an early PG intermediate (Fig. 3B). In addition, measurement of HADA fluorescence intensity revealed that AF488-RumC1 reduced transpeptidase activity, as found with the unlabeled RumC1 at the same concentration (Fig. S6D and 2E). When AF488-RumC1 was applied for 10 min at an inhibitory concentration for growth (i.e.12.5 µM, Fig. S6C), the fluorescence pattern shifted: AF488-RumC1 accumulation intensified and localized as a single and bright fluorescent band at the division site (Fig. 3A). Importantly, the AF488-RumC1-A12E mutant which lacks antibacterial activity^16^, failed to accumulate on pneumococcal cells even at high concentrations (i.e. 11.9 µM), underscoring that RumC1’s binding to the PG synthesis area is directly linked to its toxicity (Fig. 3C). By comparison, fluorescently labelled vancomycin (BODIPY*™* FL Vancomycin (van-Fl)) did not label dividing pneumococcal cells at sub-inhibitory concentration (0.17 µM), but formed a single fluorescent band at mid-cell at a higher, inhibitory concentration (0.34 µM; Fig. S5). This differential labeling further distinguishes RumC1’s interaction with the pneumococcal PG from that of vancomycin.

**Figure 3:**
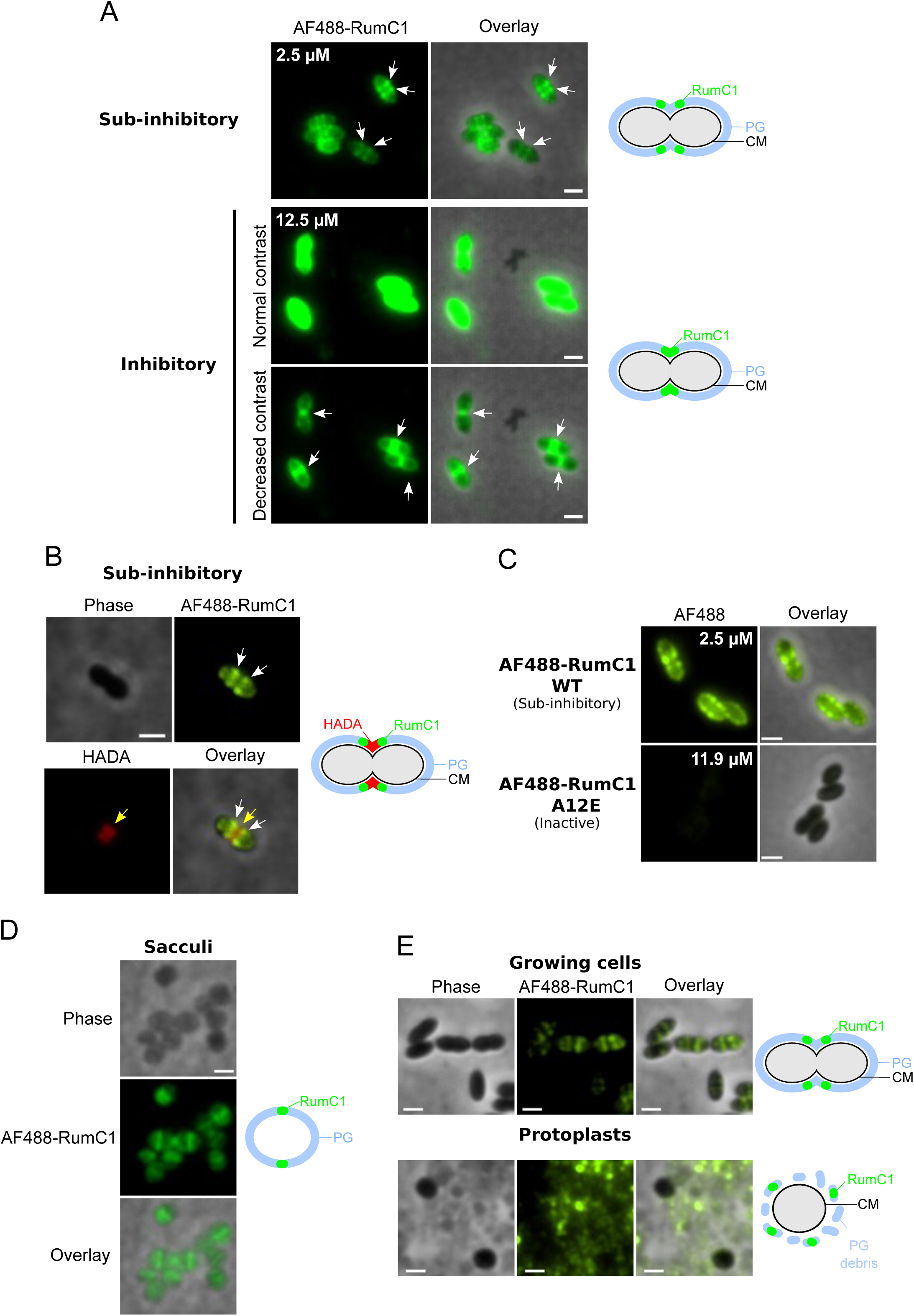
RumC1 binds to a neo-nascent PG component. (A) Binding of AF488-RumC1 to pneumococcal cells. Exponentially growing cells of a wild-type strain of *S. pneumoniae* (R1501) were labelled for 10 min with a sub-inhibitory (2.5 µM) or inhibitory (12.5 µM) concentration of AF488-RumC1. The fluorescence of AF488-RumC1 when applied at 12.5 µM was so high that both pictures with normal contrast and decreased contrast are shown. White arrows: AF488-RumC1. Scale bars, 1 µm. Images are representative of at least three independent replicates. Right panels: schemes representing the binding of AF488-RumC1 to growing cells at sub-inhibitory and inhibitory concentrations. CM: Cell Membrane (B) Co-labelling experiments with AF488-RumC1 and HADA. Exponentially growing cells of a wild-type strain of *S. pneumoniae* (R1501) were labelled for 10 min with a sub-inhibitory (2.5 µM) concentration of AF488-RumC1 and HADA (false-colored in red) before washing and imaging. White arrows: AF488-RumC1, yellow arrows: HADA. Scale bar, 1 µm. Images are representative of at least three independent replicates. The quantification of the HADA signal from cells exposed to AF488-RumC1 compared to unexposed cells is presented in Fig. S5D and demonstrates that AF488-RumC1 decreases transpeptidase activity, as is the case with the unlabeled version of RumC1. Right panel: scheme representing the respective localization of AF488-RumC1 and HADA in growing cells. (C) Binding of the innocuous derivative AF488-RumC1 A12E to pneumococcal cells. Exponentially growing cells of a wild-type strain of *S. pneumoniae* (R1501) were labelled for 10 min with 2.5 µM AF488-RumC1 WT or 11.9 µM AF488-RumC1 A12E before washing and imaging. Scale bars, 1 µm. Images are representative of two independent replicates. (D) Binding of AF488-RumC1 to pneumococcal sacculi. Sacculi isolated from a wild-type strain of *S. pneumoniae* (R1501) were labelled for 10 min with 2.5 µM AF488-RumC1 before washing and imaging. Scale bar, 1 µm. Images are representative of four independent replicates. Right panel: scheme representing the binding of AF488-RumC1 to sacculi. (E) Binding of AF488-RumC1 to pneumococcal protoplasts. Protoplasts isolated from a wild-type strain of *S. pneumoniae* (R1501) were labelled for 10 min with 2.5 µM AF488-RumC1 before washing and imaging. A control of live cells resuspended in the ‘protoplastisation’ buffer (see Materials and Methods) labelled with AF488-RumC1 was run in parallel. The background fluorescence signal probably corresponds to AF488-RumC1 bound to PG debris generated during protoplastisation. Scale bars, 1 µm. Images are representative of two independent replicates. Right panels: schemes representing the binding of AF488-RumC1 to growing cells and protoplasts.

To confirm whether RumC1 mid-cell localization resulted from a direct interaction with a PG component, we analyzed its binding to purified, deproteinized sacculi prepared from exponentially growing pneumococci. As shown in figure 3D, AF488-RumC1 accumulated on these sacculi, generating a bright fluorescent central band. This pattern is reminiscent of the mid-cell localization observed in growing cells at high AF488-RumC1 concentrations, showing that RumC1 specifically targets newly assembled PG rather than mature PG of the mother cell. Conversely, AF488-RumC1 showed no significant interaction with pneumococcal protoplasts devoid of their cell wall, but did bind to co-purified PG debris resulting from their production (Fig. 3E).

Altogether, these microscopy analyses reveal that RumC1 selectively accumulates on the nascent PG, and concurrently inhibits its synthesis.

### The RumI_c_1 immunity protein of *R. gnavus* E1 is sufficient to counteract RumC1 toxicity

Almost all bacteriocin-producing monoderm bacteria co-express dedicated immunity proteins to protect themselves against their noxious effect. Some immunity proteins interact with bacteriocins to prevent them from reaching their targets, while others act on the bacteriocin target^38–42^. To gather additional cues on the mode of action of RumC1, we undertook a functional study of its predicted immunity system in *S. pneumoniae*, which is encoded by a putative operon of six genes within the RumC1 BGC (Fig S1A)^11^. Sequence analysis and AlphaFold3 modeling revealed that the first ORF in this operon, *rumI_c_1*, encodes a protein with a short cytoplasmic N-terminal domain, a transmembrane segment, and an extracellular domain belonging to the C70 peptidase subfamily (Fig. S7). The following ORF, *rumI_c_2*, encodes a protein with a tripartite signal sequence characteristic of lipoproteins, consisting of a positively charged N-region, a hydrophobic region and a lipobox^43^ (Fig. S7). Next, *rumE_c_*, *rumG_c_* and *rumF_c_* ORFs encode components of an ABC transporter. Both RumE_c_ and RumG_c_ exhibit four transmembrane segments and a FtsX-like permease domain, while RumG_c_ contains an additional MacB-like Periplasmic Core Domain, both of which are commonly found in transmembrane domains (TMDs) of ABC transporters. RumF_c_ comprises a cytoplasmic Nucleotide-Binding Domain (NBD) typical of ABC transporters (Fig. S7). Then, *rumY_c,_* located between *rumE_c_* and *rumF_c_*, encodes a hypothetical cytosolic protein of 135 amino acids, for which no known functional or structural signature have been identified so far (Fig. S7).

To test whether these candidate immunity proteins could provide protection against RumC1 toxicity in *S. pneumoniae*, we introduced different combinations of their genes in the chromosome under the control of the synthetic IPTG-inducible P*_lac_* promoter^8^. We first assayed the individual expression of *rumI_c_1*, *rumI_c_2* and the ABC transporter *rumE_c_F_c_G_c_*. To this end, we constructed three strains carrying the chromosomal expression platforms *CEP-P_lac_-rumI_c_1*, *CEP-P_lac_-rumI_c_2* and *CEP-P_lac_-rumE_c_Y_c_F_c_G_c_*, and monitored their ability to grow in liquid medium supplemented with IPTG and increasing concentrations of RumC1, using a control strain expressing the firefly luciferase from the *CEP-P_lac_-luc* construct^44^. Cell growth was not affected by the expression of any of these immunity genes (Fig. 4). We then incrementally increased RumC1 concentration in 0.2 µM steps, starting from a sub-inhibitory concentration of 0.4 µM, to determine whether IPTG-induced expression of these constructs modified the 0.6 µM MIC value of RumC1 established for the strain containing the *CEP-P_lac_-luc* construct (Fig. 4). Of note, RumC1 applied at 0.4 µM impaired the growth of the control strain, resulting in a marked growth delay of the cell population (Fig. 4). Individual expression of RumI_c_1 mitigated this growth delay at this sub-inhibitory concentration and provided partial protection at higher, inhibitory doses of RumC1 (Fig. 4). However, the protective effect of RumI_c_1 diminished at elevated RumC1 concentrations, and the overall gain in resistance remained modest, raising the MIC by less than two-fold. This partial resistance was associated with improved tolerance, as *rumI_c_1* expression enhanced the survival of cells transiently exposed for 1 h to high doses of RumC1 (Fig. S8). In contrast, neither *rumI_c_2* nor *rumE_c_Y_c_F_c_G_c_* expression provided protection against RumC1 at any concentration tested (Fig. 4). Finally, it appeared that *rumI_C_2* expression further sensitized cells to RumC1, as shown by reduced growth at a sub-inhibitory concentration of RumC1 (0.4 µM).

**Figure 4:**
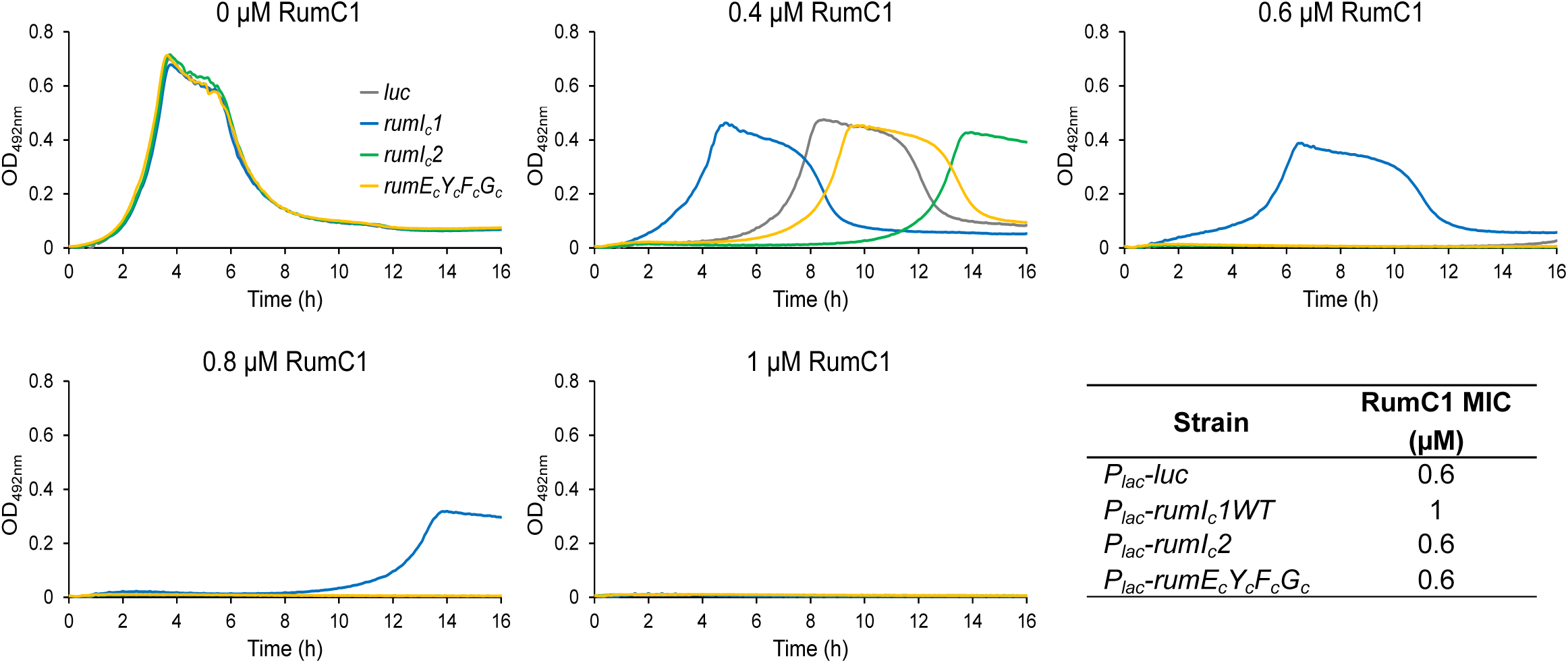
The immunity protein RumI_c_1 protects pneumococcal cells against RumC1. Growth of strains *CEPI-P_lac_-luc* (R5339), *CEPI-P_lac_-rumI_c_1* (R5231), *CEPI-P_lac_-rumI_c_2* (R5340) and *CEPI-P_lac_-rumE_c_Y_c_F_c_G_c_* (R5341), in the absence or presence of various concentrations of RumC1. The RumC1 concentrations are indicated on top of the graphs. The corresponding RumC1 MICs determined for each strain are listed in the table on the right. Precultures and cultures were performed in the presence of 50 µM IPTG. Data shown here are representative of three independent replicates.

Given that cooperation between the immunity proteins may be required for full bacteriocin resistance, we next examined whether combined expression could enhance protection^45,46^. To test this hypothesis, the immunity genes were integrated into the pneumococcal chromosome and expressed in a controlled and segmented manner using two tunable promoters, one by IPTG (*P_lac_)* and the other by anhydrotetracycline (ATC; *P_tet_*)^47^. We cloned *rumI_c_1* and *rumI_c_2* under the control of *P_lac_* at the chromosomal CEP locus, and *rumE_c_*, *Y_c_*, *F_c_* and *G_c_* under the control of *P_tet_* promoter at the separate chromosomal CEPII locus, respectively (Fig. S9A). The resulting strain was grown in a gradual range of RumC1 concentrations, with or without IPTG and ATC. Co-expression of *rumI_c_1* and *rumI_c_2* increased the resistance to RumC1 compared to wild-type, though the level of protection was lower than that achieved by *rumI_c_1* alone, likely due to the deleterious effect of *rumI_c_2* on growth in the presence of RumC1 (Fig. 4 and Fig. S9B). Expression of *rumE_c_*, *Y_c_*, *G_c_* and *F_c_* under *P_tet_* control only slightly improved the resistance to RumC1. Finally, the highest level of immunity was achieved when all six immunity genes were co-expressed, although the level of resistance remained modest and comparable to the RumC1 MIC value achieved with *rumI_c_1* alone (1 µM). Finally, to further explore the potential of RumI_c_1, we tested whether increasing its expression using both *P_lac_* and *P_tet_* promoters could enhance resistance to RumC1. As shown in Fig. S9C and D, cumulative expression of *rumI_c_1* only slightly increased resistance to RumC1 without changing the MIC value measured with a single copy of the gene. Altogether, these results showed that these genes, except *rumI_c_2*, provide immunity against RumC1 toxicity, with RumI_c_1 being necessary and sufficient for protection.

RumI_c_1 is predicted to be anchored in the cytoplasmic membrane, with a large, externally exposed globular domain (Fig. S7A and B). To enable heterologous expression in *E. coli*, we restricted the construct to the soluble catalytic domain (i.e., peptidase C70). This globular domain - named RumI_c_1_S_ - was purified to homogeneity and subsequently tested for its ability to protect *S. pneumoniae* cells against RumC1 when added to the growth medium (Fig. S10A). Remarkably, treatment with 50 to 400 µM RumI_c_1_S_ increased RumC1 MIC by 2-to 64-fold, indicating that RumI_c_1 can function outside the cell (Fig. S10B and C). We next tested whether this exogenous protection mediated by RumI_c_1 extended to *R. gnavus* E1, the original strain harboring the RumC1 BGC. *R. gnavus* E1 cells grown under laboratory conditions do not express the *rumC* regulon^6^ and are therefore sensitive to RumC1, with a MIC of 12.5 µM (Fig. S10D). Pre-treatment of *R. gnavus* E1 cells with 100 to 400 µM of RumI_c_1_S_ for 1.5 h increased the RumC1 MIC by 2 and 8-fold, respectively, confirming that RumI_c_1 is functional in its native host (Fig S10D and E).

Altogether, these results establish RumI_c_1 as an immunity protein directed against RumC1, capable of conferring protection both endo- and exogenously. Its ability to act independently of the other immunity proteins highlights its central role in neutralizing RumC1 toxicity.

### RumI_c_1 confers cross-resistance against RumC1 and vancomycin in *S. pneumoniae via* proteolytic activity

The extracellular domain of RumI_c_1 includes the typical signature of the C70 peptidases subfamily of the MEROPS database^48^ (Fig. S7). A representative C70 member is AvrRpt2, an effector of a Type III secretion system of *Pseudomonas syringae* that promotes virulence in *Arabidopsis thaliana* plants by cleaving host proteins *via* a catalytic triad comprising cysteine, histidine and aspartate residues^49–51^. Sequence alignments and structural prediction identified Cys145, His234 and Asp250 in RumI_c_1, defining the same catalytic triad as in AvrRpt2 (Fig. 5A-B). We individually substituted each residue with alanine in the P_lac_-*rumI_c_1* construct and tested the ability of the resulting mutant strains (*rumI_c_1^C145A^*, *rumI_c_1^H234A^*and *rumI_c_1^D250A^*) to confer resistance against RumC1. Unlike wild-type *rumI_c_1*, none of the mutated alleles enabled cell growth at the RumC1 MIC (0.6 µM) (Fig. 5C). Moreover, expression of *rumI_c_1^H234A^* and *rumI_c_1^D250A^* slightly increased sensitivity to a sub-inhibitory concentration of RumC1 (0.4 µM), in comparison with the *luc*-expressing strain and the *rumI_c_1^C145A^* mutant (Fig. 5C). In all, these results strongly suggest that RumI_c_1 protects cells against RumC1 through proteolytic activity.

**Figure 5:**
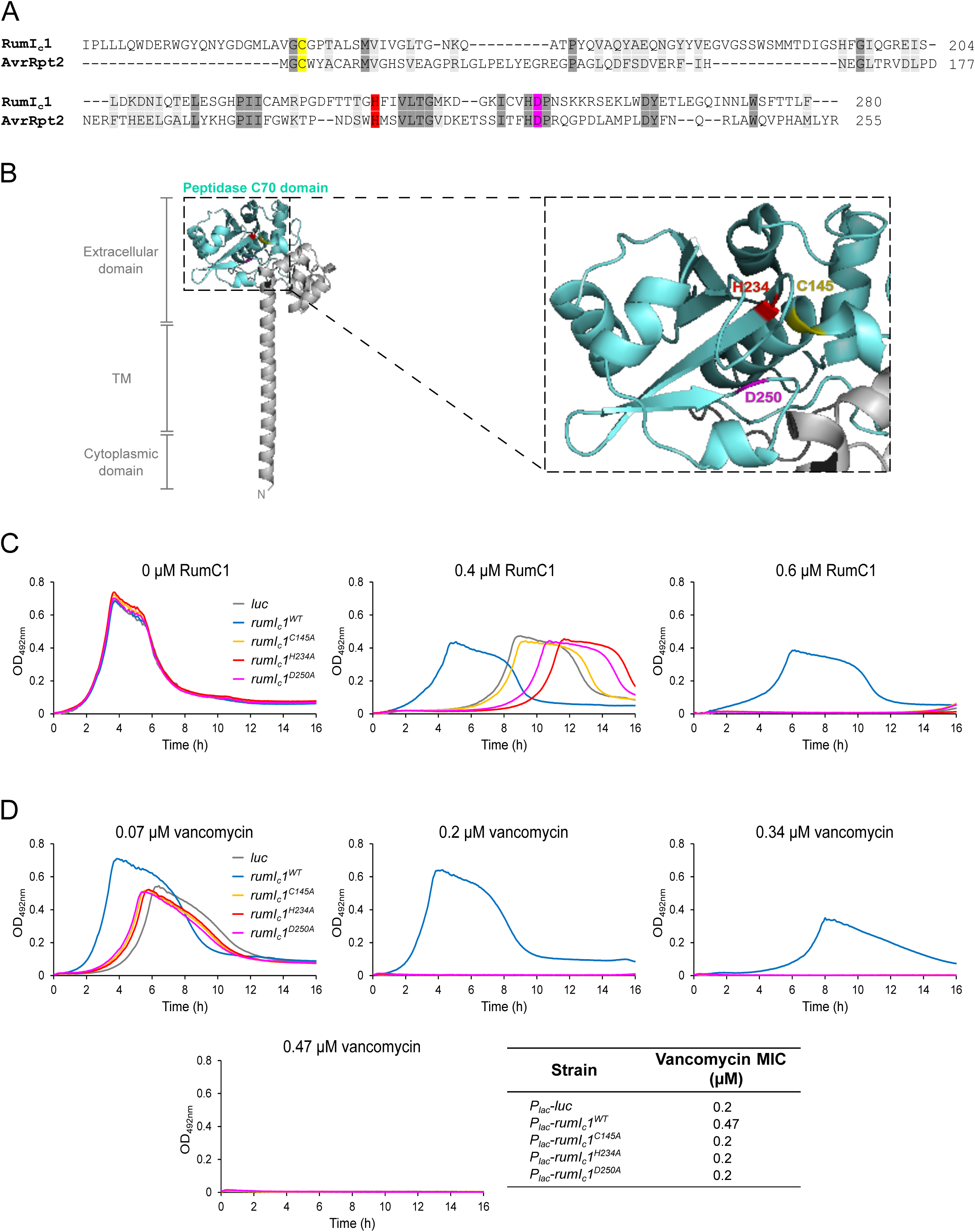
The RumI_c_1 protein confers cross-immunity to pneumococcal cells against RumC1 and vancomycin through a proteolytic activity. (A) Amino acids sequence alignment of the C70 peptidase domains of RumI_c_1 and AvrRpt2 of *Pseudomonas syringae*. Conserved residues between the two proteins are shaded in dark grey. Similar residues between the two proteins are shaded in light grey. The Cysteine, Histidine and Aspartate residues of the putative catalytic triad of RumI_c_1 and AvrRpt2 are highlighted in yellow, red and pink, respectively. (B) AlphaFold3 model of RumI_c_1. The predicted topology is indicated. The C70 peptidase domain is depicted in light blue. The position of the Cysteine, Histidine and Aspartate residues of the putative catalytic triad is indicated. (C) Growth of strains *CEPI-P_lac_-luc* (R5339), *CEPI-P_lac_-rumI_c_1^WT^* (R5231), *CEPI-P_lac_-rumI_c_1^C145A^* (R5336), *CEPI-P_lac_-rumI_c_1^H234A^* (R5337) and *CEPI-P_lac_-rumI_c_1^D250A^* (R5338), in the absence or presence of various concentrations of RumC1. The RumC1 concentrations are indicated on top of the graphs. Precultures and cultures were performed in the presence of 50 µM IPTG. Data shown here are representative of three independent replicates. (D) Growth of strains *CEPI-P_lac_-luc* (R5339), *CEPI-P_lac_-rumI_c_1^WT^* (R5231), *CEPI-P_lac_-rumI_c_1^C145A^* (R5336), *CEPI-P_lac_-rumI_c_1^H234A^* (R5337) and *CEPI-P_lac_-rumI_c_1^D250A^* (R5338), in the absence or presence of various concentrations of vancomycin. The vancomycin concentrations are indicated on top of the graphs. The corresponding vancomycin MICs of each strain are listed in the table in the right. Precultures and cultures were performed in the presence of 50 µM IPTG. Data shown here are representative of three independent replicates.

In parallel, we tested the protective effect of RumI_c_1 against several well-characterized antibiotics targeting different stages of PG synthesis. We evaluated ampicillin, which inactivates the PBPs transpeptidases, bacitracin, which sequesters the undecaprenyl pyrophosphate (UPP), daptomycin, which associates with the anionic phospholipid phosphatidylglycerol and the lipid II, and vancomycin, which specifically binds to the D-Ala-D-Ala terminus of the PG pentapeptide in the lipid II or in the assembled PG. We also included novobiocin, which damages the genome by inactivating the gyrase, and streptomycin, which inhibits protein synthesis. Expression of *rumI_c_1* had no significant effect on susceptibility to these antibiotics with the marked exception of vancomycin, resulting in a MIC value of 0.47 µM in comparison to the MIC of 0.2 µM measured for cells lacking *rumI_c_1* (Fig. 5D and S11). Although modest, this acquired vancomycin resistance, similar to that observed with RumC1, was further enhanced by dual expression of *rumI_c_1* under the control of both *P_tet_* and *P_lac_* promoters, reaching a MIC value of 0.61 µM (Fig. S12). In contrast, the points mutants of the catalytic triad of RumI_c_1 failed to protect cells against vancomycin (Fig. 5D). In all, these results suggest that RumI_c_1 mediates cross-protection against RumC1 and vancomycin, most probably through proteolytic activity.

### RumI_c_1 processes the stem peptide of PG to protect cells against RumC1 and vancomycin

Vancomycin is a glycopeptide that interferes with PG synthesis by interacting with the highly conserved terminal dipeptide D-Ala-D-Ala of the pentapeptide, thereby inhibiting transpeptidase-mediated cross-linking of nascent glycan strands^52^. The cross-protection conferred by RumI_c_1 against vancomycin and RumC1 raised the hypothesis that RumI_c_1 targets the D-Ala-D-Ala extremity of the stem peptide rather than directly inactivating these two unrelated antimicrobial peptides. To explore this hypothesis, we used a titration assay of vancomycin toxicity based on its pre-incubation with a synthetic pneumococcal pentapeptide prior to its addition to growing cells (Fig. S13A). A 50-fold molar excess of the pentapeptide over vancomycin at its MIC value fully alleviated vancomycin toxicity (Fig. S13B). Next, to test whether RumI_c_1 could inactivate the pentapeptide, either RumI_c_1_S_^WT^ or RumI_c_1_S_^C145A^ purified domains were incubated with the pentapeptide at a molar ratio of 1:20 prior to the addition of vancomycin (Fig. 6A). RumI_c_1_S_^WT^ pre-incubated with the pentapeptide totally abrogated its antagonistic effect on vancomycin, whereas RumI_c_1_S_^C145A^ had no effect (Fig. 6B). This indicates that RumI_c_1 either cleaves or masks the D-Ala-D-Ala extremity of the pentapeptide. The 1:20 molar ratio of RumI_c_1_S_ to pentapeptide, sufficient to alleviate the titration of vancomycin by the pentapeptide, is consistent with an enzymatic cleavage mechanism. Of note, RumI_c_1_S_^WT^ incubation with either vancomycin or RumC1 in the absence of the pentapeptide did not impact their toxicity, indicating that RumI_c_1 does not target these antimicrobial peptides but instead modifies the PG pentapeptide, a crucial determinant of their activity (Fig. S14). Also, while a 1:50 molar ratio of vancomycin to pentapeptide is sufficient to reduce vancomycin activity (Fig. S13A and B), a 1000-fold molar excess of the pentapeptide had no effect on RumC1 toxicity (Fig. S13A and C).

**Figure 6:**
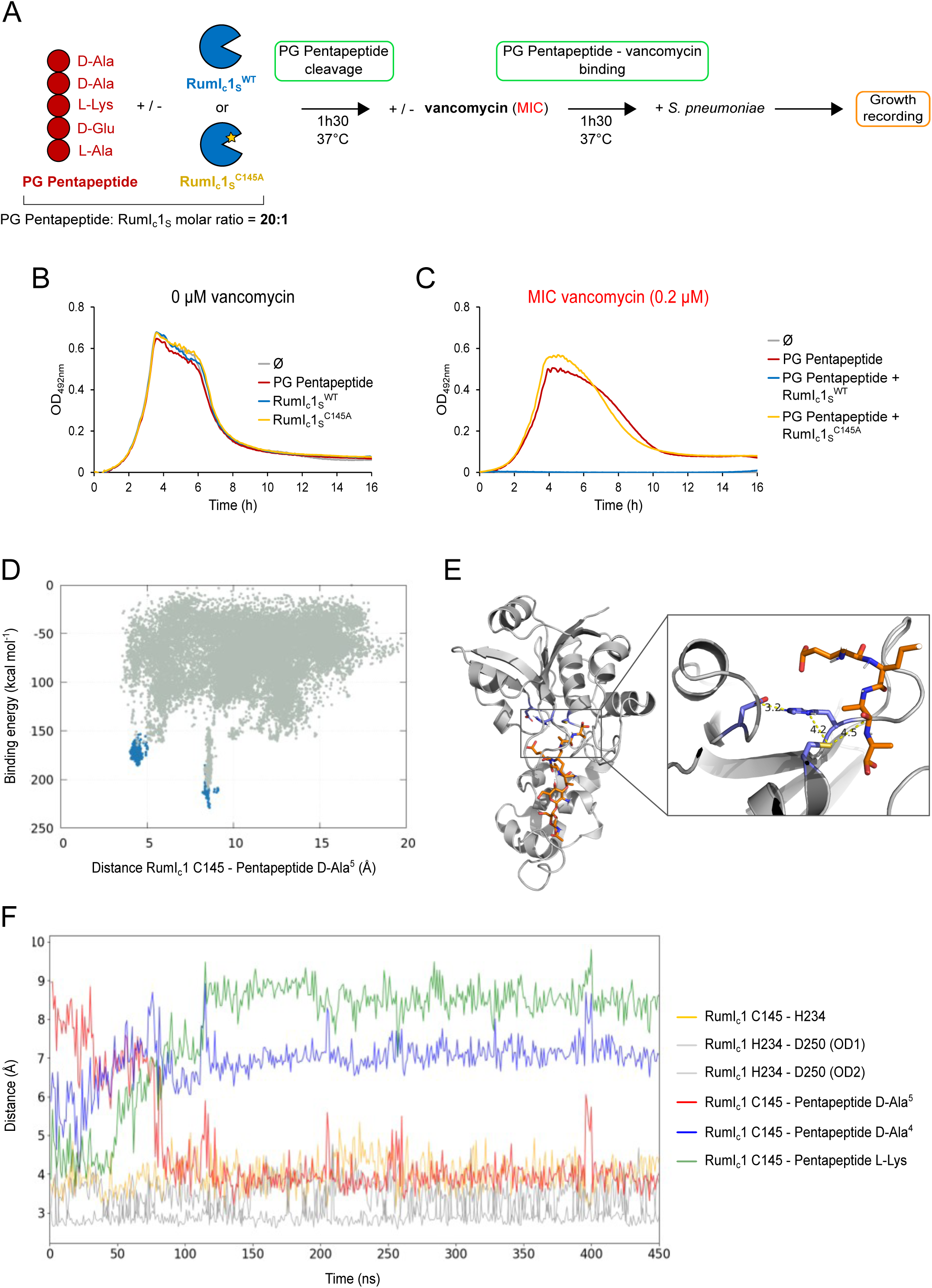
The immunity protein RumI_c_1 cleaves the PG Pentapeptide. (A) Schematic representation of experiments carried out to assess RumI_c_1 cleavage of the PG pentapeptide of *S. pneumoniae*. The chemically synthesized PG pentapeptide containing the nonamidated version of the peptide at position 2 (D-Glu instead of D-iGln) was incubated with the purified wild-type extracellular domain of RumI_c_1 (RumI_c_1_S_^WT^) or with a catalytic residue mutant of RumI_c_1 (RumI_c_1_S_^C145A^). Reaction mixtures were next incubated with vancomycin and added to *S. pneumoniae* cultures. (B) Growth of WT strain (R1501) in the presence of the PG pentapeptide and RumI_c_1_S_^WT^ or RumI_c_1_S_^C145A^. Data shown here are representative of two independent replicates. (C) Growth of WT strain (R1501) in the presence of a MIC concentration of vancomycin (0.2 µM) pre-incubated or not with the PG pentapeptide pre-treated with RumI_c_1_S_^WT^ or with RumI_c_1_S_^C145A^. Data shown here are representative of three independent replicates. (D) PELE energy landscape of the induced-fit simulation (gray) and subsequent refinement of the two lowest-energy minima (blue). (E) Structural representation of the catalytic pose of RumI_c_1 in complex with the pentapeptide, corresponding to the most populated MD cluster derived from the lowest-energy minimum identified in the PELE energy landscape. (F) Time evolution of key catalytic distances during one MD replica, including Cys145–His234, His234–Asp250 (OD1), and His234–Asp250 (OD2). The plot also shows the distance between the catalytic Cys145 and three potential cleavage bonds of the PG stem peptide substrate: the terminal D-alanine (D-Ala^5^), the penultimate D-alanine (D-Ala^4^), and the lysine cleavage sites. Initially, the substrate adopts a conformation in which Cys145 is positioned to cleave the lysine residue. However, after approximately 100 ns, the peptide reorients, placing the terminal D-alanine within catalytic distance of Cys145.

Together, these findings show that RumI_c_1 protects cells against vancomycin and RumC1 by processing the pentapeptide, although these two antimicrobial peptides operate *via* distinct mechanisms.

### RumIc1 has a D-D-carboxypeptidase activity in pneumococcal cells

To further investigate the binding and cleavage potential of the pentapeptide by RumI_c_1, we used PELE (Protein Energy Landscape Exploration) with the AlphaFold2 prediction of the RumI_c_1 structure (Fig. 6D). This Monte Carlo algorithm is very efficient in mapping protein-substrate binding interactions, allowing the exploration of the entire surface^53^. PELE’s global exploration clearly identified two binding poses for the pentapeptide (Fig. 6D). After pose refinement (blue dots in Fig. 6D) we find protein-peptide interaction energies below -150 kcal mol^-1^, corresponding to strong binding interactions; take into account that these correspond to interaction energy snapshots with an all-atom force field energies and implicit solvent, not reflecting real free energy values. The x axis describes the distance between the catalytic Cys145 residue and the terminal D-Ala residue. Thus, while PELE finds a potential catalytic minimum for this terminal residue, corresponding to x axis values around 4 Å, it is not the preferred binding pose at this level of theory.

Due to the approximate nature of PELE’s calculations, we performed more accurate molecular dynamics (MD) simulations using full explicit solvent. For each of the two binding poses we performed two 450 ns MD simulations, aiming at probing the stability of the substrate in the predicted binding mode. For the catalytic minima, one replica escaped the binding site while the second one kept the binding pose (Fig S15). For the second minimum (with best PELE’s interaction energies), one replica also escaped the binding site. Interestingly, the second replica quickly readapted itself in the binding site, adopting a terminal D-Ala catalytic pose for the remainder of the simulation (Fig. 6F). An illustrative catalytic pose conformation is also shown (Fig. 6E).

This modeling analysis inferred that RumI_c_1 acts as a D-D-carboxypeptidase (DD-CPase) that removes the terminal D-Alanine from the stem peptide. DD-CPases are widespread in bacteria and are pivotal for cell shape maintenance or PG repair^54–57^. In the pneumococcus, the DD-CPase PBP3 (*dacA*) regulates the amount of pentapeptides available for crosslinking, thereby influencing PG structure, the division site positioning, and the cell shape^57–62^. To test whether RumI_c_1 could suppress the defects of the Δ*dacA* mutant, we examined the growth profile of Δ*dacA* cells expressing either *luc*, *rumI_c_1^WT^* or *rumI_c_1^C145A^* under the control of the P*_lac_* promoter. The Δ*dacA P_lac_-luc* strain exhibited a slight growth delay and reached a lower optical density than the wild-type DacA+ *P_lac_-luc* strain (Fig. 7A). While Δ*dacA* P_lac_-*rumI_c_1^C145A^* cells showed a similar growth profile to that of Δ*dacA* P_lac_-*luc* cells, expression of *rumI_c_1^WT^* rescued both the growth defect and the exacerbated autolysis rate of the Δ*dacA* mutant. We next analyzed the effect of RumI_c_1 on Δ*dacA* cell morphology. Since the absence of DD-CPases increases incorporation of fluorescent D-amino acids into PG^36,63^, we also analyzed how RumI_c_1 modulates HADA incorporation in Δ*dacA* cells. As previously reported^57,58,60^, Δ*dacA* cells exhibit severe morphological defects, with a significant increase in cell width and a marked increase in HADA signal (Fig. 7B-D). By contrast, Δ*dacA* cells expressing RumI_c_1, but not RumI_c_1^C145A^, restored a near wild-type ovoid-shape, with significantly reduced cell width and HADA incorporation (Fig. 7B and C). Notably, expression in wild-type *dacA*+ cells also led to a reduction in HADA incorporation (Fig. S16). Taken together, these findings demonstrate that RumI_c_1 acts by trimming the stem peptide, conferring protection against RumC1.

**Figure 7:**
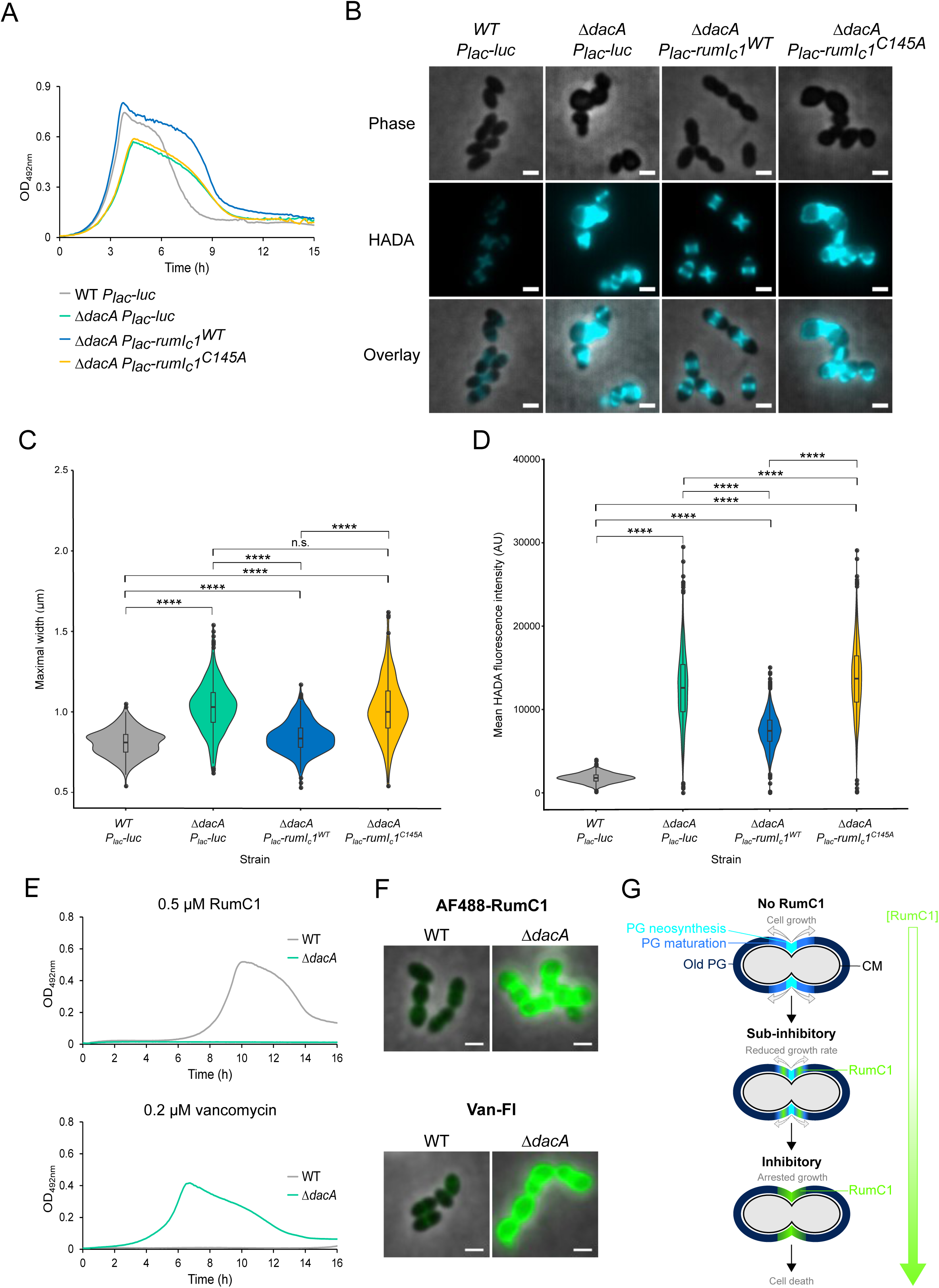
The immunity protein RumI_c_1 partially replaces the D-D-carboxypeptidase DacA. (A) Growth of *WT P_lac_-luc* (R5339), *ΔdacA P_lac_-luc* (R5342), *ΔdacA P_lac_-rumI_c_1^WT^* and *ΔdacA P_lac_-rumI_c_1^C145A^* (R5344) strains. Precultures and cultures were performed in the presence of 50 µM IPTG. Data shown are representative of three independent replicates. (B) Morphology and HADA incorporation in *WT P_lac_-luc* (R5339), *ΔdacA P_lac_-luc* (R5342), *ΔdacA P_lac_-rumI_c_1^WT^* (R5343) and *ΔdacA P_lac_-rumI_c_1^C145A^* (R5344) cells. Phase contrast, fluorescent (false-colored in light blue), and false-colored overlay images are shown. Cells were incubated for 10 min with HADA. Precultures and cultures were performed in the presence of 50 µM IPTG. Scale bar, 1 µm. Images are representative of three independent replicates. (C) Violin plots representing the distribution of the maximal width of *WT P_lac_-luc* (R5339) (grey, n=377), *ΔdacA P_lac_-luc* (R5342) (green, n=431), *ΔdacA P_lac_-rumI_c_1^WT^* (R5343) (blue, n=400) and *ΔdacA P_lac_-rumI_c_1^C145A^* (R5344) (yellow, n=397) cells. Boxes extend from the 25^th^ percentile to the 75^th^ percentile, with the horizontal line at the median. Dots represent outliers. Statistical analysis was performed using the U test of Mann-Whitney (n.s., non-significant, p-value > 0,05; ****, p-value < 0,0001). (D) Violin plots representing the mean HADA fluorescence intensity in *WT P_lac_-luc* (R5339) (grey, n=1685), *ΔdacA P_lac_-luc* (R5342) (green, n=1107), *ΔdacA P_lac_-rumI_c_1^WT^* (R5343) (blue, n=2083) and *ΔdacA P_lac_-rumI_c_1^C145A^* (R5344) (yellow, n=1162) cells. Boxes extend from the 25^th^ percentile to the 75^th^ percentile, with the horizontal line at the median. Dots represent outliers. Statistical analysis was performed using unpaired t-test (****, p-value < 0,0001). (E) Growth of wild-type (R1501) and Δ*dacA* (R5207) strains in the presence of 0.5 µM RumC1 (upper panel) and 0.2 µM vancomycin (lower panel). Data shown are representative of three independent experiments, and all resistance profiles are presented in Fig S10. (F) Overlay between phase-contrast and fluorescence of wild-type (R1501) and Δ*dacA* (R5207) cells stained with 2.5 µM AF488-RumC1 (upper panel) or 0.2 µM BODIPY^®^ FL vancomycin (Van-Fl) (lower panel). Scale bars, 1 µm. Images are representative of two independent replicates. (G) Scheme of the dose-dependent binding pattern of RumC1 and its inhibitory effect on the growth of pneumococcal cells. In the pneumococcus, the neonascent PG (light blue) is inserted at mid-cell and then matured (blue) and pushed towards the cell poles (dark blue) as the cell grows. When added at sub-inhibitory concentrations, RumC1 binds preferentially to an intermediate of PG maturation, thereby slowing down PG synthesis and cell growth. At inhibitory concentrations, RumC1 accumulates at mid-cell and causes a complete halt in PG synthesis and cell growth, while even higher concentrations of RumC1 lead to cell death.

The PG of Δ*dacA* cells is enriched in pentapeptide^58^, making them more sensitive to RumC1 and exhibiting increased AF488-RumC1 surface accumulation (Fig. 7E-G and S17). While the deletion of *dacA* also led to an increase in vancomycin-BODIBY labelling, this accumulation correlated with reduced sensitivity to vancomycin. This paradoxical effect likely arises because elevated pentapeptide levels in the Δ*dacA* cell wall titrate and reduce the amount of free vancomycin, thereby reducing its inhibitory effect on PG cross-linking^64,65^. This mode of protection by increasing the density of untrimmed pentapeptide in the cell wall is ineffective against RumC1. Altogether, these results highlight the terminal D-Ala-D-Ala dipeptide of the stem peptide as pivotal for PG targeting by both RumC1 and vancomycin. However, their mechanisms of interaction with PG synthesis appear distinct.

## Discussion

In this work, we unveiled the mode of action of the RumC1 bacteriocin, produced by the human gut symbiont *R. gnavus* E1 and characterized by its unique structural fold and lack of known functional signature. By combining genetics, protein biochemistry, structural modeling and single-cell imaging in the sensitive pathogen *S. pneumoniae*, we uncovered a novel mechanism of cell wall poisoning by RumC1. Our findings collectively demonstrate that RumC1 represents a new category of cell wall toxin, distinct from previously characterized bacteriocins and antibiotics.

First, RumC1 inhibits transpeptidase activity of the cell, thereby blocking PG synthesis and cell growth, and inducing the LiaFSR stress response, which senses PG damage. This dual effect suggests that RumC1 disrupts PG integrity in a manner recognized by the cell’s envelope stress surveillance systems. Second, RumC1 selectively binds to nascent PG, an interaction unprecedented among bacteriocins. This binding is dose-dependent, transitioning from bacteriostatic to bactericidal effects as RumC1 accumulates in the PG. Third, we showed that *S. pneumoniae* can evolve weak resistance to RumC1 by downregulating the WalRK two-component system, a master regulator of PG homeostasis, essential for coordinating cell wall synthesis with growth and division. Finally, we identified the RumI_c_1 protein, encoded and annotated in the RumC1 BGC as a potential immunity protein, to be sufficient for counteracting RumC1 toxicity in *S. pneumoniae*. RumI_c_1 functions by cleaving the terminal D-Ala of the PG stem peptide, a critical residue for cross-linking nascent PG chains. Furthermore, RumI_c_1 also confers cross-resistance to vancomycin, further underscoring its role in modulating PG structure and integrity. However, two key findings demonstrate that RumC1 interaction with the PG differs from that of vancomycin: the purified stem peptide titrates vancomycin but does not alter RumC1 toxicity; accumulation of the full-length pentapeptide in the PG of the *ΔdacA* mutant reduced vancomycin toxicity, while enhancing RumC1 activity. In all, these findings demonstrate that RumC1 binds to PG and interferes with its synthesis *via* a unique, concentration-dependent mechanism (Fig. 7G): at low doses, RumC1 preferentially targets a premature PG component, resulting in a slower PG synthesis and cell growth. At high doses, RumC1 accumulates at mid-cell where nascent PG is inserted, causing a complete halt in PG synthesis and ultimately leading to cell death.

RumC1 fundamentally differs from all bacteriocins previously characterized as cell-envelope toxins (including all sactipeptides that have already been characterized)^66,67^, which typically act by disrupting membrane integrity (i.e. through pore formation or depolarization) or by inhibiting one of the multiple sequential steps driving the synthesis of the lipid II precursor of PG assembly^66–71^. Unlike these bacteriocins, RumC1 does not compromise membrane integrity and does not depolarize it^13^, but instead targets late stages of PG metabolism. This particular property aligns RumC1 with only three other natural AMPs - vancomycin, corbomycin and complestatin - all glycopeptides originating from actinomycetes. Corbomycin and complestatin interfere with PG maturation by blocking access to hydrolases required for remodeling the PG during cell growth and division^72^. Similarly, RumC1 appears to obstruct PG maturation, as inferred from *walRK* mutations in resistant strains, which downregulate PG hydrolases essential for coordinating cell wall dynamics. As for RumC1, however, the molecular anchor of corbomycin and complestatin in the PG remains to be precisely identified. RumC1 exhibits key distinctions from these glycopeptides. While corbomycin and complestatin are bacteriostatic, RumC1 shows both bacteriostatic and bactericidal activity depending on its concentration. In addition, RumC1’s interaction with PG is highly dependent on the terminal D-Ala-D-Ala dipeptide of the stem peptide, a key determinant for its binding and toxicity. The protective mechanism mediated by RumI_C_1, which trims this stem peptide, further highlights its importance for the interaction of RumC1 with the PG. Intriguingly, inactivation of the DD-CPase PBP3, which increases the density of full length pentapeptides at the cell surface, enhances RumC1 toxicity and accumulation in the cell-wall. This effect contrasts sharply with vancomycin, whose toxicity is reduced in the *ΔdacA* mutant, highlighting a fundamental difference in their mode of interaction with PG. Furthermore, vancomycin-resistant strains (i.e. Vancomycin Resistant Enterococcus; VRE), which modify the terminal dipeptide to D-Lac or a D-Ser, remain fully sensitive to RumC1^13,14^, reinforcing the idea that RumC1’s interaction with the PG is independent of the nature of the terminal end of the pentapeptide. This property is strongly supportive for an interaction of RumC1 with an early crosslinked PG intermediate in the cell wall, as directly visualized on purified sacculi prepared from growing cells (Fig. 2C).

The low level of resistance observed in *walRK* mutants or through *rumI_c_1* expression suggests that RumC1 does not target a specific enzyme but rather a structural intermediate within the PG mesh which cannot directly accumulate suppressing mutations – a component that is highly conserved across bacterial species despite variations in PG synthetic and remodeling enzymes. This conservation likely underpins RumC1’s broad spectrum activity against diverse bacteria, including multidrug resistant pathogens. Future studies should explore RumC1’s effect in species with distinct PG metabolic frameworks, which may reveal further insights into its target specificity and potential evolutionary constraints.

In conclusion, RumC1 represents a breakthrough in our understanding of bacteriocin-mediated PG disruption. By targeting a late PG intermediate, RumC1 operates through a mechanism distinct from all known antibiotics. In addition, its lack of toxicity to human cells and efficacy in complex natural environments (e.g., blocking the *ex vivo* colonization of hen’s gut contents by *C. perfringens* with minimal alteration of the microbiota^73^), positions it as a promising candidate for antimicrobial development. As such, RumC1 is one of these rare biomolecules, defined as ‘sophisticated natural products’^9^ – a class of biomolecules prioritized for their innovative mode of action. Future research should focus on elucidating the precise molecular anchor of RumC1 within the PG mesh and exploring its therapeutic applications across diverse bacterial pathogens, with a particular focus on MDR species.

## Materials and Methods

### Strains and growth media

All *S. pneumoniae* strains used were derived from R800 strain (ref) and are listed in Table S1. All pneumococcal strains were rendered unable to spontaneously develop competence either by deletion of the *comC* gene (*comC0*) (ref) or by combining the non-compatible alleles *comC_2_* and *comD_1_* (ref). Stock cultures were routinely grown in C + Y medium (ref) and were kept frozen at -70 °C after addition of 15% glycerol (vol/vol). Standard procedures for transformation were used^74^. Briefly, pre-competent cells were treated at 37°C for 10 min with synthetic 100 ng mL^-1^ CSP1 to induce competence and exposed to transforming DNA for 20 min at 30°C. Transformants were then plated in CAT-agar supplemented with 4% horse blood and incubated for 2h at 37°C. Transformants were then selected by addition of a second layer of agar medium containing the appropriate antibiotic and incubated overnight at 37°C. Antibiotic concentrations used for selection were: kanamycin, 250 µg mL^-1^; spectinomycin, 100 µg mL^-1^ and erythromycin 0.1 µg mL^-1^. Unless otherwise described, pre-competent cultures were prepared by growing cells to an OD_550nm_ of 0.1 in C + Y medium. Cells were then 10-fold concentrated in C + Y medium supplemented with 15% glycerol and stored at -70°C.

### Heterologous expression and purification of RumC1

Two synthetic plasmids containing *E. coli* codon-optimized genes of *R. gnavus* E1 were obtained from Genscript: pET-15b-*rumMc1* encodes the maturation enzyme RumMc1 and pETM-40-*rumC1* allows the expression of a Maltose Binding Protein tagged-RumC1 (MBP-RumC1) with a TEV protease site between the MBP tag and the RumC1 sequence. pET-15b-*rumMc1*, pETM-40-*rumC1* as well as pSuf plasmid containing *sufABCDSE* genes were used to transform competent *E. coli* BL21 (DE3) cells for expression. The resulting strain was grown in M9 medium containing kanamycin (50 µg mL^-1^), ampicillin (100 µg mL_-1_), chloramphenicol (34 µg mL^-1^), vitamin B1 (0.5 µg mL^-1^), FeCl_3_ (50 µM), MgSO_4_ (1 mM) and glucose (4 mg mL^-1^). The culture was done in a bioreactor (Inceltech LH.SGi, Discovery 100) of 12 L, at 37°C with moderate aeration (140 L h^-1^) and stirring (90 rpm). At an OD_600_ of 0.6, FeCl_3_ (50 µM) and L-cysteine (300 µM) were added and the culture was induced using 1 mM IPTG. The temperature was reduced to 25°C and the cells were grown for 15 h under gentle aeration and stirring. Cells were then harvested by centrifugation (4,000 rpm for 20 min at 4°C). Cell pellet was resuspended in 150 mL of buffer A (50 mM Tris, pH 8, 50 mM NaCl) supplemented with three tablets of a protease inhibitor cocktail (cOmpleteTM, EDTA-free Protease inhibitor cocktail tablets, Roche). Cell pellet was then sonicated, and the lysate was clarified by centrifugation at 40,000 rpm for 30 min at 4°C. The supernatant was collected and passed over a Dextrin Sepharose High Performance column equilibrated with buffer A coupled to an FPLC (ÄKTA Pure 25, Cytiva). MBP-RumC1 was eluted with buffer B (50 mM Tris, pH 8, 50 mM NaCl, 40 mM Maltose). Fractions containing MBP-RumC1 were pooled and concentrated in a 10,000 MWCO Amicon® Ultra centrifugal filter device. The sample was digested by TEV protease with a final TEV:MBP-RumC1 ratio of 1:100 (w/w) for 30 min at room temperature. MBP-tag, TEV protease and RumC1 were separated by loading over a HiLoad 26/600 Superdex® 75 prep grade column (Cytiva) equilibrated in buffer C (HEPES 50 mM, NaCl 100 mM, pH 7.5). The peptide concentration was estimated by UV-visible spectroscopy on a Cary 50 UV-Vis Spectrophotometer (Varian) by using an extinction coefficient at 280 nm of 8,480 M^-1^.cm^-1^. Mature RumC1 was treated with Trypsin (TPCK, Sigma) for 1 h at 37°C, at a w/w ratio of 50:1 for RumC1:trypsin, to remove the leader peptide. RumC1 mature peptide free of the leader peptide was then purified by using a C18 preparative column (Phenomenex, Kinetex, 5 μm, 100Å; 250 x 10 mm). The column was equilibrated with buffer D (90% H_2_O, 10% acetonitrile, 0.1% trifluoroacetic acid) and subjected to a gradient from 0% to 20% of buffer E (90% acetonitrile, 10% H_2_O, 0.1% trifluoroacetic acid) in 35 min on a ÄKTA Pure 25. The fractions containing pure RumC1, determined by mass spectrometry (ESI-MS) analysis^13^, were then lyophilized and stored at -20°C.

### Selection of RumC1-resistant mutants

RumC1-resistant mutants were selected using the library of mutagenic PCR fragments as described for antibiotics streptomycin and rifampicin used for calibration (see supplementary Information) except that transformants were plated in 500 µL of CAT-agar medium supplemented with 4% horse blood and 1x solid MIC RumC1 (3.5 µM) in 24-well plates.

### Effect of antimicrobial compounds on *S. pneumoniae* growth and MIC determination

To determine the MIC of antimicrobial compounds and to monitor their effect on growth at low cell density (OD_550_=0.005; ≈7.10^6^ CFU/mL), *S. pneumoniae* cells were grown in C + Y medium until mid-log phase (OD_550_= 0.3). When needed, 50 µM IPTG or 100 ng mL^-1^ ATC were added to induce expression of genes under the control of the P_lac_ or P_tet_ promoters, respectively. Cells were then diluted to a final OD_550_=0.005 in a 96-well plate containing 300 µL of C + Y medium containing serial dilutions of antimicrobial compounds (with IPTG or ATC when required). Growth (OD_492_) was monitored by a Varioskan Flash (Thermo 399 Electron Corporation) microplate reader at 37°C without agitation. OD_492_ values were recorded every 5 min for 12 h to 16h. Liquid MIC was defined as the lowest concentration of antimicrobial compound that inhibited cell growth after 16 h of incubation at 37°C. For determination of MIC of Daptomycin, cells were grown in presence of 0.5 mM CaCl_2_.

To monitor the effect of RumC1 and Vancomycin on growth at high cell density (OD_550_=0.1; ≈2.10^8^ CFU/mL), pre-cultures of strain R1501 grown in C + Y medium to an OD_550_ of 0.3 were diluted to an OD_550_ of 0.005 in C + Y medium in a 96-well plate (290 µL per well) and incubated at 37°C in a Varioskan Flash microplate reader until OD_550_=0.1. Ten microliters of 3X-concentrated solutions of RumC1 or Vancomycin were then added to each well and cells were incubated at 37°C for a further 14h.

To determine the solid MIC of RumC1 used for selection of RumC1-resistant mutants, mock transformations were plate on CAT-agar medium supplemented with 4% horse blood and containing serial dilutions of RumC1. The solid MIC was defined as the lowest concentration of RumC1 that inhibited colony formation after 30 h of incubation at 37°C.

### Survival assays with RumC1 and vancomycin

Pre-cultures of strain R1501 grown in C + Y medium to an OD_550_ of 0.3 were diluted to an OD_550_ of 0.005 in C + Y medium and incubated at 37°C. When OD_550_ reached 0.1, cells were treated with RumC1 or vancomycin and incubated at 37°C. At 10, 30, 60 and 120 min, 100 μL of each cell suspension were collected, serially diluted and plated on CAT-agar medium supplemented with 4% horse blood. After 24 h of incubation at 37°C, colonies were counted to determine the CFU/mL.

### Sacculi preparation

Deproteinated sacculi were prepared from *S. pneumoniae* R1501 mid-log growing cells as previously described^75^, with some modifications. Briefly, 2 L of culture grown in THY medium to an OD_550nm_ of 0.3 were harvested and resuspended in 40 mL of ice-cooled PBS. Cell suspension was then poured dropwised into 160 mL of boiling 5% SDS and boiled for another 30 min. Sacculi were then centrifugated at 40,000 x g for 30 min at room temperature and washed 5 times with 50 mL of water and 2 times with 50 mL of 1M NaCl prior to incubation with 10 μg mL^-1^ DNase and 50 μg mL^-1^ RNase in 100 mM Tris-HCl pH 7.5, 20 mM MgSO_4_ for 2 h at 37°C on a rotating wheel. 10 mM CaCl_2_ and 100 μg mL^-1^ trypsin were then added and sacculi were incubated overnight at 37°C on a rotating wheel. Sacculi were then incubated in 1% SDS for 15 min at 80°C, harvested by centrifugation (40,000 x g, 30 min, RT) and resuspended in 20 mL of 8 M LiCl. Sacculi were submitted to a second washing step with 25 mL 10 mM EDTA pH 7 followed by a third washing step with 25 mL of water. The final pellet was resuspended in 1 mL of water and stored at 4°C.

### Protoplast preparation

To generate *S. pneumoniae* protoplasts, 20 mL of a pneumococcal preculture grown in C + Y medium at 37°C to an OD_550_=0.1 was centrifugated (2900 x g, 10 min), washed with 20 mL of resuspension buffer (100 mM Tris-HCl pH 7.6, 1 mM MgCl_2_ and 3 mM β-mercaptoethanol) and centrifugated a second time. The pellet was resuspended in 2 mL of protoplastization buffer (100 mM Tris-HCl pH 7.6, 1 mM MgCl_2_, 3 mM β-mercaptoethanol and 1 M Sucrose) and incubated at 37°C for 10 min to 1 h. Throughout the incubation, aliquots of 100 µL of cell suspension were diluted into 1 mL H_2_0 and the OD_550_ of the resulting suspension was measured. Protoplastization is considered complete when OD_550_≤0.01. Protoplast suspension was diluted at 1:10 with protoplastization buffer before further utilization.

### Fluorescent labelling of RumC1

Alexa Fluor 488 5-SDP ester (AF488, Thermo Fisher Scientific) resuspended in 50% DMSO was mixed with RumC1 (WT or A12E) in 500 µL of 100 mM NaHCO_3_, pH 8.3. The concentrations of AF488 and RumC1 are 2 mg mL^-1^. The coupling was performed at room temperature for 1h under gentle stirring in the dark. The conjugated AF488-RumC1 was then purified by using a C18 preparative column (Phenomenex, Kinetex, 5 μm, 100Å; 250 x 10 mm) with a detection at 280 and 495 nm. The column was equilibrated with buffer A (90% H_2_O, 10% acetonitrile, 0.1% trifluoroacetic acid) and subjected to a gradient from 0% to 20% of buffer B (90% acetonitrile, 10% H_2_O, 0.1% trifluoroacetic acid) in 52 min on a ÄKTA Pure 25. The fractions containing pure conjugated AF488-RumC1, determined by mass spectrometry (ESI-MS) analysis, were then lyophilized and stored at -20°C.

### Labelling of *S. pneumoniae* with AF488-RumC1 and BODIPY^®^ FL vancomycin

For staining of log-phase cultures of *S. pneumoniae* with fluorescent antibiotics, pre-cultures grown in C + Y medium to an OD_550_ of 0.3 were diluted to an OD_550_ of 0.005 in C + Y medium and incubated at 37°C until early log phase (OD_550_=0.1). When needed, IPTG (50 µM) or ATC (100 ng mL^-1^) were added to both the pre-culture and the culture. One hundred and fifty microliters of cells were then incubated with AF488-RumC1, AF488-RumC1-A12E or BODIPY^®^ FL vancomycin at the indicated concentration (Thermo Fisher Scientific, USA) for 10 min at 37°C, washed three times with cold pre-C medium by centrifugation (3000 x g, 3 min) and resuspended in 10 µL of cold pre-C medium. Two microliters of this suspension were spotted on a microscope slide containing a slab of 1.2% agarose diluted in pre-C medium as previously described^76^ and observed using a Nikon ECLIPSE Ti microscope. Images were captured and analysed with NIS-element AR software (Nikon).

For staining of peptidoglycan, a sacculi suspension was centrifugated and diluted at 1:50 in pre-C medium. 50 µL of this suspension were incubated with 2.5 µM AF488-RumC1 for 10 min at 37°C before washing with pre-C medium and imaging.

For staining of protoplasts, 150 µL of a protoplast suspension in protoplastization buffer was incubated with 2.5 µM AF488-RumC1 for 10 min at 37°C, washed three times with protoplastization buffer and resuspended in 10 µL of protoplastization buffer before imaging. A control of AF488-RumC1 binding to exponentially growing cells was carried out in parallel.

For image analysis, cells were detected with the threshold command of NIS-element software. The AF488-RumC1 and BODIPY^®^ FL vancomycin fluorescence of each cell was then measured using the automatic mean FITC measurement option. For each image, the background fluorescence of the agar slide was measured from the average fluorescence of 5 different cell-free regions. This average background fluorescence was then subtracted from the fluorescence measured for each cell of the image.

### Peptidoglycan labeling with Fluorescent D-amino acids

For all peptidoglycan labelling experiments, pneumococcal pre-cultures grown in C + Y medium to an OD_550_ of 0.3 were diluted to an OD_550_ of 0.005 in C + Y medium and incubated at 37°C. When needed, 50 µM IPTG or 100 ng mL^-1^ ATC were added to induce expression of genes under the control of the P_lac_ or P_tet_ promoters, respectively. When cultures reached early log phase (OD_550_=0.1), cells were treated with 70 mM fluorescent D-amino acid Hydroxycoumarincarbonylamino-D-alanine (HADA)^77^ for 10 min at 37°C except for experiments with the RumC1-resistant strains shown in Fig. 1D in which HADA labeling was for 15 min. Cells were then washed three times with cold pre-C medium by centrifugation at 4°C and imaging. To measure the effect of RumC1 and vancomycin on peptidoglycan incorporation, cells were co-incubated with 70 mM HADA and RumC1 or vancomycin at the indicated concentration for 10 or 30 min at 37°C before washing and imaging. For co-labelling experiments with AF488-RumC1, cells were treated with 70 mM HADA and 2.5 µM AF488-RumC1 for 10 min at 37°C before washing and imaging.

### Heterologous expression and purification of RumI_c_1_S_^WT^ and RumI_c_1_S_^C145A^

To produce and purify the extracellular domain of RumI_c_1_S_^WT^ or RumI_c_1_S_^C145A^ with a 6His tag at the C-terminus, the sequences coding for the predicted extracellular domain (E53 to F280) of RumI_c_1 WT and C145A were synthesized by GeneScript USA with codons optimized for *Escherichia coli* and cloned into the pET-21(a)+ expression vector. The resulting plasmids (pET21-rumI_c_1^WT^ and pET21-rumI_c_1^C145A^) were transformed into *E. coli* BL21-Gold (DE3) cells. The resulting strains were then grown aerobically in LB medium at 37 °C until OD_600_ reaches 0.7 and protein overproduction was induced with 1 mM IPTG for 3 h at 37°C. Cells were then harvested by centrifugation, resuspended in buffer A (0.1 M Tris-HCl pH 8, 0.5 M NaCl and 2.5 mM DTT) and disrupted by sonication. Lysates were centrifuged 20 min at 39,000 × *g* and the supernatants were loaded onto a HisTrap FF column (GE Healthcare). Elution was performed with a gradient of Imidazole ranging from 0 to 500 mM in buffer A and *proteins were* eluted at 50 mM Imidazole. The eluted fractions were then loaded onto a HiLoad 16/60 Superdex 200 column (GE healthcare) equilibrated with buffer A. Proteins were stored at −80 °C after addition of 10 % glycerol and freezing in liquid nitrogen.

### PG Pentapeptide cleavage assays by RumI_c_1_S_^WT^ and RumI_c_1_S_^C145A^

To assess the cleavage of the PG Pentapeptide by RumI_c_1_S_^WT^ and RumI_c_1_S_^C145A^, an indirect assay of the effect of RumI_c_1_S_^WT^ and RumI_c_1_S_^C145A^ on the antagonization of Vancomycin activity by the PG Pentapeptide was performed (see SI). A 120 µM solution of an unamidated version of the pneumococcal PG Pentapeptide (L-Ala-D-Glu-L-Lys-D-Ala-D-Ala, GenScript, USA) diluted in cleavage buffer (20 mM Tris pH 7.4, 5 mM MgCl_2_ and 2.5 mM DTT) was pre-incubated at 37°C for 1h30 with His6-tagged RumI_c_1_S_^WT^ or RumI_c_1_S_^C145A^ at a PG Pentapeptide:RumI_c_1_S_ molar ratio of 20:1. Twenty microliters of these reaction solutions were then mixed with 10 µL of a 6 µM Vancomycin solution diluted in cleavage buffer or with 10 µL of cleavage buffer. After 1h30 of incubation at 37°C, the reaction mixtures were added to individual wells of a 96-well plate (30 µL per well). Two hundred and seventy microliters of a cell suspension of strain R1501 (*comC0*) grown to OD_550_=0.3 and diluted to an OD_550_ of 0.005 in C + Y medium were then added to each well and growth (OD_492_) was monitored by a Varioskan Flash (Thermo 399 Electron Corporation) microplate reader at 37°C without agitation. OD_492_ values were recorded every 5 min for 16h.

### Modeling and simulation of RumI_c_1 interaction with the pentapeptide

We used AlphaFold2 Multimer v2.3.0^78^ to predict the structure of RumI_c_1 in complex with the pentapeptide. Among the predicted 20 models, we selected the top-ranked structure in which the peptide was positioned within the catalytic cleft. Because AlphaFold2 generates only L-amino acids, the resulting complex did not reproduce the correct stereochemistry of the peptidoglycan substrate. Thus, taking into account the catalytic site proposed by AlphaFold2, we performed molecular docking using Glide (version 2023-1)^79^ to generate 10 binding poses of the peptidoglycan within the catalytic cleft, incorporating the correct stereochemistry and including the muramic acid and N-acetylglucosamine moieties.

The 10 docked poses were used as starting structures for Monte Carlo sampling with PELE^80^, employing the default induced-fit protocol implemented in the PELE platform. Each pose was simulated for ∼15 hours using 24 computing cores on the MareNostrum 4 supercomputer. From the resulting energy landscapes, we identified the two main energy minima. The lowest-energy pose from each minimum was selected for further refinement using PELE with 60 CPUs for ∼2 hours.

The lowest-energy structures from the two refined minima were subsequently subjected to unbiased MD simulations using AdaptivePELE^81^. This framework automates ligand parametrization with AmberTools (v18)^82^ and applies the ff99SB force field^83^, followed by MD simulations performed with the OpenMM engine^84^. For each selected structure, two independent replicas were carried out. Systems were equilibrated for 200 ps under NVT conditions and 500 ps under NPT conditions prior to ∼450 450 ns of production simulation, corresponding to 48 GPU hours (with minor variations in the final trajectory length).

### Sequence analysis and sequence alignments

Identification of putative protein domains was performed with InterPro^85^ and prediction of transmembrane helices was carried out using the TMHMM online software (https://services.healthtech.dtu.dk/services/TMHMM-2.0/). Sequence alignments were generated with the Clustal Omega program (European Molecular Biology Laboratory) with default parameters.

### Statistical analyses

All statistical analyses were performed with GraphPad Prism 10 (GraphPad Software, LLC). Pairwise comparison between two conditions were done with a Mann–Whitney *U* test or with an unpaired t-test. *P*-values are displayed as follows: ****, *P* < 0.0001; ***, 0.0001 < *P* < 0.001; **, 0.001 < *P* < 0.01; *, 0.01 < *P* < 0.05; ns, *P* > 0.05. Note that GraphPad Prism computes an exact *P*-value when the size of the smallest sample is less than or equal to 100, and approximates the *P*-value from a Gaussian approximation when the samples are large.

## Acknowledgments

We thank Isabelle Mortier-Barrière and André Zapun for their experimental assistance and for their helpful discussion. We thank O. Glushonkov and J.-P. Kleman for support on the M4D fluorescence microscopy platform. This work was funded by grants from the Agence Nationale de la Recherche (ANR-20-CE44-0021 to VD, ML and PP; ANR-23-CE11-0029 to CM), by the Centre National de la Recherche Scientifique, Université de Toulouse and by a predoctoral fellowship from the Spanish Ministry of Science and Innovation to A.M.D-R (FPU21/03921). This work used the platforms of the Grenoble Instruct-ERIC center (ISBG; UAR 3518 CNRS-CEA-UGA-EMBL) within the Grenoble Partnership for Structural Biology (PSB), supported by FRISBI (ANR-10-INBS-0005-02) and Labex GRAL and ARCANE, financed within the University Grenoble Alpes graduate school (Ecoles Universitaires de Recherche) CBH-EUR-GS (ANR-17-EURE-0003).

