## Extended data for "The broad-spectrum RumC1 bacteriocin targets a transient peptidoglycan intermediate of the nascent cell wall"

### Supplementary information

#### Strains construction

All primers used for strain construction are listed in Table S2. The $rumI_{c}1$, $rumI_{c}2$, $rumE_{c}$, $rumY_{c}$, $rumF_{c}$ and $rumG_{c}$ genes were synthesized with codons optimized for *S. pneumoniae* by Integrated DNA Technologies (IDT, USA).

Two platforms were used to induce gene-expression. The CEP-$P_{lac}$ platform allows the expression of a gene of interest under the control of an IPTG-inducible promoter and it is inserted at the *ami* locus. The CEPII-$P_{tet}$ platform allows the expression of a gene under the control of an ATC-inducible promoter and it is inserted at the *cpsN* locus.

To generate strain R5172 (*comC2D1, CEP-*$P_{lac}$*-luc, CEPII-*$P_{tet}$*-luc*), the regions upstream and downstream of *dprA-mturquoise* in the CEPII-$P_{tet}$ platform were amplified from strain R4850 (laboratory stock) using primer pairs CJ662-CJ964 and OCN592-CJ667 respectively, and the $P_{tet}$*-luc* fragment was amplified from R4830 (laboratory stock) using primer pair CJ965-OCN591. SOE PCR using these three fragments and primer pair CJ662-CJ667 generated a *CEPII-*$P_{tet}$*-luc* DNA fragment which was transformed into R5339, with transformants selected with erythromycin.

To generate strain R5177 (*comC2D1, CEP-*$P_{lac}$*-*$rumI_{c}1$*-*$rumI_{c}2$*, CEPII-*$P_{tet}$*-*$rumE_{C}Y_{C}F_{C}G_{C}$), the regions upstream and downstream of *luc* in the CEPII-$P_{tet}$ platform were amplified from strain R5172 using primer pairs CJ662-OCN596 and OCN599-CJ667 respectively, and the $rumE_{C}Y_{C}F_{C}G_{C}$ genes were amplified from R5341 using primer pair OCN597-OCN598. SOE PCR using these three fragments and primer pair CJ662-CJ667 generated a *CEPII-*$P_{tet}$*-*$rumE_{C}Y_{C}F_{C}G_{C}$ DNA fragment which was transformed into R5230, with transformants selected with erythromycin.

To generate the R5207 strain (*comC0, dacA::spc*), a *dacA::spc* DNA fragment was created by initial amplification of the regions upstream and downstream of the *dacA* (*spr0776*) gene using primer pairs OCN512-OCN513 and OCN516-OCN517 and R1501 DNA as template. The *aad9* gene conferring spectinomycin resistance was amplified using the primer pair OCN514-OCN515 and plasmid DNA pR412[1] as template. SOE PCR on these three fragments with the primer pair OCN512-OCN517 generated a DNA fragment with the *dacA* gene replaced with the spectinomycin resistance cassette, which was transformed into R1501, with transformants selected with spectinomycin. To generate the R5208 strain (*comC0, dacB::trim*), a *dacB::trim* DNA fragment was created by initial amplification of the regions upstream and downstream of the *dacB* (*spr0554*) gene using primer pairs AB129-AB130 and AB133-AB134 and R1501 DNA as template. The *trim* resistance cassette was amplified using the primer pair AB131-AB132 and strain TK108[2] as template. SOE PCR on these three fragments with the primer pair AB129-AB134 generated a DNA fragment with the *dacB* gene replaced with the *trim* resistance cassette, which was transformed into R1501, with transformants selected with trimethoprim.

To generate strain R5230 (*comC2D1, CEP-$P_{lac}$-rumI$_{c}$1-rumI$_{c}$2*), the regions upstream and downstream of *dprA* in the CEP-$P_{lac}$ platform were amplified from R3831[3] using primer pairs MM43-AB2 and AB4-OEC11 respectively, *rumI$_{c}$1 and rumI$_{c}$2* were amplified from a codon-optimized sequence synthesized by IDT using primer pair AB1-AB3. SOE PCR using these three fragments and primer pair MM43-OEC111 generated a *CEP-$P_{lac}$-rumI$_{c}$1-rumI$_{c}$2* DNA fragment which was transformed without selection into R3831 with a 4h phenotypic expression phase in liquid culture. Positive clones were determined by PCR and sequencing (Eurofins genomics). To generate strain R5231 (*comC2D1, CEP-$P_{lac}$-rumI$_{c}$1*), the regions upstream and downstream of *rumI$_{c}$1-rumI$_{c}$2* in the CEP-*$P_{lac}$-rumI$_{c}$1-rumI$_{c}$2* platform were amplified from R5230 using primer pairs MM43-AB21 and AB22-OEC11 respectively. SOE PCR using these two fragments and primer pair MM43-OEC111 generated a *CEP-$P_{lac}$-rumI$_{c}$1* DNA fragment which was transformed without selection into R5230 with a 4h phenotypic expression phase in liquid culture. Positive clones were determined by PCR and sequencing.

To generate strain R5232 (*comC2D1, CEP-$P_{lac}$-rumI$_{c}$1, CEPII-$P_{tet}$-rumI$_{c}$1*), the regions upstream and downstream of *rumE$_{C}$Y$_{C}$F$_{C}$G$_{C}$* in the CEPII-$P_{tet}$ platform were amplified from R5177 using primer pairs CJ662-AB135 and AB138-CJ667 respectively, *rumI$_{c}$1* was amplified from a codon-optimized sequence synthesized by IDT using primer pair AB136-AB137. SOE PCR using these three fragments and primer pair CJ662-CJ667 generated a *CEPII-$P_{tet}$-rumI$_{c}$1* DNA fragment which was transformed into R5231, with transformants selected with erythromycin.

To generate strain R5336 (*comC2D1, CEP-$P_{lac}$-rumI$_{c}$1$^{C145A}$)*, two PCR reactions, with primer pairs AB40-AB42 and AB43-AB41, were used to generate 2 fragments that incorporate a mutant primer (AB42) at one extremity of the first fragment and its complement (AB43) at the other, overlapping, extremity of the second fragment. SOE PCR using these two fragments and primer pair AB40-AB41 generated a *CEP-$P_{lac}$-rumI$_{c}$1$^{C145A}$* DNA fragment with the mutant sequence in the middle. This fragment was transformed into R5152 and transformants were screened by PCR sequencing for insertion of the point mutation. Strains R5337 (*comC2D1, CEP-$P_{lac}$-rumI$_{c}^{H234A}$*) and R5338 (*comC2D1, CEP-$P_{lac}$-rumI$_{c}^{D250A}$*) were generated in the same way using primer pairs AB18-AB44 and AB45-AB41 for R5337 and AB18-AB46 and AB47-AB41 for R5338 for the first PCR and primer pair AB18-AB41 for the SOE PCR of both strains. To generate strain R5339 (*comC2D1, CEP-$P_{lac}$-luc)*, R3369[3] cells were transformed with genomic DNA from strain R3310[3] (*comC0, CEP-$P_{lac}$-luc*) and transformants were selected with kanamycin.

To generate strain R5340 (*comC2D1, CEP-$P_{lac}$-rumI$_{c}$2*), the regions upstream and downstream of *rumI$_{c}$1-rumI$_{c}$2* in the CEP-*$P_{lac}$-rumI$_{c}$1-rumI$_{c}$2* platform were amplified from

R5230 using primer pairs MM43-AB23 and AB24-OEC111 respectively. SOE PCR using these two fragments and primer pair MM43-OEC111 generated a *CEP-P*$_{lac}$*-rumI*$_{c}$*2* DNA fragment which was transformed without selection into R5230 with a 4h phenotypic expression phase in liquid culture. Positive clones were determined by PCR and sequencing.

To generate strain R5341 (*comC2D1, CEP-P*$_{lac}$*-rumE*$_{C}$*Y*$_{C}$*F*$_{C}$*G*$_{C}$), the regions upstream and downstream of *dprA* in the CEP-*P*$_{lac}$ platform were amplified from R3831[3] using primer pairs MM43-AB48 and AB9-OEC111 respectively, *rumE*$_{c}$ *and rumY*$_{c}$ were amplified from a codon-optimized sequence synthesized by IDT using primer pair AB49-AB8. SOE PCR using these three fragments and primer pair MM43-OEC111 generated a *CEP-P*$_{lac}$*-rumE*$_{C}$*Y*$_{C}$ DNA fragment which was transformed without selection into R3831 with a 4h phenotypic expression phase in liquid culture. Positive clones were determined by PCR and sequencing. The regions upstream and downstream of *rumE*$_{C}$*Y*$_{C}$ in the CEP-*P*$_{lac}$*-rumE*$_{C}$*Y*$_{C}$ platform were then amplified from the resulting strain (*comC2D1, CEP-P*$_{lac}$*-rumE*$_{C}$*Y*$_{C}$) using primer pairs AB40-AB10 and AB13-OEC111 respectively, and *rumF*$_{c}$ *and rumG*$_{c}$ were amplified from a codon-optimized sequence synthesized by IDT using primer pair AB11-AB12. SOE PCR using these three fragments and primer pair AB40-OEC111 generated a *CEP-P*$_{lac}$*-rumE*$_{C}$*Y*$_{C}$*F*$_{C}$*G*$_{C}$ DNA fragment which was transformed without selection into strain *comC2D1, CEP-P*$_{lac}$*-rumE*$_{C}$*Y*$_{C}$ with a 4h phenotypic expression phase in liquid culture. Positive clones were determined by PCR and sequencing.

To generate strains R5342 (*comC2D1, CEP-P*$_{lac}$*-luc, dacA::spc)*, R5343 (*comC2D1, CEP-P*$_{lac}$*-rumI*$_{c}$*1, dacA::spc)* and R5344 (*comC2D1, CEP-P*$_{lac}$*-rumI*$_{c}$*1*$^{C145A}$*, dacA::spc)*, R5339, R5231 and R5336 cells, respectively, were transformed with genomic DNA from strain R5207 (*comC0, dacA::spc*) and transformants were selected with spectinomycin.

**Generation of the library of mutagenic PCR fragments**

PCR primers were designed using Vector NTI software (InforMax, North Bethesda, Md.) with a target length of 25 to 30 nucleotides, a PCR product size of 4.5 kb and a desired melting temperature of 60 to 65°C. Default settings were used for all other parameters. The primers sequence can be found in Table S3. PCR primers were synthesized and purified with a standard salt-free purification process by Eurofins Genomics (Ebersberg, Germany). Error-prone PCR reactions were carried out in the presence of 25 µM $MnCl_2$ using the low-fidelity DreamTaq DNA Polymerase (Thermo Fisher scientific) and *S. pneumoniae* R800 genomic DNA as a template. Cycling conditions for the reaction were as follows: 95°C for 3 min, followed by 40 cycles of amplification (95°C for 30 s, 55°C for 30 s and 72°C for 4 min) and by a final elongation step at 72°C for 5 min 30 s. The size of the amplification products was confirmed by electrophoresis on agarose gels. Of the 505 primer pairs of the library, 88 pairs did not produce amplification products and were thus manually redesigned. All of these new primer pairs except one gave the correct PCR product when retested. Two primer pairs generating two 2 kb amplicons were designed to amplify the remaining amplicon (fragment O, pool No. 16). To create the amplicon pools, 10 µL of each individual PCR product were combined so that pools No. 1 to 20 contained 23 to 25 consecutive amplicons and pool No. 21 contained 10 consecutives amplicons. The exact composition of the amplicon pools is listed in Table S3. Amplicons and amplicon pools were stored at -20°C.

**Calibration of the NT-based mutagenesis method with streptomycin and rifampicin**

To screen for streptomycin or rifampicin-resistant mutants, 100 µL aliquots of pre-competent cells of strain R1818 (*comC0, hexAΔ3::ermAM*) were resuspended in 900 µL C+Y medium with 100 ng mL$^{-1}$ CSP and incubated at 37°C for 8 min. 3 µL of amplicon pools were then added to a 100 µL aliquot of this culture, followed by incubation at 30°C for 20 min. 1 mL of fresh C+Y medium was then added into each tube before incubation at 37°C for 1 hr 30 min. Cells were then diluted and plated on 10 mL CAT agar with 4% horse blood containing 100 µg mL$^{-1}$ streptomycin or 2 µg mL$^{-1}$ rifampicin. After 24 h of incubation at 37°C, colonies were counted to determine the CFU. The frequency of antibiotic-resistant mutants was calculated by dividing the number of CFU obtained on agar plates containing antibiotic by the number of CFU obtained antibiotic-free plates. A second round of transformation was performed with each of the individual amplicons comprising the pools that gave positive results in the first round.

**Effect of RumI$_c$1 on tolerance to RumC1**

Strains R5339 and R5231 were grown in C + Y medium containing 50 µM IPTG to an OD$_{550}$ of 0.3 and diluted to an OD$_{550}$ of 0.005 in C + Y medium containing 50 µM IPTG. Cells were then treated with 0, 1.5 or 3 µM RumC1 and incubated at 37°C. After 60 min, 100 µL of each cell suspension were collected, serially diluted and plated on CAT-agar medium supplemented with 4% horse blood. After 24 h of incubation at 37°C, colonies were counted to determine the CFU/mL. Survival percentages were determined by comparing CFU of cells exposed to RumC1 with those of non-exposed cells.

**Protection of *S. pneumoniae* and *R. gnavus* E1 against RumC1 by RumI$_c$1**

To assess the protective effect of RumI$_c$1$_S^{WT}$ on *S. pneumoniae* and *R. gnavus* E1 during exposure to RumC1, MICs were determined by broth microdilution in microtiter plates, with experiments performed in three independent biological replicates. Briefly, a bacterial suspension of *S. pneumoniae* R1501 in MH broth supplemented with 5% lysed horse blood and 20 mg L$^{-1}$ β-nicotinamide adenine dinucleotide, or *R. gnavus* E1 in BHI-YH-Cys were used. Two hundred microliters of cell suspension containing 400 µM of RumI$_c$1$_S^{WT}$ were dispensed into the last row of a sterile flat-bottom polypropylene 96-well microplate. The remaining wells were filled with 100 µL of bacterial suspension in culture medium. Two-fold serial dilutions of RumI$_C$1$_S^{WT}$ were then performed across the rows using 100 µL transfers to obtain decreasing concentrations in culture medium. Microplates were pre-incubated at 37°C during 150 min under anaerobic conditions for *R. gnavus* E1, while 90 min under aerobic atmosphere for *S. pneumoniae*. Subsequently, 100 µL of bacterial suspension containing RumC1 at 50 µM (*R. gnavus* E1) or 25 µM (*S. pneumoniae*) were added to the wells of the last column. Two-fold serial dilutions were then performed across the columns to generate decreasing concentrations of RumC1. No dilutions were performed in the last row or the last column, which served to assess the effects of RumI$_C$1$_S^{WT}$ or RumC1 alone, respectively. Microplates were incubated under anaerobic conditions at 37°C for 48 h for *R. gnavus* E1 or 24 h under aerobic conditions for *S. pneumoniae*. The MIC was defined as the lowest concentration of RumC1 that inhibited visible bacterial growth in the presence of RumI$_C$1$_S^{WT}$. Sterility and growth controls were included in each assay.

**Determination of RumC1 and AF488-RumC1 oligomerization state**

The oligomerization state of RumC1 and AF488-RumC1 in solution was evaluated by loading over a Superdex® Peptide 10/300 GL column (Cytiva) equilibrated with HEPES 50 mM, NaCl 100 mM, pH 7.5 on AKTA Pure 25. The calibration of the column was performed by using Ribonuclease A (MW 13.7 kDa), Aprotinin (MW 6.5 kDa) and pF5 (MW 1.6 kDa) as standard samples. The observed elution volumes were 9.38, 10.95 and 14.4 mL for Ribonuclease A, Aprotinin and pF5, respectively. Under these conditions, the elution volumes for RumC1 and AF488-RumC1 were 12.30 mL, which correspond approximately to an apparent MW of 4.3 kDa for both sequences.

**Antagonization of Vancomycin and RumC1 activity by the pneumococcal PG Pentapeptide**

For the antagonism assays with the pneumococcal PG Pentapeptide, Vancomycin was used at a MIC (0.2 µM) final concentration and RumC1 was used at sub-MIC (0.4 µM) and MIC (0.6 µM) final concentrations. The unamidated pneumococcal PG Pentapeptide (L-Ala-D-Glu-L-Lys-D-Ala-D-Ala) was synthesized by GenScript, USA and resuspended in water. 30X-concentrated solutions of RumC1 and Vancomycin diluted in 50 mM HEPES pH 7.5 were incubated with the PG Pentapeptide at Vancomycin:PG Pentapeptide molar ratios of 0:1000, 1:0, 1:25, 1:50, 1:100, 1:250 and 1:500 and at a RumC1:PG Pentapeptide molar ratio of 1:1000. After 1h30 of incubation at 37°C, 10 µL of each mixture were added to individual wells of a 96-well plate. Two hundred and ninety microliters of a cell suspension of strain R1501 (*comC0*) grown to $OD_{550}$=0.3 and diluted to an $OD_{550}$ of 0.005 in C + Y medium were then added to each well and growth ($OD_{492}$) was monitored by a Varioskan Flash (Thermo 399 Electron Corporation) microplate reader at 37°C without agitation. $OD_{492}$ values were recorded every 5 min for 16h.

**RumC1 and Vancomycin cleavage assays by RumI$_c$1$_S$**

RumC1 and Vancomycin cleavage by His6-tagged RumI$_c$1$_S$ were assessed indirectly by the effect of RumI$_c$1 pre-treatment on the toxicity of RumC1 and Vancomycin to pneumococcal cells. RumC1 and Vancomycin were used at sub-MIC (0.5 µM and 0.07 µM, respectively) and MIC (0.6 µM and 0.2 µM, respectively) final concentrations. 30X-concentrated solutions of RumC1 and Vancomycin diluted in cleavage buffer (20 mM Tris pH 7.4, 5 mM $MgCl_2$ and 2.5 mM DTT) were treated with His6-tagged RumI$_c$1$_S$ at an antibiotic:RumI$_c$1 molar ratio of 1:1. After 3 h of incubation at 37°C, 10 µL of each mixture were added to individual wells of a 96-well plate. Two hundred and ninety microliters of a cell suspension of strain R1501 (*comC0*) grown to $OD_{550}$=0.3 and diluted to an $OD_{550}$ of 0.005 in C + Y medium were then added to each well and growth ($OD_{492}$) was monitored by a Varioskan Flash (Thermo 399 Electron Corporation) microplate reader at 37°C without agitation. $OD_{492}$ values were recorded every 5 min for 16h. In parallel of each experiment, a pentapeptide cleavage assay by RumI$_c$1 was performed as a control for RumI$_c$1 activity.

**Table S1: Pneumococcal strains used in this study**

| Strain | Genotype / Relevant features | References |
|---|---|---|
| R800 | *Wild-type* (R6 derivative) | [4] |
| R391 | *comA::kan,* Kan$^R$ | Laboratory stock |
| R1114 | *comA::kan, lytB::ery,* Kan$^R$, Ery$^R$ | Laboratory stock |
| R1501 | *comC0* | [5] |
| R1818 | *comC0, hexAΔ3::ermAM*, Ery$^R$ | [6] |
| R5172 | *comC2D1, CEP-P$_{lac}$-luc, CEPII-P$_{tet}$-luc,* Kan$^R$, Ery$^R$ | This study |
| R5177 | *comC2D1, CEP-P$_{lac}$-rumI$_c$1-rumI$_c$2, CEPII-P$_{tet}$-rumE$_C$Y$_C$F$_C$G$_C$,* Kan$^R$, Ery$^R$ | This study |
| R5207 | *comC0, dacA::spc*, Spc$^R$ | This study |
| R5208 | *comC0, dacB::trim*, Trim$^R$ | This study |
| R5230 | *comC2D1, CEP-P$_{lac}$-rumI$_c$1-rumI$_c$2,* Kan$^R$ | This study |
| R5231 | *comC2D1, CEP-P$_{lac}$-rumI$_c$1,* Kan$^R$ | This study |
| R5232 | *comC2D1, CEP-P$_{lac}$-rumI$_c$1, CEPII-P$_{tet}$-rumI$_c$1,* Kan$^R$, Ery$^R$ | This study |
| R5336 | *comC2D1, CEP-P$_{lac}$-rumI$_c$1$^{C145A}$,* Kan$^R$ | This study |
| R5337 | *comC2D1, CEP-P$_{lac}$-rumI$_c$1$^{H234A}$,* Kan$^R$ | This study |
| R5338 | *comC2D1, CEP-P$_{lac}$-rumI$_c$1$^{D250A}$,* Kan$^R$ | This study |
| R5339 | *comC2D1, CEP-P$_{lac}$-luc,* Kan$^R$ | This study |
| R5340 | *comC2D1, CEP-P$_{lac}$ -rumI$_c$2,* Kan$^R$ | This study |
| R5341 | *comC2D1, CEP-P$_{lac}$-rumE$_C$Y$_C$F$_C$G$_C$,* Kan$^R$ | This study |
| R5342 | *comC2D1, CEP-P$_{lac}$-luc, dacA::spc,* Kan$^R$, Spc$^R$ | This study |
| R5343 | *comC2D1, CEP-P$_{lac}$-rumI$_c$1, dacA::spc,* Kan$^R$, Spc$^R$ | This study |
| R5344 | *comC2D1, CEP-P$_{lac}$-rumI$_c$1$^{C145A}$, dacA::spc,* Kan$^R$, Spc$^R$ | This study |

**Table S2: Plasmids used in this study**

| Plasmid | Genotype | References |
|---|---|---|
| pET21-rumI$_c$1$^{WT}$ | Sequence coding for the extracellular domain (E53 to F280) of RumI$_c$1 cloned into pET-21a(+), Amp$^R$ | This study |
| pET21-rumI$_c$1$^{C145A}$ | Sequence coding for the extracellular domain (E53 to F280) of RumI$_c$1 with the point mutation C145A cloned into pET-21a(+), Amp$^R$ | This study |

**Table S3: Primers used in this study**

*rum$_{Immunity}$* refers to *rumI$_c$1*, *rumI$_c$2* or *rumE$_C$Y$_C$F$_C$G$_C$* genes.

| Primer | Sequence (5’-3’) | Use |
|---|---|---|
| AB1 | ATTAAGCTTAAGGAGGTGTACATATGGTCAGAAAGAGCCGCAAAAAGAGA | Generation of *CEP-P$_{lac}$-rumI$_c$1-rumI$_c$2* |
| AB2 | CTTTTTGCGGCTCTTTCTGACCATATGTACACCTCCTTAAGCTTAATTGT | Generation of *CEP-P$_{lac}$-rumI$_c$1-rumI$_c$2* |
| AB3 | CCATTAAAAATCAAACGGATCCTTTATCTCCTCCTACTACTTTTGGAAACGA | Generation of *CEP-P$_{lac}$-rumI$_c$1-rumI$_c$2* |

| | | |
|---|---|---|
| AB4 | CGTTTCCAAAAGTAGTAGGAGGAGATAAAGGATCCGTTTGATTTTTAATGGATAATG | Generation of $CEP\text{-}P_{lac}\text{-}rumI_{c}1\text{-}rumI_{c}2$ |
| AB8 | CCATTAAAAATCAAACGGATCCATTATCGCCTCCTAATCAATCCACATTTTCA | Generation of $CEPII\text{-}P_{lac}\text{-}rumE_{C}Y_{C}F_{C}G_{C}$ |
| AB9 | GAAAATGTGGATTGATTAGGAGGCGATAATGGATCCGTTTGATTTTTAATG | Generation of $CEPII\text{-}P_{lac}\text{-}rumE_{C}Y_{C}F_{C}G_{C}$ |
| AB10 | CTTTACTTCTAGAACCAC ATTATCGCCTCCTAATCAATCCACATTTTCA | Generation of $CEPII\text{-}P_{lac}\text{-}rumE_{C}Y_{C}F_{C}G_{C}$ |
| AB11 | TGAAAATGTGGATTGATTAGGAGGCGATAATGTGGTTCTAGAAGTAAAG | Generation of $CEPII\text{-}P_{lac}\text{-}rumE_{C}Y_{C}F_{C}G_{C}$ |
| AB12 | CCATTAAAAATCAAACGGATCCTTTTCTTTCCTCCTAACACTTTTTCCAA | Generation of $CEPII\text{-}P_{lac}\text{-}rumE_{C}Y_{C}F_{C}G_{C}$ |
| AB13 | GGAAAAAGTGTTAGGAGGAAAGAAAAGGATCCGTTTGATTTTTAATGG | Generation of $CEPII\text{-}P_{lac}\text{-}rumE_{C}Y_{C}F_{C}G_{C}$ |
| AB18 | CACCAGCATATTCAGCAACATC | Generation of $CEP\text{-}P_{lac}\text{-}rumI_{c}1^{H234A}$ and $rumI_{c}1^{D250A}$ |
| AB21 | TCCATTAAAAATCAAACGGATCCTTAAAACAACGTAGTAAAAGACCAT | Generation of $CEP\text{-}P_{lac}\text{-}rumI_{c}1$ |
| AB22 | ATGGTCTTTTACTACGTTGTTTTAAGGATCCGTTTGATTTTTAATGGA | Generation of $CEP\text{-}P_{lac}\text{-}rumI_{c}1$ |
| AB23 | TAACCATAGCTATCTTCTTCTTCATATGTACACCTCCTTAAGCTTAATTG | Generation of $CEP\text{-}P_{lac}\text{-}rumI_{c}2$ |
| AB24 | CAATTAAGCTTAAGGAGGTGTACATATGAAGAAGAAGATAGCTATGGTTA | Generation of $CEP\text{-}P_{lac}\text{-}rumI_{c}2$ |
| AB40 | GAACAGCAGCTTTACAAGCTTCAAC | Generation of $CEP\text{-}P_{lac}\text{-}rumE_{C}Y_{C}F_{C}G_{C}$ *and* $CEP\text{-}P_{lac}\text{-}rumI_{c}1^{C145A}$ |
| AB41 | CAGGCTTGATCCCCAGTAAGTC | Generation of $CEP\text{-}P_{lac}\text{-}rumI_{c}1^{C145A}/rumI_{c}1^{H234A}$ and $rumI_{c}1^{D250A}$ |
| AB42 | GAATGCTAGCAGTCAACGGTGCTGGGCCGACAGCGTTGAGCAT | Generation of $CEP\text{-}P_{lac}\text{-}rumI_{c}1^{C145A}$ |
| AB43 | ATGCTCAACGCTGTCGGCCCAGCACCGTTGACTGCTAGCATTC | Generation of $CEP\text{-}P_{lac}\text{-}rumI_{c}1^{C145A}$ |
| AB44 | GTGACTTTACCACTACGGGTGCTTTCATAGTTCTAACGGGGAT | Generation of $CEP\text{-}P_{lac}\text{-}rumI_{c}1^{H234A}$ |
| AB45 | ATCCCCGTTAGAACTATGAAAGCACCCGTAGTGGTAAAGTCAC | Generation of $CEP\text{-}P_{lac}\text{-}rumI_{c}1^{H234A}$ |
| AB46 | ACGGGAAGATATGTGTACACGCTCCAAATTCAAAAAAGCGAAG | Generation of $CEP\text{-}P_{lac}\text{-}rumI_{c}1^{D250A}$ |
| AB47 | CTTCGCTTTTTTGAATTTGGAGCGTGTACACATATCTTCCCGT | Generation of $CEP\text{-}P_{lac}\text{-}rumI_{c}1^{D250A}$ |
| AB48 | CACTGCACGTTTGGCACTGCCCAAATGTACACCTCCTTAAGCTTAATTGT | Generation of $CEP\text{-}P_{lac}\text{-}rumE_{C}Y_{C}F_{C}G_{C}$ |
| AB49 | ATTAAGCTTAAGGAGGTGTACATTTGGGCAGTGCCAAACGTGCAGTG | Generation of $CEP\text{-}P_{lac}\text{-}rumE_{C}Y_{C}F_{C}G_{C}$ |
| AB102 | TCTCAACCAATTGCAGAAGCTCCA | RT-qPCR – *spr1875* |
| AB103 | TGTTTCGGCCGTTTGCTCCT | RT-qPCR – *spr1875* |
| AB104 | CGTAGTGCTCAAACAAATGGAGCC | RT-qPCR - *pcsB* |
| AB105 | CACTCATTGCAGCAACACGTGAA | RT-qPCR - *pcsB* |
| AB106 | TTGTGGGTGAGGCGTCCAAC | RT-qPCR - *liaR* |
| AB107 | GCTAAGGTCGCGTCAATCCCA | RT-qPCR - *liaR* |
| AB108 | GTTGCAAACTTTGGACTTGGCGTG | RT-qPCR - *pspC* |
| AB109 | ATGGCTTGGGCTTCTTTGCGT | RT-qPCR - *pspC* |
| AB135 | CCTCCTTAAGCTTAATTGTTACTCTATCAATGATAGAGTTATTATAC | Generation of $CEPII\text{-}P_{tet}\text{-}rumI_{c}1$ |
| AB136 | ACTCTATCATTGATAGAGTAACAATTAAGCTTAAGGAGGTGTAC | Generation of $CEPII\text{-}P_{tet}\text{-}rumI_{c}1$ |
| AB137 | AAGCTTCTCGAGGGTACCTTAAAACAACGTAGTAAAAGACCATAAATTG | Generation of $CEPII\text{-}P_{tet}\text{-}rumI_{c}1$ |
| AB138 | CTTTTACTACGTTGTTTTAAGGTACCCTCGAGAAGCTTAACAAAG | Generation of $CEPII\text{-}P_{tet}\text{-}rumI_{c}1$ |
| CJ662 | GCGGGTGAGGATTTTATTGGTGACG | Generation of $CEPII\text{-}P_{tet}\text{-}rumE_{C}Y_{C}F_{C}G_{C}$ and $CEPII\text{-}P_{tet}\text{-}rumI_{c}1$ |
| CJ667 | GCGAACTCACGGATAGTTCCTAATCT | Generation of $CEPII\text{-}P_{tet}\text{-}rumE_{C}Y_{C}F_{C}G_{C}$ and $CEPII\text{-}P_{tet}\text{-}rumI_{c}1$ |
| CJ964 | TAATAATAAGGATCGGAGACCACCCGTCGACCGGCCGTCTAG | Generation of $CEPII\text{-}P_{tet}\text{-}luc$ |
| CJ965 | CTAGACGGCCGGTCGACGGGTGGTCTCCGATCCTTATTATTA | Generation of $CEPII\text{-}P_{tet}\text{-}luc$ |
| MB | | RT-qPCR - *rpoD* |

| | | |
|---|---|---|
| MB | | RT-qPCR - *rpoD* |
| MM43 | CAATGCTTACTTGTTCTGGATTCCATTG | Generation of *CEP-P*$_{lac}$*-rum*$_{immunity}$ |
| OEC111 | CCACCAGCATCCGCAATCA | Generation of *CEP-P*$_{lac}$*-rum*$_{immunity}$ |
| OCN512 | GGCAGCTAGAGCGTAAGAAGTG | Generation of *dacA::spc* |
| OCN513 | GTATTCAAATATATCCTCCTCACCTTGAGCAACAGCAGTAGAAACAC | Generation of *dacA::spc* |
| OCN514 | GTTTCTACTGCTGTTGCTCAAGGTGAGGAGGATATATTTGAATACATACG | Generation of *dacA::spc* |
| OCN515 | GCGGACAAACTGATTCCACCAATTTTTTTAATCTGTTATTTAAATAGTTT | Generation of *dacA::spc* |
| OCN516 | TAAATAACAGATTAAAAAAATTGGTGGAATCAGTTTGTCCGCT | Generation of *dacA::spc* |
| OCN517 | GGCAGTGGTAGCAGACCATTCGA | Generation of *dacA::spc* |
| OCN591 | GTTAAGCTTCTCGAGGGGTACCTTACAATTTGGGCTTTCCGCCCT | Generation of *CEPII-P*$_{tet}$*-luc* |
| OCN592 | GGGCGGAAAGCCCAAATTGTAAGGTACCCTCGAGAAGCTTAAC | Generation of *CEPII-P*$_{tet}$*-luc* |
| OCN596 | CTGCACGTTTGGCACTGCCCATTATTTTTCCTCCTTATTTATTTAGATCTACTCTATC | Generation of *CEPII-P*$_{tet}$*-rum*$E_C Y_C F_C G_C$ |
| OCN597 | GATAGAGTAGATCTAAATAAATAAGGAGGAAAAATAATGGGCAGTGCCAAACGTGCAG | Generation of *CEPII-P*$_{tet}$*-rum*$E_C Y_C F_C G_C$ |
| OCN598 | GTTAAGCTTCTCGAGGGTACCCAACTCAGCTCATCTCAGAAAGGATC | Generation of *CEPII-P*$_{tet}$*-rum*$E_C Y_C F_C G_C$ |
| OCN599 | GATCCTTTCTGAGATGAGCTGAGTTGGGTACCCTCGAGAAGCTTAAC | Generation of *CEPII-P*$_{tet}$*-rum*$E_C Y_C F_C G_C$ |

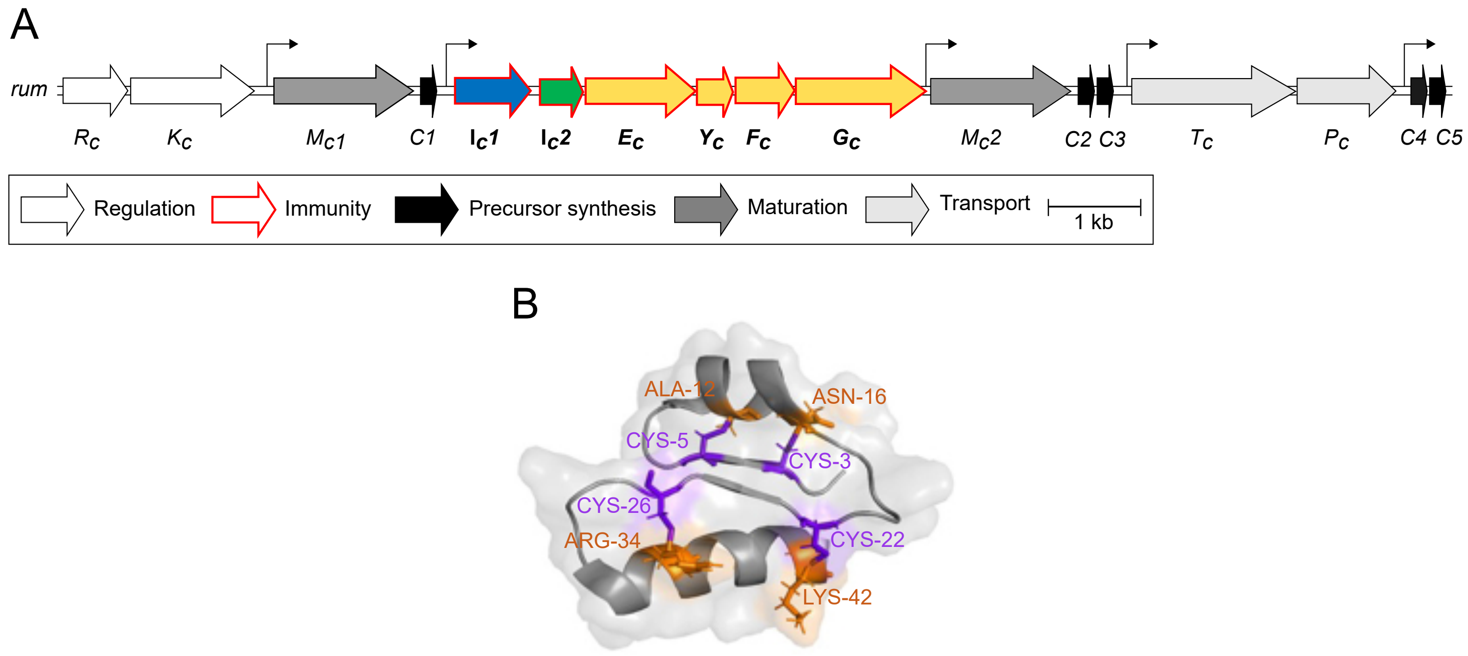
A
rum
Rc
Kc
Mc1
C1
Ic1
Ic2
Ec
Yc
Fc
Gc
Mc2
C2 C3
Tc
Pc
C4 C5
Regulation
Immunity
Precursor synthesis
Maturation
Transport
1 kb
B
ALA-12
ASN-16
CYS-5
CYS-3
CYS-26
ARG-34
CYS-22
LYS-42

Figure S1

**Figure S1: Biosynthesis and structure of RumC1.**

(A) Biosynthetic gene cluster of RumC1. (B) 3D structure of RumC1. Cysteine residues and the residues partners of the sactionine bonds are colored in purple and orange, respectively (PDB: 6T33).

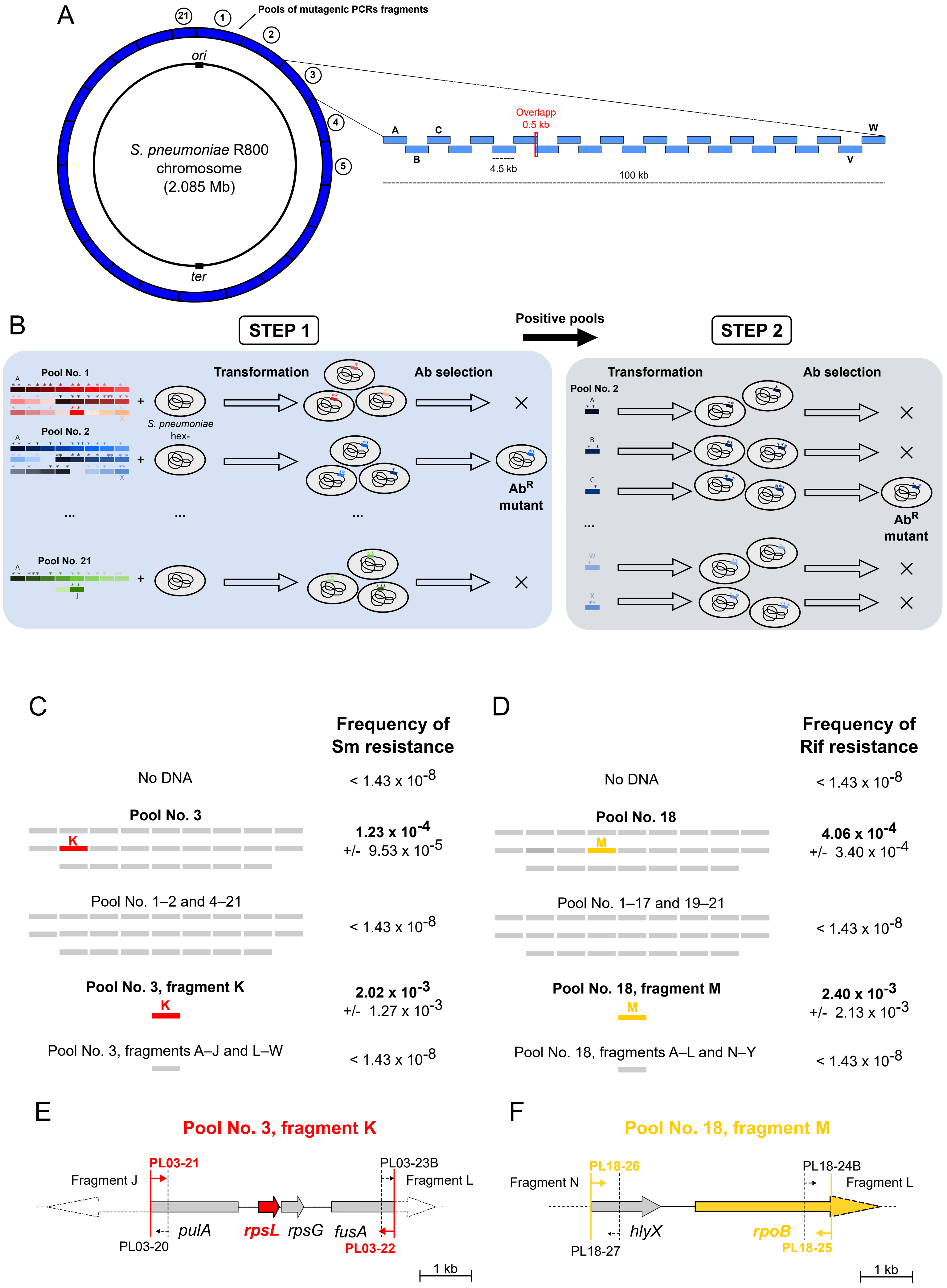
A
Pools of mutagenic PCRs fragments
ori
S. pneumoniae R800
chromosome
(2.085 Mb)
ter
Overlapp
0.5 kb
4.5 kb
100 kb
B
STEP 1
Positive pools
STEP 2
Pool No. 1
Transformation
Ab selection
S. pneumoniae
hex-
Pool No. 2
Ab^R
mutant
Pool No. 21
C
Frequency of
Sm resistance
No DNA
< 1.43 x 10^-8
Pool No. 3
1.23 x 10^-4
+/- 9.53 x 10^-5
Pool No. 1–2 and 4–21
< 1.43 x 10^-8
Pool No. 3, fragment K
2.02 x 10^-3
+/- 1.27 x 10^-3
Pool No. 3, fragments A–J and L–W
< 1.43 x 10^-8
D
Frequency of
Rif resistance
No DNA
< 1.43 x 10^-8
Pool No. 18
4.06 x 10^-4
+/- 3.40 x 10^-4
Pool No. 1–17 and 19–21
< 1.43 x 10^-8
Pool No. 18, fragment M
2.40 x 10^-3
+/- 2.13 x 10^-3
Pool No. 18, fragments A–L and N–Y
< 1.43 x 10^-8
E
Pool No. 3, fragment K
PL03-21
PL03-23B
Fragment J
Fragment L
pulA
rpsL
rpsG
fusA
PL03-20
PL03-22
1 kb
F
Pool No. 18, fragment M
PL18-26
PL18-24B
Fragment N
Fragment L
hlyX
rpoB
PL18-27
PL18-25
1 kb

Figure S2

**Figure S2: Description and calibration of the NT-based mutagenesis method with streptomycin and rifampicin.**

(A) Schematic representation of the library of mutagenic PCRs fragments used to identify antibiotic-resistant mutants. (B) Schematic representation of the two-step NT-based mutagenesis method used to identify antibiotic-resistant mutants. In the first step, *S. pneumoniae* R1818 (*comC0, hexAΔ3::ermAM*) competent cells were transformed with each of the 21 independent pools of mutagenic amplicons constituting the library and then selected on antibiotic-containing plates. A second transformation and antibiotic selection step was performed with individual amplicons of positive pool(s) identified during step 1. Asterisks represent point mutations. (C) Frequency of streptomycin-resistant mutants obtained by transformation with the library of mutagenic PCR fragments. Competent cells of strain R1818 (*comC0, hexAΔ3::ermAM*) were transformed with each of the 21 amplicon pools of the library and were then submitted to a second round of transformation with individual amplicons from pool No. 3. Means and standard deviations obtained from four independent experiments are presented. (D) Frequency of rifampicin-resistant mutants obtained by transformation with the library of mutagenic PCR fragments. Competent cells of strain R1818 (*comC0, hexAΔ3::ermAM*) were transformed with each of the 21 amplicon pools of the library and were then submitted to a second round of transformation with individual amplicons from pool No. 18. Means and standard deviations obtained from four independent experiments are presented. (E) Schematic representation of the PCR fragment K of pool No. 3 containing the *rpsL* gene (right panel). Position of primers PL03-21 and PL03-22 used to amplify the PCR fragment K is indicated. The reverse primer used to amplify the PCR fragment J (PL03-20) and the forward primer used to amplify the PCR fragment L (PL03-23B) are also represented. (F) Schematic representation of the PCR fragment M of pool No. 18 containing the *rpoB* gene (right panel). Position of primers PL18-26 and PL18-25 used to amplify the PCR fragment K is indicated. The reverse primer used to amplify the PCR fragment N (PL18-27) and the forward primer used to amplify the PCR fragment L (PL18-24B) are also represented.

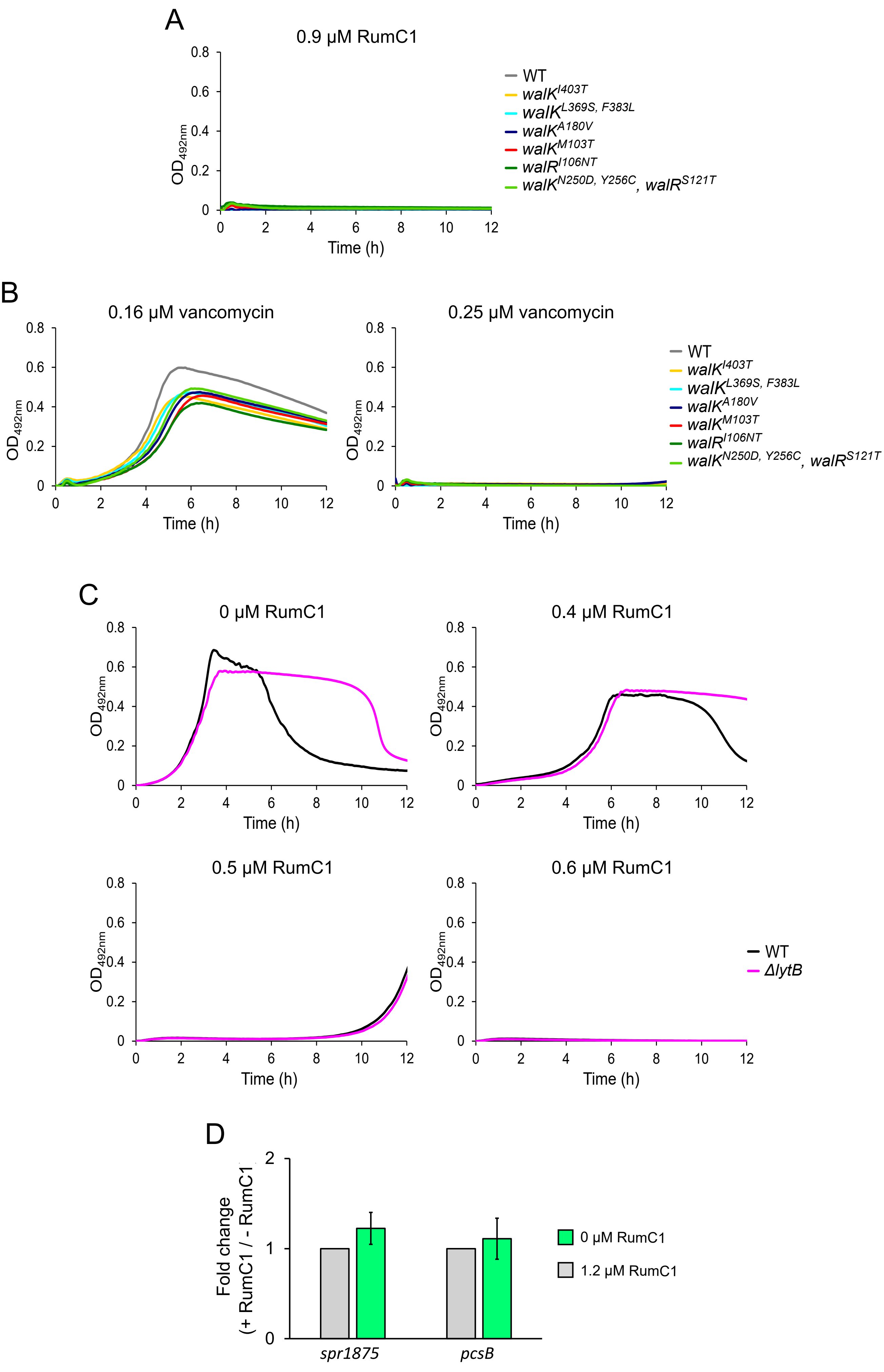

A
0.9 μM RumC1
OD492nm
Time (h)
WT
walKI403T
walKL369S, F383L
walKA180V
walKM103T
walRI106NT
walKN250D, Y256C, walRS121T
B
0.16 μM vancomycin
0.25 μM vancomycin
C
0 μM RumC1
0.4 μM RumC1
0.5 μM RumC1
0.6 μM RumC1
ΔlytB
D
Fold change
(+ RumC1 / - RumC1)
0 μM RumC1
1.2 μM RumC1
spr1875
pcsB

Figure S3

**Figure S3: RumC1 and vancomycin resistance of the RumC1-resistant mutants, effect of LytB on RumC1 resistance and effect of RumC1 on the expression level of WalR-regulated genes.**

(A) Growth of wild-type (strain R3584) and RumC1-resistant strains (strains R4884, R4885, R4886, R4887, R4888 and R4889) in the presence of 0.9 µM RumC1 (related to Fig. 1C). Data shown here are representative of two independent experiments. (B) Growth of wild-type (strain R3584) and RumC1-resistant strains (strains R4884, R4885, R4886, R4887, R4888 and R4889) in the presence of sub-MIC (0.16 µM) and MIC (0.2 µM) concentrations of vancomycin. The vancomycin concentrations are indicated on top of the graphs. Data shown here are representative of two independent experiments. (C) Growth of wild-type (R391) and Δ*lytB* (R1114) strains in the presence of increasing concentrations of RumC1. The RumC1 concentrations are indicated on top of the graphs. Data shown here are representative of three independent experiments. (D) Effect on RumC1 on the expression level of two WalR-regulated genes (*pcsB* and *spr1875*). S. pneumoniae WT (strain R3584) was grown until early log phase ($OD_{550}$=0.1) and incubated with or without 1.2 µM RumC1 for 10 min before RNA extraction and RT-qPCR. The relative expression of each gene was standardized with the expression of *rpoD*. For each gene, relative expression versus the wild-type strain is indicated. Standard deviations from two independent replicates are indicated.

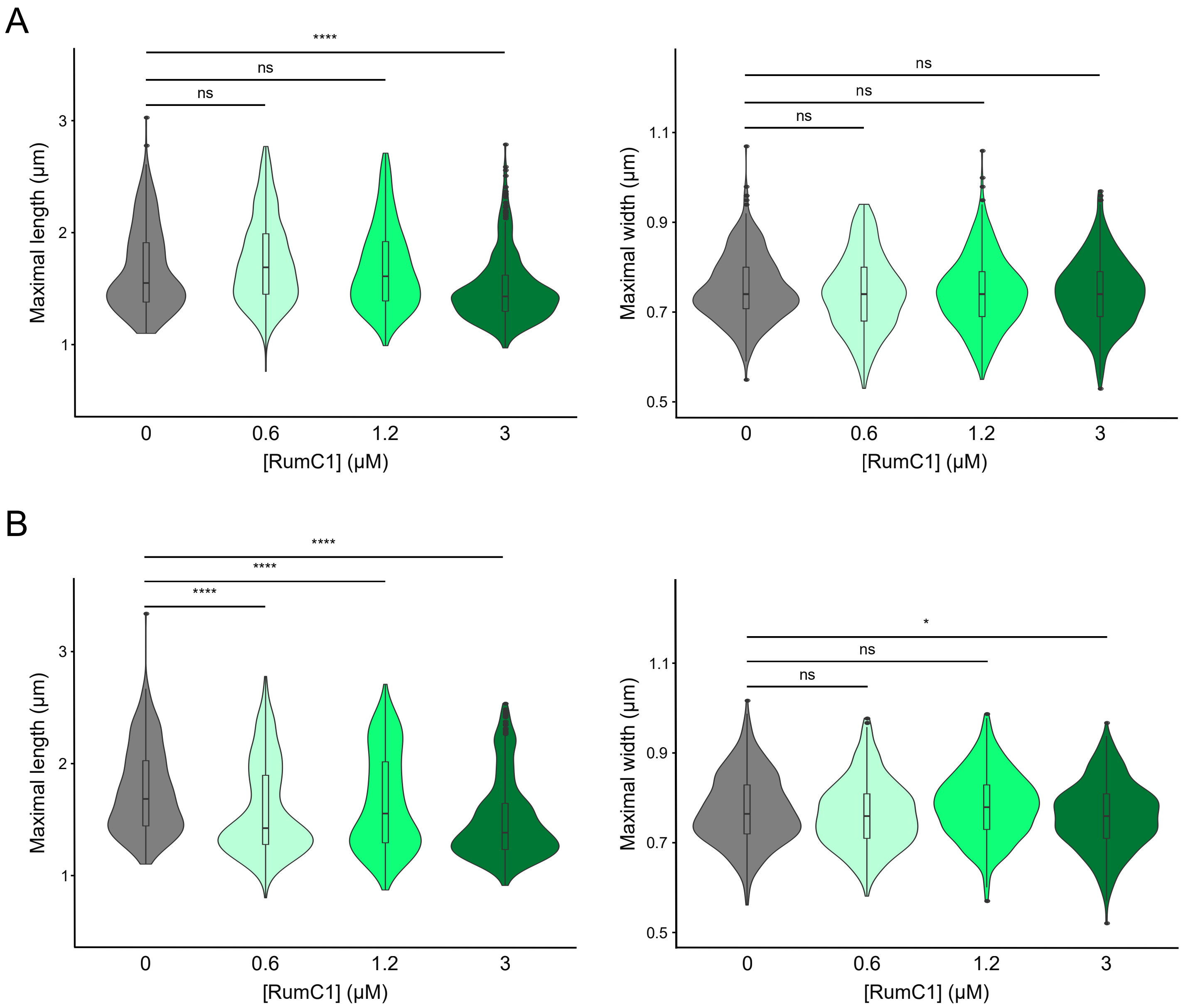
A
****
ns
ns
Maximal length (µm)
3
2
1
0
0.6
1.2
3
[RumC1] (µM)
ns
ns
ns
Maximal width (µm)
1.1
0.9
0.7
0.5
B
****
****
****
*

Figure S4

**Figure S4: Effect of RumC1 on the length and width of *S. pneumoniae* cells.**

(A) Violin plots representing the distribution of the maximal length (left panel) and the maximal width (right panel) of WT cells (R1501) untreated (n=424) or treated with 0.6 (n=407), 1.2 (n=410) or 3 µM (n=408) RumC1 for 10 min at 37°C. (B) Violin plots representing the distribution of the maximal length (left panel) and the maximal width (right panel) of WT cells (R1501) untreated (n=408) or treated with 0.6 (n=407), 1.2 (n=405) or 3 µM (n=405) RumC1 for 30 min at 37°C. Cell length and width measured during the experiments presented in Fig 2. Data obtained from two independent experiments. Boxes extend from the 25$^{th}$ percentile to the 75$^{th}$ percentile, with the horizontal line at the median. Dots represent outliers. Statistical analysis was performed using the U test of Mann-Whitney (n.s., non-significant, p-value > 0,05; *, p-value < 0,05; **, p-value < 0,01; ****, p-value < 0,0001).

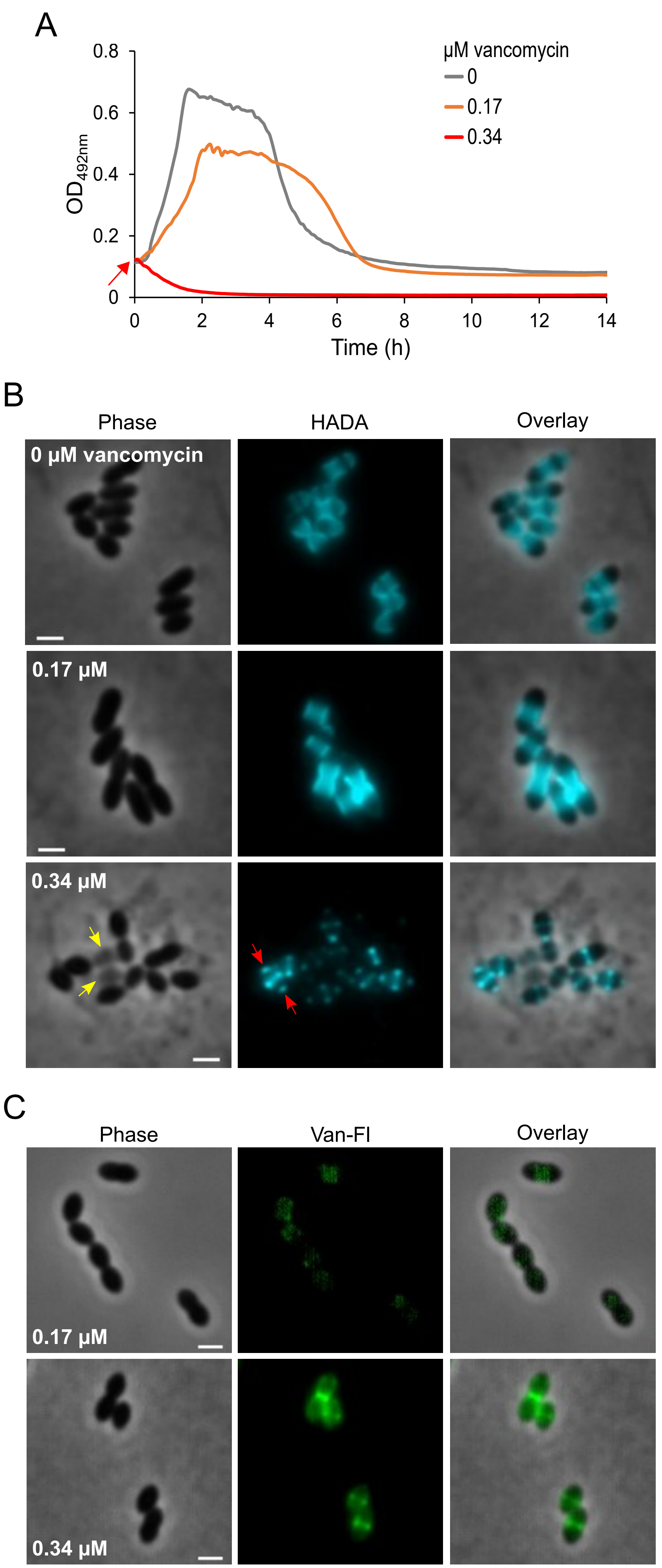
A
0.8
0.6
0.4
0.2
0
$OD_{492nm}$
µM vancomycin
0
0.17
0.34
0
2
4
6
8
10
12
14
Time (h)
B
Phase
HADA
Overlay
0 µM vancomycin
0.17 µM
0.34 µM
C
Phase
Van-Fl
Overlay
0.17 µM
0.34 µM

Figure S5

**Figure S5: Effect of vancomycin on growth, cell morphology and PG synthesis in *S. pneumoniae*.**

(A) Effect of vancomycin on growth of high-cell-density cultures of a wild-type strain of *S. pneumoniae* (R1501). Two concentrations of vancomycin were added at $OD_{550}$=0.1 (red arrow). (B) Morphology and HADA incorporation in *S. pneumoniae* WT (R1501) untreated or treated for 30 min with 0.17 or 0.34 µM vancomycin. Phase contrast, fluorescent (false-colored in light blue), and false-colored overlay images are shown. The yellow arrows indicate the “ghost cells”. The red arrows indicate an example of cells with HADA foci at both cell poles. Scale bar, 1 µm. Images are representative of two independent replicates. (C) Binding of vancomycin to pneumococcal cells. Exponentially growing cells of a wild-type strain of *S. pneumoniae* (R1501) were labelled for 10 min with 0.17 or 0.34 µM BODIPY® FL vancomycin (Van-Fl). Phase contrast, fluorescent, and overlay images are shown. Scale bar, 1 µm. Images are representative of two independent replicates.

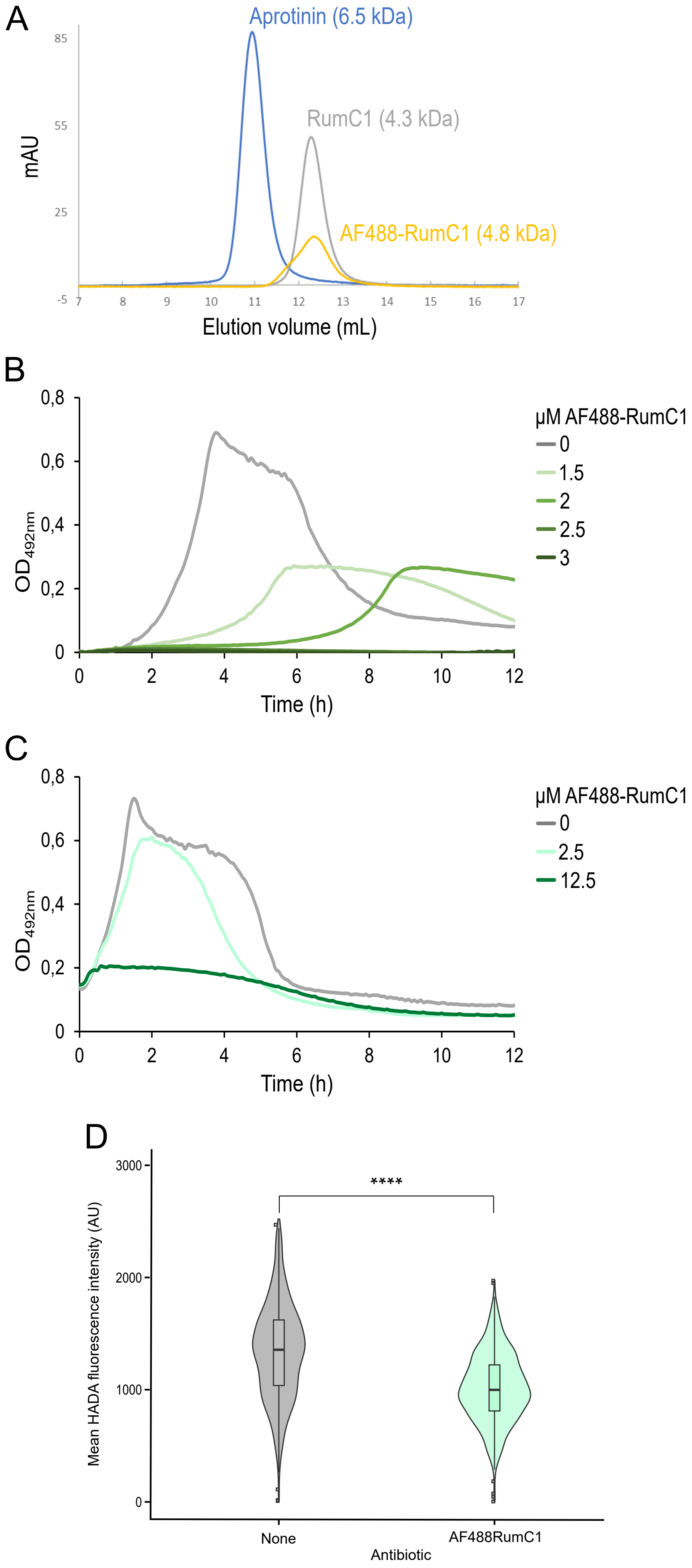
A
Aprotinin (6.5 kDa)
RumC1 (4.3 kDa)
AF488-RumC1 (4.8 kDa)
mAU
85
55
25
-5
Elution volume (mL)
B
0,8
0,6
0,4
0,2
0
$OD_{492nm}$
0
2
4
6
8
10
12
Time (h)
µM AF488-RumC1
0
1.5
2
2.5
3
C
0,8
0,6
0,4
0,2
0
$OD_{492nm}$
0
2
4
6
8
10
12
Time (h)
µM AF488-RumC1
0
2.5
12.5
D
****
3000
2000
1000
0
Mean HADA fluorescence intensity (AU)
None
AF488RumC1
Antibiotic

Figure S6

**Figure S6: Properties and activity of AF488-RumC1.**

(A) Determination of the oligomerization state of RumC1 and AF488-RumC1. Gel filtration chromatogram profile of Aprotinin (blue), RumC1 (gray) and AF488-RumC1 (yellow). The indicated MW correspond to the monomeric form of the three protein samples. (B) Determination of the MIC of AF488-RumC1 in *S. pneumoniae* WT (strain R1501). (C) Effect of AF488-RumC1 on growth of high-cell-density cultures ($OD_{550}$=0.1) of a wild-type strain of *S. pneumoniae* (strain R1501). (D) Violin plots representing the mean HADA fluorescence intensity in WT (R1501) cells untreated (n=808) or treated with 2.5 µM AF488-RumC1 (n=821) for 10 min at 37°C (related to Fig 3B). Boxes extend from the 25$^{th}$ percentile to the 75$^{th}$ percentile, with the horizontal line at the median. Dots represent outliers. Statistical analysis was performed using the U test of Mann-Whitney (****, p-value < 0,0001).

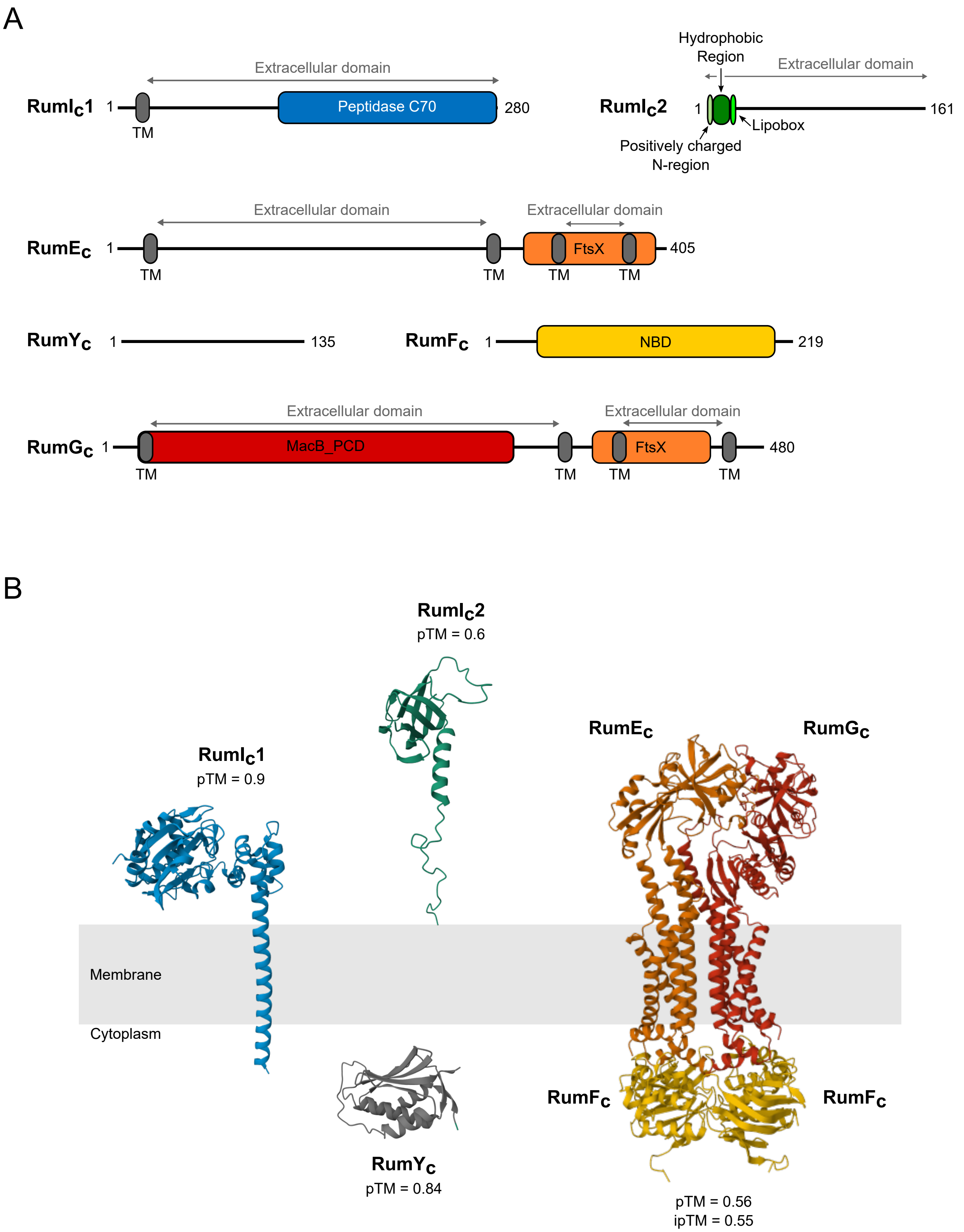
A
Extracellular domain
RumI_C1
1
Peptidase C70
280
TM
Hydrophobic Region
Extracellular domain
RumI_C2
1
161
Lipobox
Positively charged N-region
Extracellular domain
Extracellular domain
RumE_C
1
FtsX
405
TM
TM
TM
TM
RumY_C
1
135
RumF_C
1
NBD
219
Extracellular domain
Extracellular domain
RumG_C
1
MacB_PCD
FtsX
480
TM
TM
TM
TM
B
RumI_C2
pTM = 0.6
RumE_C
RumG_C
RumI_C1
pTM = 0.9
Membrane
Cytoplasm
RumF_C
RumF_C
RumY_C
pTM = 0.84
pTM = 0.56
ipTM = 0.55

Figure S7

**Figure S7: Domain and structure prediction of the immunity proteins.**

(A) Predicted domain organization and topology of $RumI_c1$, $RumI_c2$, $RumE_c$, $RumY_c$, $RumF_c$ and $RumG_c$. TM: TransMembrane segment, FtsX: FtsX-like permease family, NBD: Nucleotide Binding Domain, MacB_PCD: MacB-like Periplasmic Core Domain. (B) AlphaFold3 models of $RumI_c1$, $RumI_c2$, $RumY_c$ and $RumE_cF_cG_c$. The predicted structure of the mature form of the lipoprotein $RumI_c2$ is presented (the residue at position +1 is the cysteine of the lipobox (C21)). The predicted structure of the ABC transporter $RumE_cF_cG_c$ composed *of two copies of $RumF_c$,* one copy of $RumE_c$ and one copy of $RumG_c$ is presented. The predicted template modeling (pTM) scores of each prediction are indicated. The interface predicted template modeling (ipTM) score of the $RumE_cF_cG_c$ complex prediction is indicated.

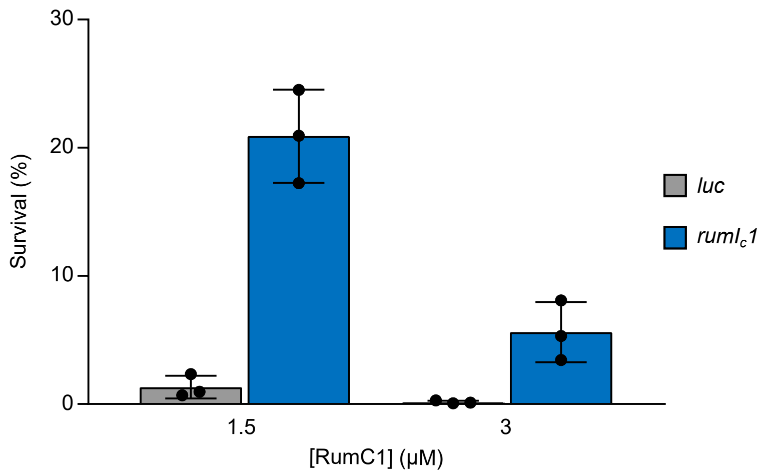

Figure S8

**Figure S8: RumI$_c$1 confers tolerance to RumC1 on pneumococcal cells.**

Survival of strains *CEPI-P$_{lac}$-luc* (R5339) and *CEPI-P$_{lac}$-rumI$_c$1* (R5231) exposed to RumC1. Early log phase cells were exposed to 0, 1.5 or 3 µM RumC1 for 1h before serial dilution and plating. Survival percentages are calculated relative to cells not exposed to RumC1. Precultures and cultures were performed in the presence of 50 µM IPTG. Individual data points, means and standard deviations obtained from three independent experiments are presented.

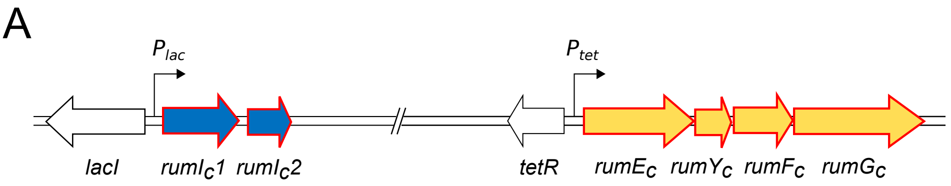

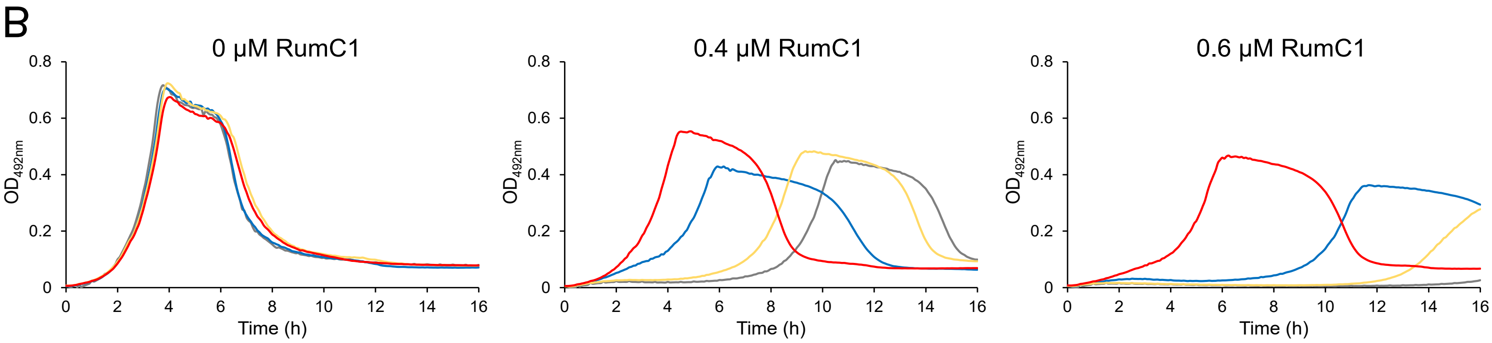

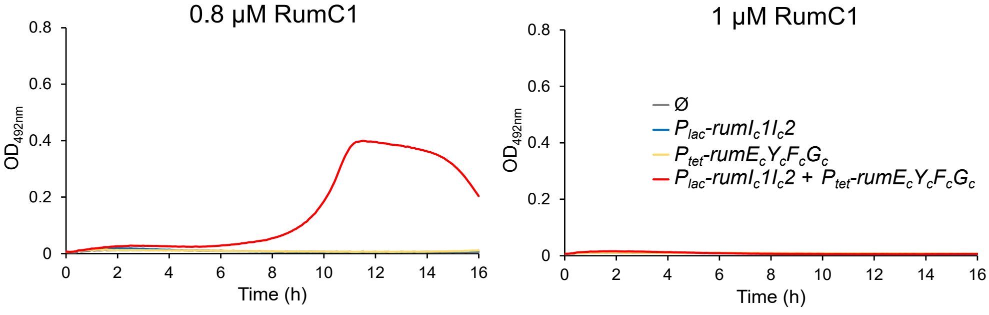

| Expressed genes | RumC1 MIC (µM) |
|---|---|
| Ø | 0.6 |
| $P_{lac}$-rumI$_c$1I$_c$2 | 0.8 |
| $P_{tet}$-rumE$_c$Y$_c$F$_c$G$_c$ | 0.8 |
| $P_{lac}$-rumI$_c$1I$_c$2 + $P_{tet}$-rumE$_c$Y$_c$F$_c$G$_c$ | 1 |

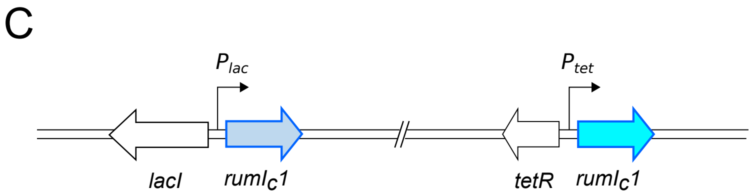

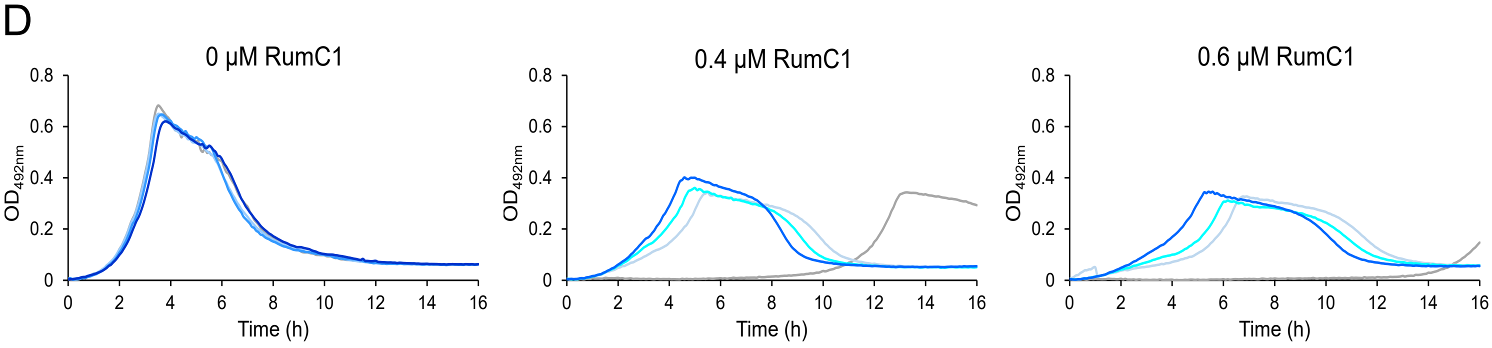

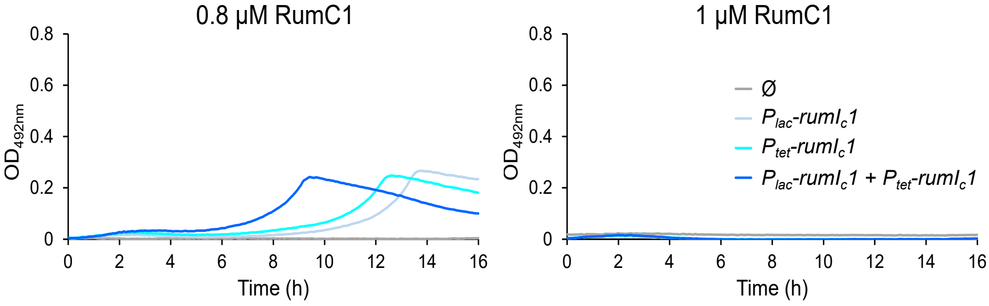

| Expressed genes | RumC1 MIC (µM) |
|---|---|
| Ø | 0.6 |
| $P_{lac}$-rumI$_c$1 | 1 |
| $P_{tet}$-rumI$_c$1 | 0.6 |
| $P_{lac}$-rumI$_c$1 + $P_{tet}$-rumI$_c$1 | 0.6 |

Figure S9

**Figure S9: Effect of expression of all RumC immunity genes and effect of increased expression of *rumI*$_c$*1* on resistance of pneumococcal cells to RumC1.**

(A) Genetic organization of *CEPI-P*$_{lac}$*-rumI*$_c$*1I*$_c$*2* and *CEPII-P*$_{tet}$*-rumE*$_C$*Y*$_C$*F*$_C$*G*$_C$ expression platforms. (B) Growth of strain *CEPI-P*$_{lac}$*-rumI*$_c$*1I*$_c$*2, CEPII-P*$_{tet}$*-rumE*$_C$*Y*$_C$*F*$_C$*G*$_C$ (R5177), in the absence or presence of various concentrations of RumC1. The RumC1 concentrations are indicated on top of the graphs. Precultures and cultures were performed in the absence of inductor (Ø) or in the presence of 50 µM IPTG, 100 ng $mL^{-1}$ ATC or 50 µM IPTG and 100 ng $mL^{-1}$ ATC to induce expression of *rumI*$_c$*1I*$_c$*2*, *rumE*$_C$*Y*$_C$*F*$_C$*G*$_C$ or *rumI*$_c$*1I*$_c$*2* and *rumE*$_C$*Y*$_C$*F*$_C$*G*$_C$, respectively. The corresponding RumC1 MICs of each condition are listed in the table in the right. Data shown here are representative of two independent replicates. (C) Genetic organization of *CEPI-P*$_{lac}$*-rumI*$_c$*1* and *CEPII-P*$_{tet}$*-rumI*$_c$*1* expression platforms. (D) Growth of strain *CEPI-P*$_{lac}$*-rumI*$_c$*1, CEPII-P*$_{tet}$*-rumI*$_c$*1* (R5232), in the absence or in the presence of various concentrations of RumC1. Precultures and cultures were performed in the absence of inductor (Ø) or in the presence of 50 µM IPTG, 100 ng $mL^{-1}$ ATC or 50 µM IPTG and 100 ng $mL^{-1}$ ATC to induce expression of *rumI*$_c$*1* under the control of the *lac* promoter, the *tet* promoter or both promoters, respectively. The RumC1 concentrations are indicated on top of the graphs. The corresponding RumC1 MICs of each condition are listed in the table on the right. Data shown here are representative of two independent experiments.

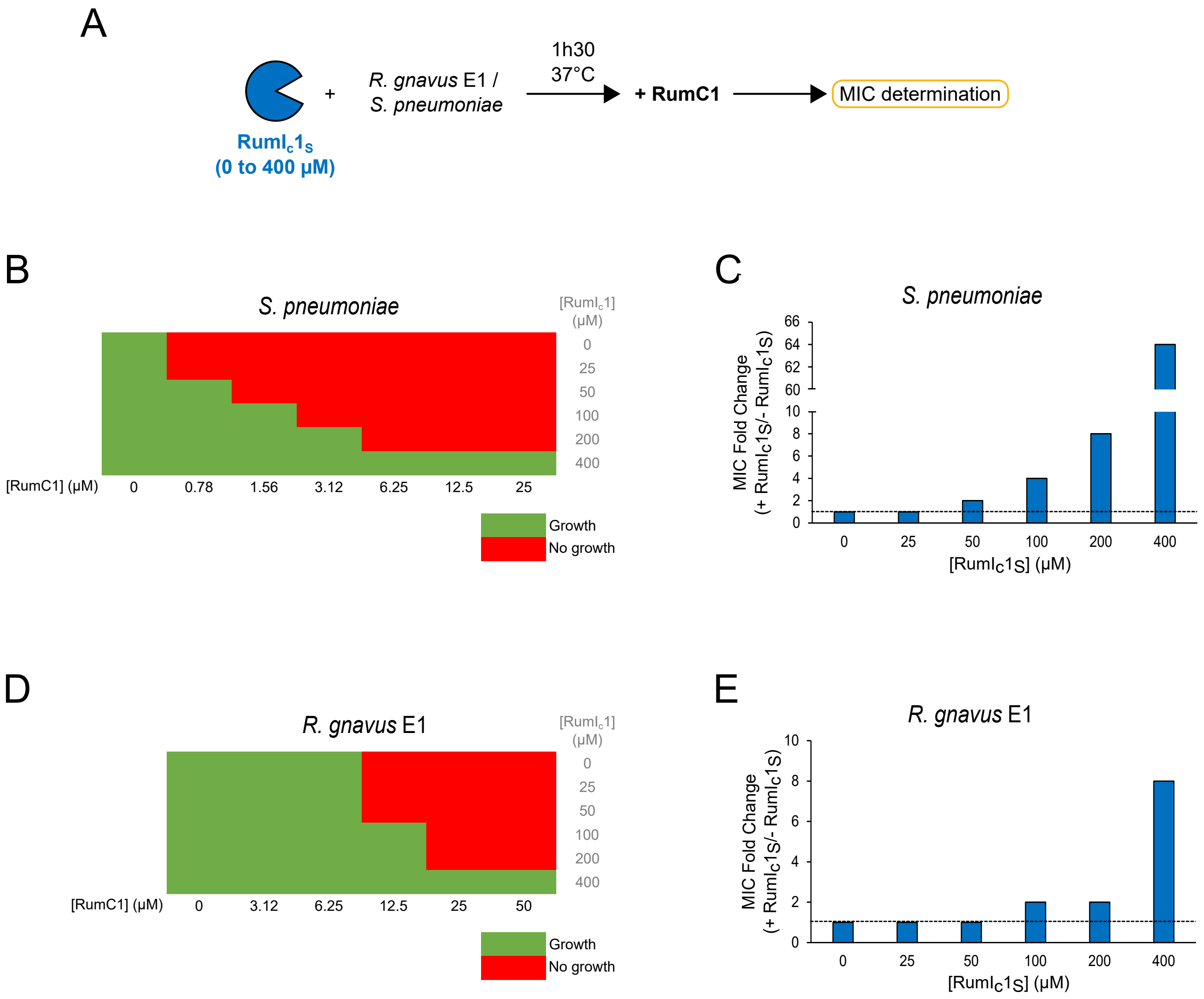
A
+
R. gnavus E1 /
S. pneumoniae
1h30
37°C
+ RumC1
MIC determination
RumIc1S
(0 to 400 µM)
B
S. pneumoniae
[RumIc1] (µM)
0
25
50
100
200
400
[RumC1] (µM)
0
0.78
1.56
3.12
6.25
12.5
25
Growth
No growth
C
S. pneumoniae
MIC Fold Change (+ RumIc1S/- RumIc1S)
66
64
62
60
10
8
6
4
2
0
0
25
50
100
200
400
[RumIc1S] (µM)
D
R. gnavus E1
[RumIc1] (µM)
0
25
50
100
200
400
[RumC1] (µM)
0
3.12
6.25
12.5
25
50
Growth
No growth
E
R. gnavus E1
MIC Fold Change (+ RumIc1S/- RumIc1S)
10
8
6
4
2
0
0
25
50
100
200
400
[RumIc1S] (µM)

Figure S10

**Figure S10: Addition of the soluble domain of RumI$_c$1 to the extracellular medium protects *S. pneumoniae* and *R. gnavus* E1 cells from RumC1.**

(A) Schematic representation of experiment carried out to assess RumI$_c$1 can confer immunity to RumC1 in *R. gnavus* E1 and *S. pneumoniae* when added in the extracellular medium. *R. gnavus E1* and *S. pneumoniae* were treated with increasing concentrations of the His6-tagged purified extracytoplasmic domain of RumI$_c$1 (RumI$_c$1$_S$) for 1h30 at 37°C. Cells were then exposed to increasing concentrations of RumC1 and the RumC1 MIC for each condition was determined. Growth of (B) *S. pneumoniae* (strain R1501) and (D) *R. gnavus* E1 cells pre-treated with different concentrations of RumI$_c$1$_S$ in presence of increasing concentrations of RumC1. Protocol carried out as in panel (A). Growth is indicated by green whereas growth inhibition is indicated in red. (C) and (E) The relative MIC values obtained with RumI$_c$1$_S$ compared to those obtained without RumI$_c$1$_S$ (0.625 µM for *S. pneumoniae* and 12.5 µM for *R. gnavus* E1) determined from panels (B) and (D) are shown.

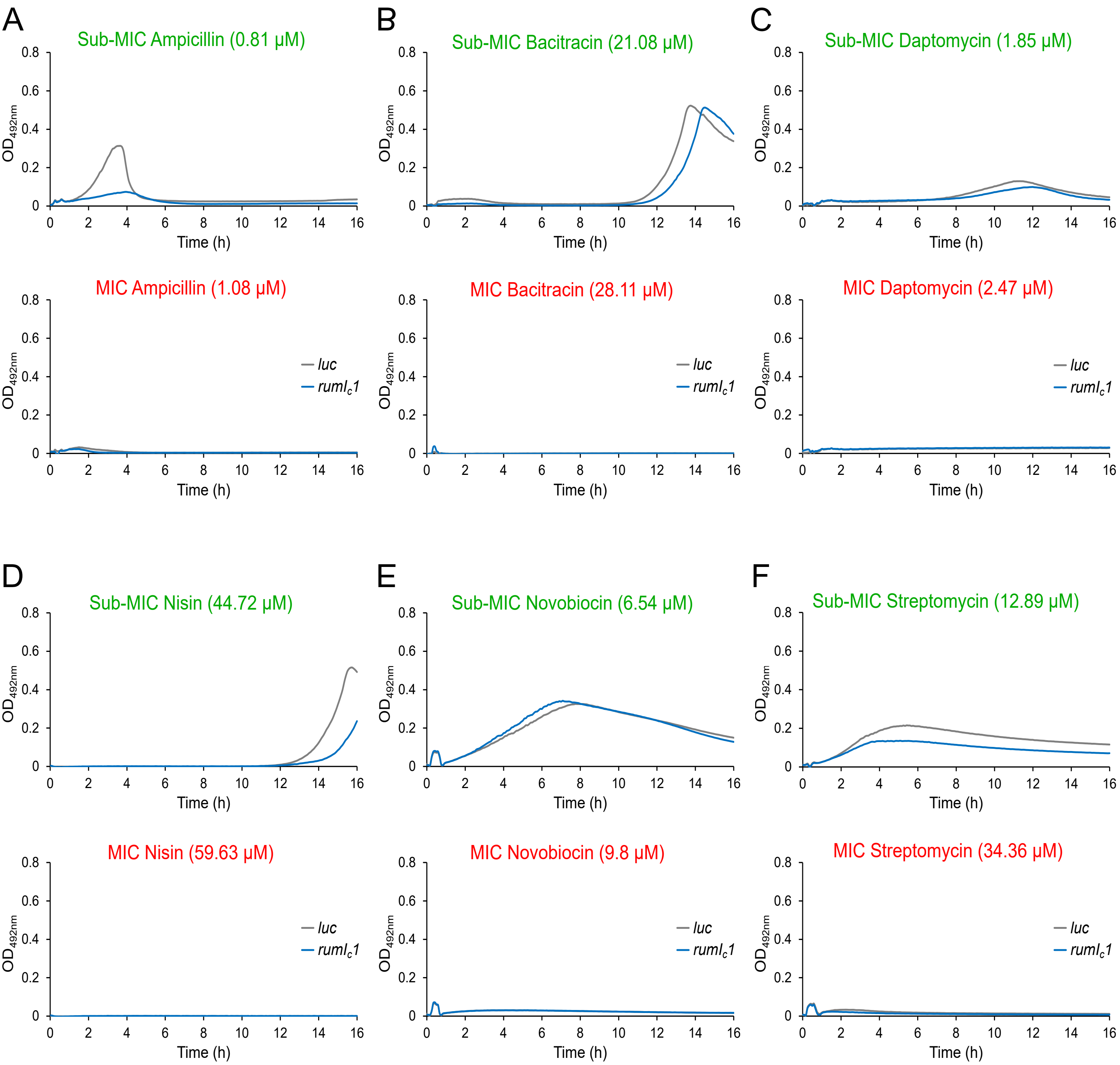
A
Sub-MIC Ampicillin (0.81 μM)
MIC Ampicillin (1.08 μM)
B
Sub-MIC Bacitracin (21.08 μM)
MIC Bacitracin (28.11 μM)
C
Sub-MIC Daptomycin (1.85 μM)
MIC Daptomycin (2.47 μM)
D
Sub-MIC Nisin (44.72 μM)
MIC Nisin (59.63 μM)
E
Sub-MIC Novobiocin (6.54 μM)
MIC Novobiocin (9.8 μM)
F
Sub-MIC Streptomycin (12.89 μM)
MIC Streptomycin (34.36 μM)
OD492nm
Time (h)
luc
rumlc1
0
0.2
0.4
0.6
0.8
0
2
4
6
8
10
12
14
16

Figure S11

**Figure S11: The immunity protein RumI$_c$1 does not protect pneumococcal cells against Ampicillin, Bacitracin, Daptomycin, Nisin, Novobiocin and Streptomycin.**

Growth of strains *CEPI-P$_{lac}$-luc* (R5339) and *CEPI-P$_{lac}$-rumI$_c$1* (R5231) in the presence of sub-MIC (upper panels) and MIC (lower panels) concentrations of (A) Ampicillin (sub-MIC: 0.81 µM, MIC: 1.08 µM) (B) Bacitracin (sub-MIC: 21.08 µM, MIC: 28.11 µM), (C) Daptomycin (sub-MIC: 1.85 µM, MIC: 2.47 µM), (D) Nisin (sub-MIC: 44.72 µM, MIC: 59.63 µM), (E) Novobiocin (sub-MIC: 6.54 µM, MIC: 9.8 µM) and (F) Streptomycin (sub-MIC: 12.89 µM, MIC: 34.36 µM). Precultures and cultures were performed in the presence of 50 µM IPTG. Data shown here are representative of two or three independent replicates.

A

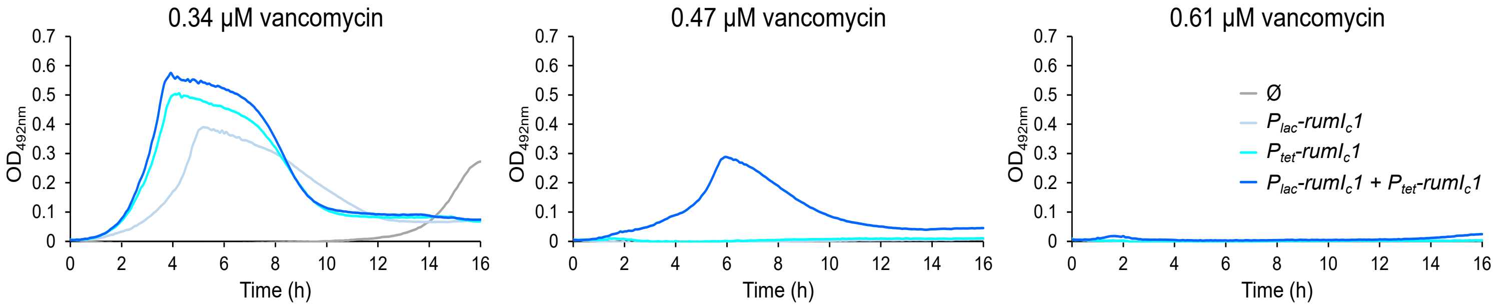

B

| **Expressed genes** | **Vancomycin MIC (µM)** |
|---|---|
| Ø | *0.34* |
| *$P_{lac}$-rumI$_c$1* | *0.47* |
| *$P_{tet}$-rumI$_c$1* | *0.47* |
| *$P_{lac}$-rumI$_c$1* + *$P_{tet}$-rumI$_c$1* | *0.61* |

Figure S12

**Figure S12: Effect of increased expression of *rumI$_c$1* on vancomycin resistance in pneumococcal cells.**

(A) Growth of strain *CEPI-P$_{lac}$-rumI$_c$1, CEPII-P$_{tet}$-rumI$_c$1* (R5232), in the presence of various concentrations of vancomycin. Precultures and cultures were performed in the absence of inductor (Ø) or in the presence of 50 µM IPTG, 100 ng mL$^{-1}$ ATC or 50 µM IPTG and 100 ng mL$^{-1}$ ATC to induce expression of *rumI$_c$1* under the control of the *lac* promoter, the *tet* promoter or both promoters, respectively. The vancomycin concentrations are indicated on top of the graphs. Data shown here are representative of two independent experiments. (B) Corresponding vancomycin MICs determined from (A).

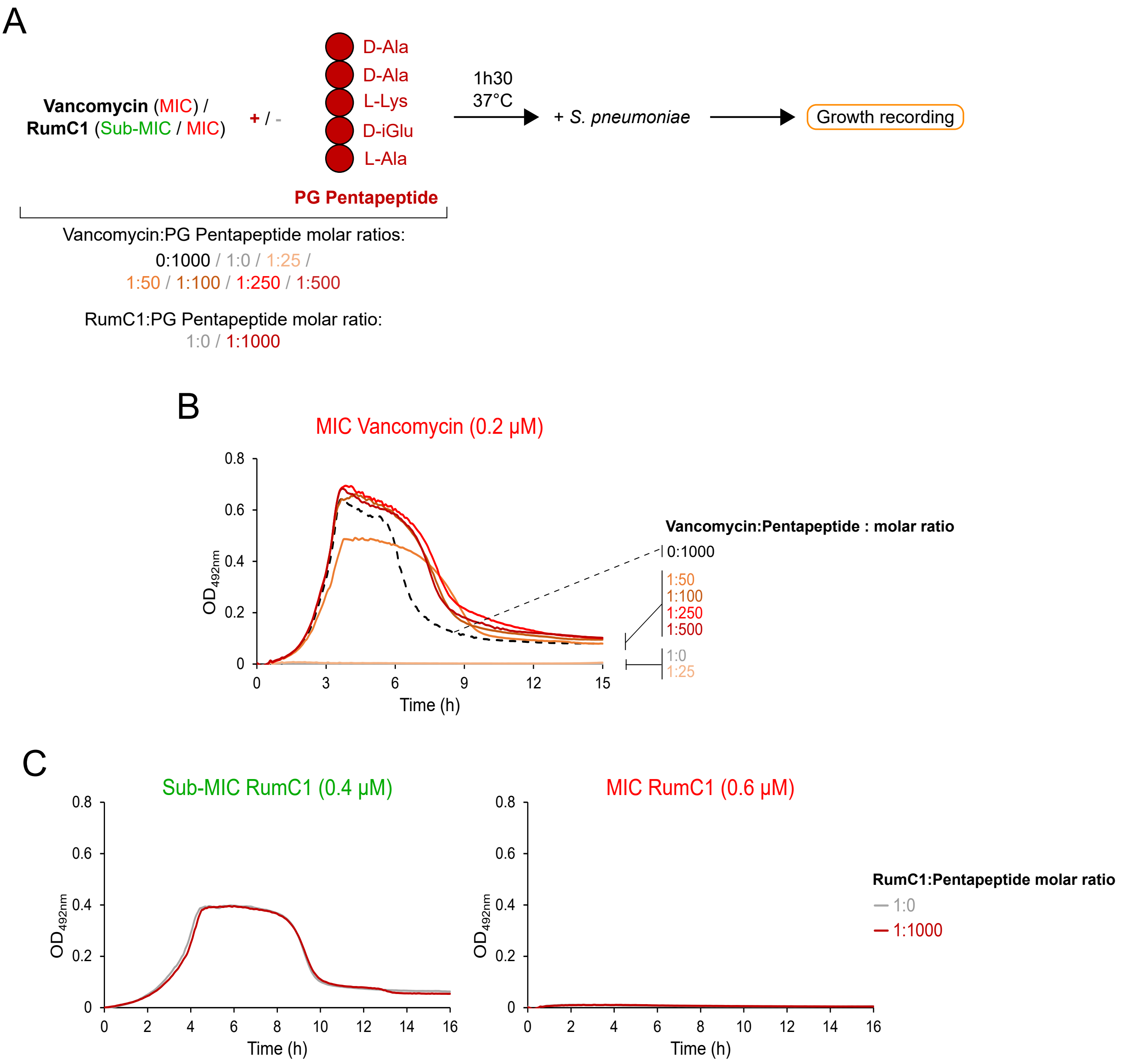

A
Vancomycin (MIC) /
RumC1 (Sub-MIC / MIC)
+ / -
D-Ala
D-Ala
L-Lys
D-iGlu
L-Ala
PG Pentapeptide
1h30
37°C
+ S. pneumoniae
Growth recording
Vancomycin:PG Pentapeptide molar ratios:
0:1000 / 1:0 / 1:25 /
1:50 / 1:100 / 1:250 / 1:500
RumC1:PG Pentapeptide molar ratio:
1:0 / 1:1000
B
MIC Vancomycin (0.2 μM)
OD492nm
Time (h)
Vancomycin:Pentapeptide : molar ratio
0:1000
1:50
1:100
1:250
1:500
1:0
1:25
C
Sub-MIC RumC1 (0.4 μM)
MIC RumC1 (0.6 μM)
OD492nm
Time (h)
RumC1:Pentapeptide molar ratio
1:0
1:1000

Figure S13

**Figure S13: The PG Pentapeptide antagonizes vancomycin but not RumC1.**

(A) Schematic representation of experiment carried out to assess whether vancomycin and RumC1 are antagonized by the chemically synthesized PG Pentapeptide of *S. pneumoniae*. Concentrated solutions of vancomycin and RumC1 were incubated or not with the PG Pentapeptide for 1h30 at 37°C before being added to a cell suspension of *S. pneumoniae* WT (strain R1501). The vancomycin solution used is concentrated to give a final concentration equal to its MIC (0.2 µM) when added to *S. pneumoniae* cultures. The RumC1 solutions used are concentrated to give final concentration below (0.4 µM) or equal (0.6 µM) to its MIC. Cells were then incubated for 16h at 37°C and cell growth was recorded throughout the incubation. (B) Growth of WT strain (R1501) in the presence of a MIC concentration of vancomycin (0.2 µM) pre-incubated with the chemically synthesized PG Pentapeptide at different vancomycin:PG Pentapeptide molar ratios as described in panel (A). A control of cell growth in the absence of vancomycin and in the presence of the maximum concentration of PG Pentapeptide (vancomycin:PG Pentapeptide molar ratio of 0:1000) is represented by the dashed black curve. Data shown here are representative of two independent experiments. (C) Growth of WT strain (R1501) in the presence of Sub-MIC (0.4 µM) or MIC (0.6 µM) concentrations of RumC1 pre-incubated with the PG Pentapeptide at a RumC1:PG Pentapeptide molar ratio of 1:0 and 1:1000 as described in panel (A). Data shown here are representative of three independent experiments.

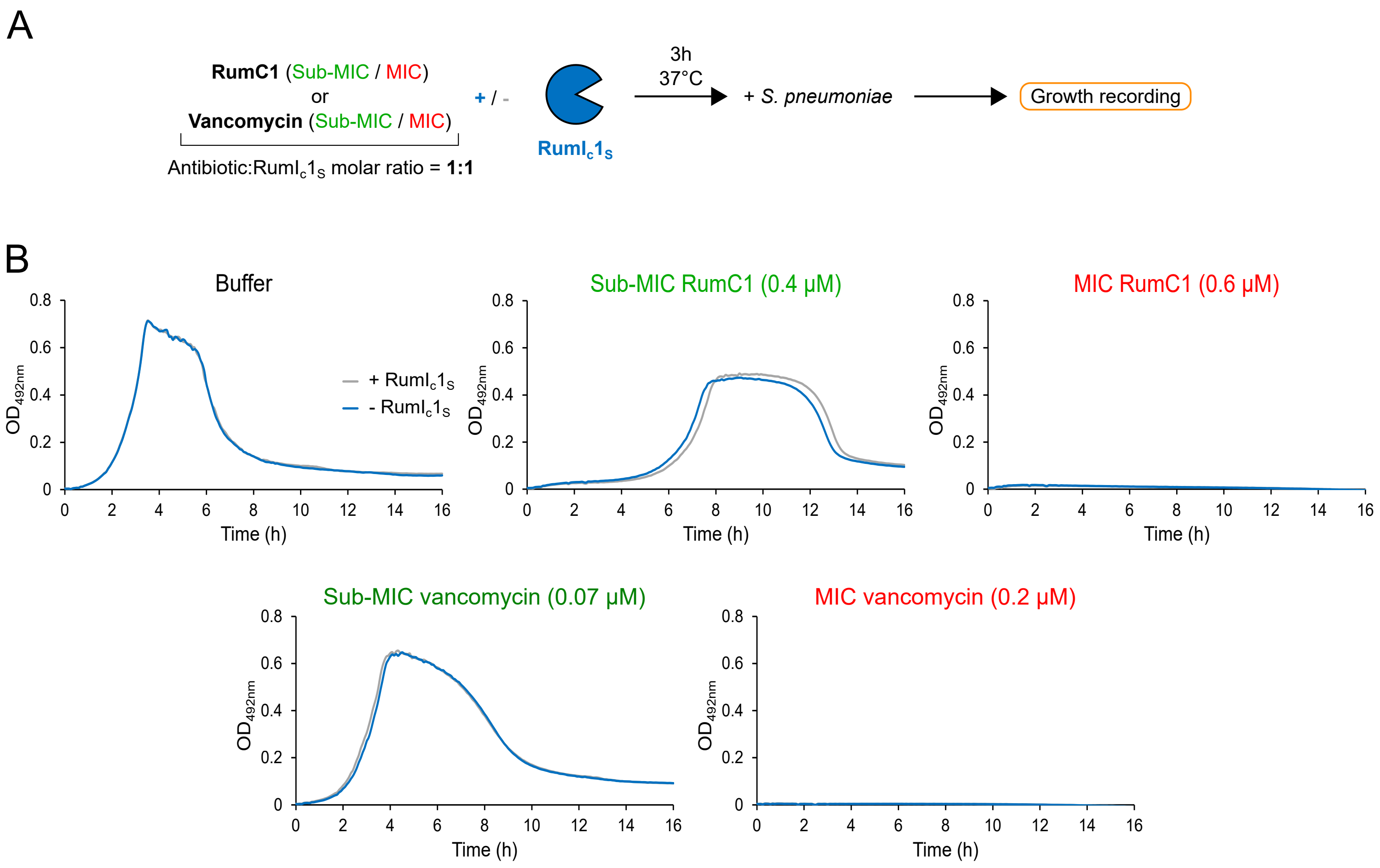
A
RumC1 (Sub-MIC / MIC)
or
Vancomycin (Sub-MIC / MIC)
Antibiotic:$RumI_c1_S$ molar ratio = 1:1
+ / -
$RumI_c1_S$
3h
37°C
+ S. pneumoniae
Growth recording
B
Buffer
Sub-MIC RumC1 (0.4 µM)
MIC RumC1 (0.6 µM)
Sub-MIC vancomycin (0.07 µM)
MIC vancomycin (0.2 µM)
+ $RumI_c1_S$
- $RumI_c1_S$
$OD_{492nm}$
Time (h)
0
0.2
0.4
0.6
0.8
0
2
4
6
8
10
12
14
16

Figure S14

**Figure S14: RumI$_c$1 does not inactivate RumC1 or vancomycin.**

(A) Schematic representation of experiment carried out to assess whether pre-treatment with RumI$_c$1 reduces the toxicity of RumC1 and vancomycin to pneumococcal cells. Concentrated solutions of RumC1 and vancomycin were treated or not with the His6-tagged purified extracytoplasmic domain of RumI$_c$1 (RumI$_c$1$_S^{WT}$) for 3h at 37°C before being added to a cell suspension of *S. pneumoniae* WT (strain R1501). The RumC1 and vancomycin solutions used are concentrated to give final concentrations below (0.4 µM RumC1; 0.07 µM vancomycin) or equal (0.6 µM RumC1; 0.2 µM Vancomycin) to their MIC when added to *S. pneumoniae* cultures. Cells were then incubated for 16h at 37°C and cell growth was recorded throughout the incubation. (B) Growth of the wild-type strain (R1501) of *S. pneumoniae* in the presence of buffer or in the presence of sub-MIC (0.4 µM RumC1; 0.0.7 µM vancomycin) or MIC (0.6 µM RumC1; 0.2 µM vancomycin) concentrations of RumC1 and vancomycin pre-treated (blue curves) or not (grey curves) with the purified extracellular domain of RumI$_c$1 as described in panel (A). Data shown here are representative of two or three independent replicates.

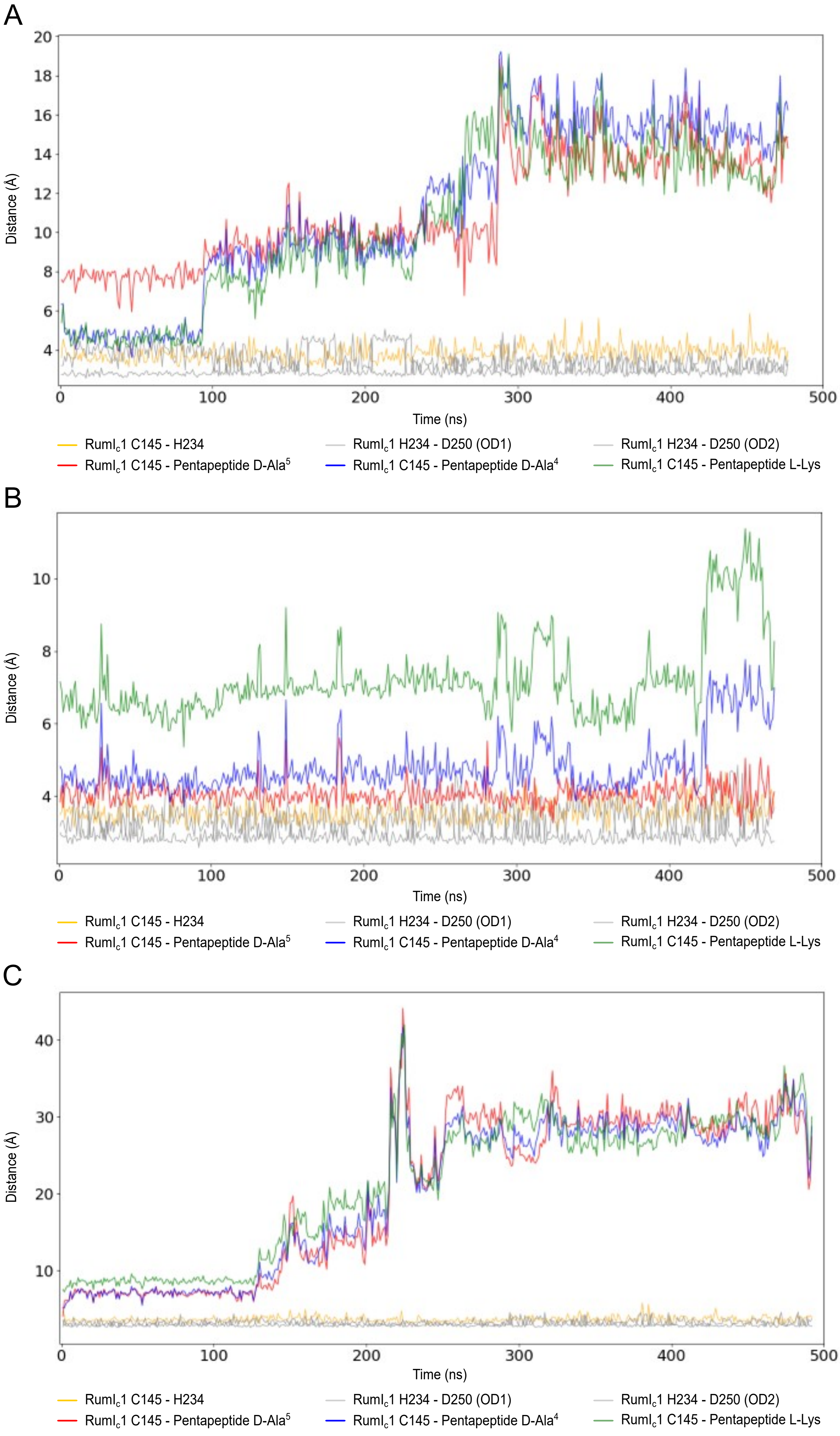
A
Distance (Å)
Time (ns)
RumI$_c$1 C145 - H234
RumI$_c$1 H234 - D250 (OD1)
RumI$_c$1 H234 - D250 (OD2)
RumI$_c$1 C145 - Pentapeptide D-Ala$^5$
RumI$_c$1 C145 - Pentapeptide D-Ala$^4$
RumI$_c$1 C145 - Pentapeptide L-Lys
B
Distance (Å)
Time (ns)
RumI$_c$1 C145 - H234
RumI$_c$1 H234 - D250 (OD1)
RumI$_c$1 H234 - D250 (OD2)
RumI$_c$1 C145 - Pentapeptide D-Ala$^5$
RumI$_c$1 C145 - Pentapeptide D-Ala$^4$
RumI$_c$1 C145 - Pentapeptide L-Lys
C
Distance (Å)
Time (ns)
RumI$_c$1 C145 - H234
RumI$_c$1 H234 - D250 (OD1)
RumI$_c$1 H234 - D250 (OD2)
RumI$_c$1 C145 - Pentapeptide D-Ala$^5$
RumI$_c$1 C145 - Pentapeptide D-Ala$^4$
RumI$_c$1 C145 - Pentapeptide L-Lys

Figure S15

**Figure S15: Catalytic distance dynamics across independent MD replicas.**

(A) Second replica initiated from the lowest-energy minimum identified in the PELE energy landscape (the first replica is shown in Figure 6F). (B,C) Two independent replicas initiated from the second-lowest energy minimum identified in the PELE simulations. In all cases, the same catalytic distances described in Figure 6F were monitored to assess the stability of the catalytic triad and the positioning of the substrate relative to Cys145 over time.

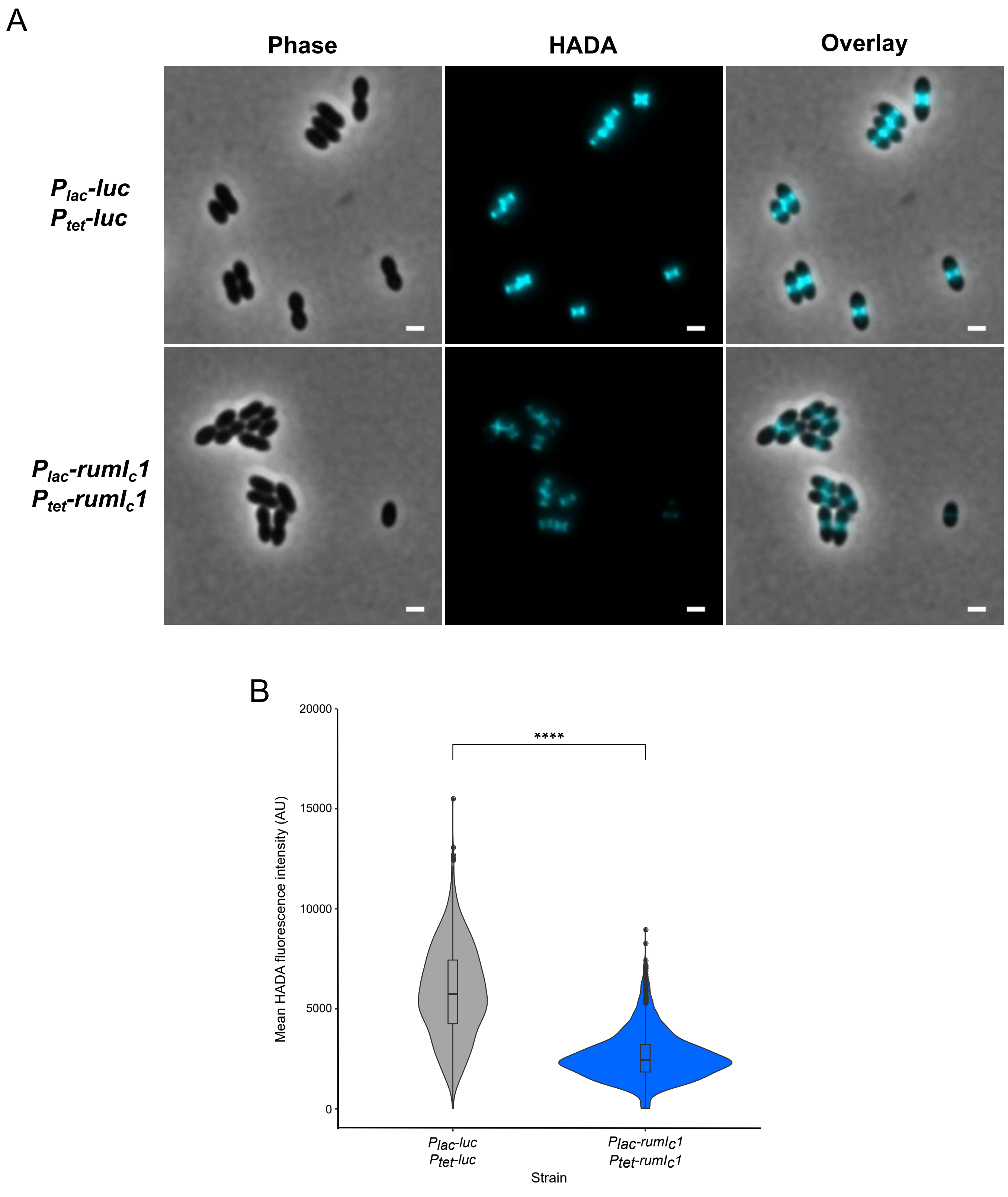
A
Phase
HADA
Overlay
Plac-luc
Ptet-luc
Plac-rumlc1
Ptet-rumlc1
B
20000
15000
10000
5000
0
****
Mean HADA fluorescence intensity (AU)
Plac-luc
Ptet-luc
Plac-rumlc1
Ptet-rumlc1
Strain

Figure S16

**Figure S16: RumI$_c$1 reduces transpeptidase activity.**

(A) HADA incorporation in cells expressing *luc* (*CEPI-P$_{lac}$-luc, CEPII-P$_{tet}$-luc,* strain R5172) or *rumI$_c$1* (*CEPI-P$_{lac}$-rumI$_c$1, CEPII-P$_{tet}$- rumI$_c$1,* strain R5232) from the P$_{lac}$ and P$_{tet}$ promoters. Phase contrast, fluorescent (false-colored in light blue), and false-colored overlay images are shown. Cells were incubated for 10 min with HADA. Precultures and cultures were performed in the presence of 50 µM IPTG and 100 ng mL$^{-1}$ ATC. Scale bar, 1 µm. Images are representative of three independent replicates. (B) Violin plots representing mean HADA fluorescence intensity in cells expressing *luc* (*CEPI-P$_{lac}$-luc, CEPII-P$_{tet}$-luc,* strain R5172) (grey, n=1205) and in cells expressing *rumI$_c$1* (*CEPI-P$_{lac}$-rumI$_c$1, CEPII-P$_{tet}$- rumI$_c$1,* strain R5232) (blue, n=1119). Boxes extend from the 25$^{th}$ percentile to the 75$^{th}$ percentile, with the horizontal line at the median. Dots represent outliers. Statistical analysis was performed using the U test of Mann-Whitney (****, p-value < 0.0001).

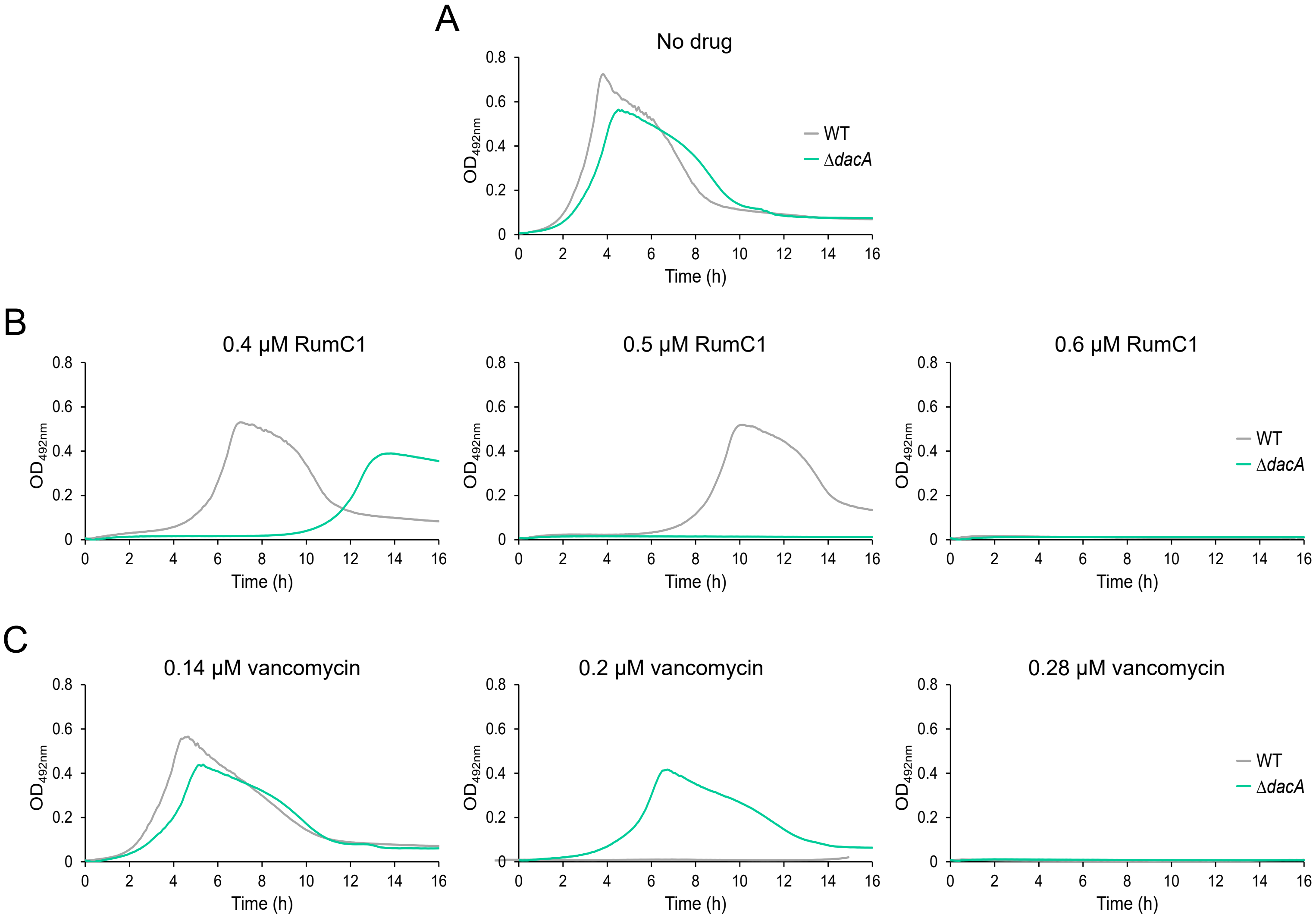

D

| Strain | RumC1 MIC (µM) | Vancomycin MIC (µM) |
|---|---|---|
| *WT* | 0.2 | 0.28 |
| *ΔdacA* | 0.5 | 0.4 |

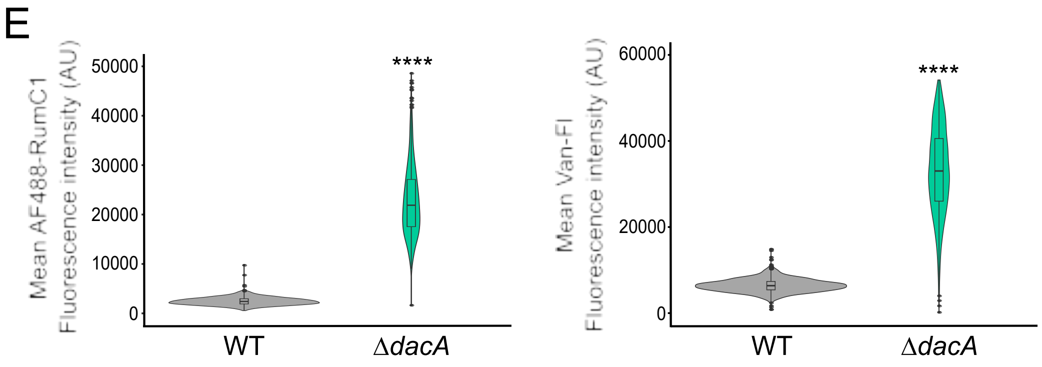

Figure S17

**Figure S17: The DD-Carboxypeptidase DacA has opposite effects on resistance to RumC1 and vancomycin.**

Full RumC1 and vancomycin resistance profiles of the WT and *ΔdacA* strains related to Fig. 7E. (A) Growth of wild-type (R1501) and Δ*dacA* (R5207) strains in the absence of antibiotics. Data shown here are representative of three independent experiments. Growth of wild-type (R1501) and Δ*dacA* (R5207) strains in the presence of increasing concentrations of (B) RumC1 and (C) vancomycin. Data shown here are representative of three independent experiments. (D) Corresponding RumC1 and vancomycin MICs determined from (B) and (C). (E) Quantification of the AF488-RumC1 and BODIPY® FL vancomycin (Van-Fl) fluorescence intensity in WT and *ΔdacA* cells related to Fig. 7F. Left panel: violin plots representing the mean AF488-RumC1 fluorescence intensity in WT (R1501) (grey, n=1524) and *ΔdacA* (R5207) (green, n=1291) cells. Right panel: violin plots representing the mean BODIPY® FL vancomycin (Van-Fl) fluorescence intensity in WT (R1501) (grey, n=1234) and *ΔdacA* (R5207) (green, n=977) cells. Boxes extend from the 25$^{th}$ percentile to the 75$^{th}$ percentile, with the horizontal line at the median. Dots represent outliers. Statistical analysis was performed using the U test of Mann-Whitney (****, p-value < 0.0001).
